# Training constrains neural routes to knowledge assembly

**DOI:** 10.64898/2026.03.13.711547

**Authors:** Q Wang, C French, P Bansiya, N Rabii, S Nelli

**Affiliations:** Department of Cognitive Science, Occidental College, Los Angeles, CA 90041, USA

## Abstract

A hallmark of human intelligence is the ability to rapidly restructure existing knowledge when new information reveals unexpected connections, a capacity termed knowledge assembly. This cognitive flexibility distinguishes human learning from artificial systems, which catastrophically forget when acquiring new relationships. Understanding the mechanisms underlying flexible knowledge reorganization thus has implications for continual learning in both humans and algorithms. Here, using electroencephalography we show that successful knowledge assembly depends on temporally orchestrated reactivation of prior neural representations. Critically, training schedules bias learners toward different representational strategies: blocked training promotes compressed certainty-weighted codes, while interleaved training yields high-dimensional factorized representations. Vanilla recurrent networks failed to develop human-like certainty geometries despite identical training, revealing missing computational principles in current artificial systems. These findings demonstrate that cognitive flexibility emerges through creative reuse of learned representations, with training history constraining available neural routes to reorganization.

## Introduction

When we discover that two previously separate social circles share a mutual acquaintance, we instantly reconfigure our understanding of both groups. This capacity for few-shot reorganization of relational knowledge, termed knowledge assembly, is a cornerstone of abstract reasoning and flexible cognition. Unlike artificial systems that catastrophically forget when acquiring new relationships, humans excel at integrating novel connections while preserving established knowledge. Understanding the neural mechanisms enabling this balance between stability and flexibility has profound implications for educational design, artificial intelligence development, and theories of adaptive cognition.

The neural substrates of flexible knowledge reorganization remain contentious. Classical frameworks emphasize medial temporal structures as primary sites for rapid relational learning^1–5^, with cortical representations emerging only through gradual consolidation^6–8^. Yet recent evidence demonstrates that knowledge assembly depends on global reorganization of frontoparietal representations^9^, suggesting cortical mechanisms drive rapid structure learning and online reconfiguration^9–15^. This raises fundamental questions regarding how cortical networks maintain stability while permitting flexible reorganization and what distinguishes cortical from hippocampal contributions to learning.

Standard connectionist models typically fail at knowledge assembly, exhibiting catastrophic interference when learning new relations^16^. One class of solutions emphasizes selective stabilization of critical synapses, mirroring biological findings on differential plasticity^17–20^. Another approach minimizes interference through high dimensional sparse coding strategies including orthogonalization and context-dependent gating^14,21–27^. Recent work synthesized these accounts, demonstrating that relational certainty encoded alongside task structure stabilizes critical relations while guiding representational updates during assembly^9^. This framework proposes that curved manifolds naturally encode comparison difficulty, contributing to documented positional effects in behavior^28–31^. Such U-shaped geometries may arise from concurrent rate coding and rank selectivity in cortical pyramidal neurons^32,33^. However, while prior work showed neural networks encoding within-context certainty can assemble knowledge^9^, whether humans rely on similar mechanisms or employ dissociable processes for context separation and certainty stabilization remains an open question.

Training schedules offer a powerful lens for addressing this question. Blocked training strengthens within-context associations that may promote local certainty signals^34–37^, but could impair cross-context discrimination^36,38–41^. Conversely, interleaved training promotes discriminable item-level codes supporting transfer, but may preclude certainty-based assembly by forcing high-dimensional representational strategies^35,36,40,42–44^. Here we leverage this dissociation to reveal how learning history shapes the neural mechanisms of knowledge assembly using electroencephalography (EEG). Participants learned two independent transitive orderings under blocked, alternating, or interleaved schedules, followed by minimal boundary training revealing how the two sets are related. We examine the dynamic reactivation of neural manifolds supporting knowledge assembly, how training shapes representational geometry, and test via recurrent neural network (RNN) modeling whether recurrent dynamics and temporal sequencing are critical for assembly. Together, our findings provide a computational account of how different learning shapes the temporal dynamics and representational geometries that enable or constrain rapid knowledge reassembly in humans.

## Results

Forty-eight participants (17 blocked, 15 alternating, 16 interleaved) learned two independent transitive orderings of four novel objects, each assigned an arbitrary “brispiness” rank within its context (Fig. 1A). Training consisted of adjacent pair comparisons with feedback (Fig. 1A), continuing until participants reached at least 90% accuracy within both contexts (Fig. 1B).

**Figure 1.**
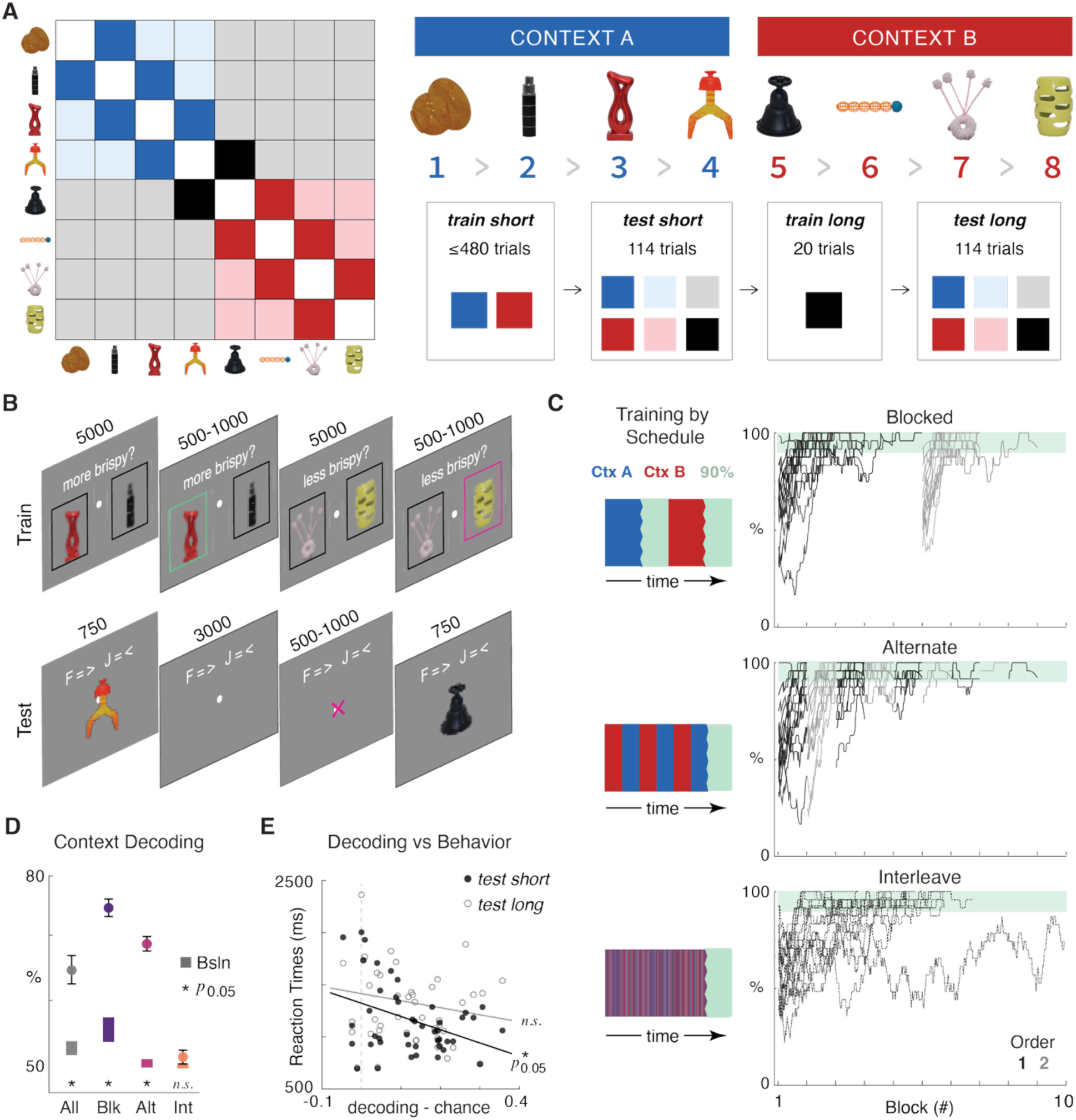
Task design and initial training results (**A**) Left: matrix illustrating training and testing conditions for an example set of objects ordered by rank (on the x and y axes). Each entry indicates a pair of stimuli defined by their row and column. Colors signal when the pair was trained or tested. Dark blue and red squares are within-context pairs, shown during train short. In addition to these, lighter blue and red squares (non-adjacent) and gray (untrained) are within-context pairs not seen during train short. Black squares are the pairs shown during boundary training (train long). All pairs are tested during test short and test long. Right: schematic of experimental sequence and legend. Although we use the same set of objects in these figures for display purposes, note that each participant viewed a unique, randomly sampled set of novel objects. The colored squares refer to the pairs trained or tested in each phase, using color conventions from the leftmost panel. (**B**) Example trial sequence during training (upper) and test (lower). Numbers above each example screen show the frame duration in ms. (**C**) Left panel: Example schematics for how learning proceeded in each of the training schedules, note that context A or B could be encountered first. Right panel: Percentage accuracy in the first encountered (black) and second encountered (gray) contexts. Stopping criterion of 90% is shown as a green rectangle. The excluded interleaved participant is apparent in the bottom panel. A training ‘‘block’’ (x axis) consists of 48 trials. (**D**) We observed above chance context decoding across all participants, as well as within the blocked and interleaved groups compared to within-participant baselines. Error bars indicate SEM for observed decoding accuracy, rectangles represent mean ± SEM within participant baseline decoding accuracies. (**E**) Relationship between context decoding accuracy and behavioral performance during test. Left panel shows significant negative correlation between decoding strength and reaction times (r = -[value], P < 0.05). Right panel shows a non-significant trend relating decoding to accuracy. Filled circles represent short test trials; open circles represent long test trials. Black and gray lines indicate linear fits for significant and non-significant relationships, respectively.

Training schedules differed in temporal structure: blocked schedules required participants to complete one context before starting the second, interleaved schedules intermixed contexts within each training block, and alternating schedules switched contexts across blocks (Fig. 1C). One interleaved participant was excluded for not reaching criterion (Fig. 1C). Remaining participants (N = 47) reached criterion after 4.7 ± 1.4 SD blocks per context (mean ± SD; range 2–9). While overall training trials did not differ across schedules (ME Schedule: *F*_2,82_ = 1.5, p = 0.2) or by context identity (ME Identity: *F*_1,82_ = 0.05, p = 0.8; see Supplementary), blocked and alternating schedules facilitated transfer to the second-encountered context (Fig. S1) (Schedule × Order: *F*_2,82_ = 6.8, p = 0.002; see Supplementary).

Context identity was reliably decoded relative to within-participant baselines across participants (Fig. 1E) (*t*_46_ = 8.1, p < 0.001). Context codes remained stable across time windows (Fig. S1) (6.8 < *t*_46_ < 8.4, p’s < p_FDR_ = 0.001) and leave-one-channel-out analysis suggested distributed cortical encoding (Fig. S1) (see Supplementary). Importantly, context codes emerged only under blocked and alternating schedules (Fig. 1D) (*F*_2,41_ = 28.4, p < 0.001), with interleaved training producing at-chance accuracy (–0.3 < *t*_14_ < 1.4, p’s > 0.2). We next asked whether these context codes influenced the ability to assemble knowledge across separately learned associations. Prior research suggests that not all participants spontaneously integrate distinct associations into unified knowledge structures. However, this previous work assessed assembly only through transitive inference performance during pairwise comparisons. To directly measure explicit structural knowledge, we administered a post-experiment free arrangement task where participants ordered all eight items (Figs. S15, S16) (see Supplementary, see Methods). This approach enabled us to quantify the degree to which participants had assembled a coherent mental representation of the full relational structure. The strength of context codes during the initial training phase did not predict assembly (ME Assembly: *F*_1,41_ = 0.8, p = 0.4; Schedule×Assembly: *F*_2,41_ = 1.1, p = 0.3). Instead, context codes related primarily to performance during *test short* (Fig. S1E) (see Supplementary), suggesting context codes support within-phase performance rather than cross-phase reorganization.

Following training, participants performed a one-back task (*test short*) comparing items along the entire ordering (Fig. 1B). Performance averaged 87.4 ± 10.4% accuracy with 1151.4 ± 321.0ms reaction times (mean ± SD; Fig. 2A). Accuracy did not differ across schedules (Fig. 2C) (*F*_2,44_ = 0.6, p = 0.6), though interleaved training marginally slowed responses (Fig. 2D) (*F*_2,44_ = 3.1, p = 0.05, one-way ANOVAs), consistent with increased retrieval demands^36,45,46^.

**Figure 2:**
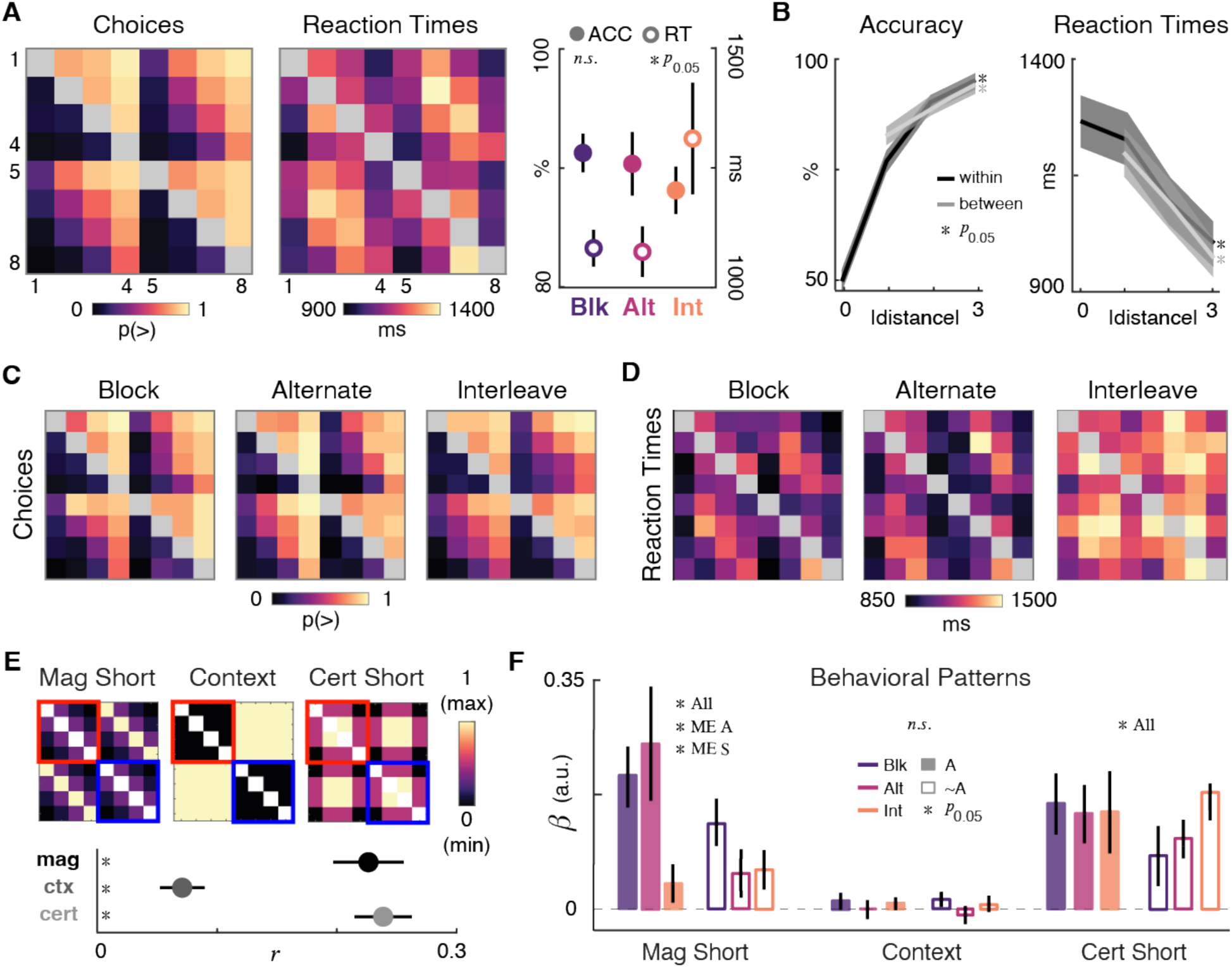
Behavior during *test short* in humans (**A**) Left panel: choice matrix over all participants. The color of each entry indicates the probability of responding “greater than” during *test short* for the pair of items defined by the row and column. Color scale is shown below the plot. Object identities are shown for illustration only (and were in fact resampled for each participant). Middle panel: Similarly, reaction time (RT) matrix shown averaged over all participants. Right panel: Average accuracies (ACC - left ordinate) and reaction times (RT- right ordinate) (**B**) Left panel: accuracy as a function of symbolic distance, shown separately for within-context (e.g., *i*_2_ and *i*_4_; gray dots) and between-context (e.g., *i*_3_ and *i*_6_; black dots) judgments. For between-context judgments, accuracy data are with respect to a ground truth in which ranks are perfectly generalized across contexts (e.g., they infer that *i*_2_ > *i*_8_). Errors bars indicate SEM. Right panels: Equivalent data for reaction times Note that a symbolic distance of zero was possible across contexts (e.g., *i*_1_ vs. *i*_5_) for which there was no “correct” answer, but an RT was measurable. p value indicates significance from paired t tests of RT values. (**C**) Choice matrices averaged within the three *train short* schedules. (**D**) Choice matrices averaged within the three *train short* schedules. (**E**) Top panels: Idealized (I) behavioral matrices for reaction times used for test short. The magnitude matrix reflects item rank, and is scaled linearly with symbolic distance, ranging from 0 (fastest RT for comparisons with symbolic distance = 3) to 1 (slowest RT for comparisons of symbolic distance = 0). Context reflects the context of the item, while certainty reflects position relative to axis center (see Methods). These idealized matrices were used to quantify Human participant behavioral choice and reaction time patterns, and so diagonal axes were set to empty values (indicated with white cells). Bottom panels: Pearson correlations with idealized behavioral matrices over all participants. Error bars indicate SEM. (**F**) Participant RT patterns were predicted by idealized behavioral matrices in a competitive regression. Resultant beta estimates are plotted for idealized magnitude (left), context (middle) and certainty (right) reaction time patterns during the *test short* phase. To assess which behavioral patterns were associated with future assembly, data is shown separately for assemblers (filled bar plots) and non-assemblers (open bar plots). Error bars indicate SEM.

Participants generalized beyond trained pairs – performance exceeded chance for both trained adjacent pairs (82.8 ± 12.9%, *t*_46_ = 44.3, p < 0.001) and untrained non-adjacent pairs requiring transitive inference (Fig. 2A,B) (91.9 ± 12.2%, *t*_46_ = 52.1, p < 0.001). Robust symbolic distance effects (SDEs) emerged, with reaction times decreasing and accuracy increasing as ordinal distance grew (Fig. 2B) (RT: *β* = –114.3 ± 15.6ms/rank, *t*_46_ = –7.3, p < 0.01; Accuracy: *β* = 5.9 ± 0.9%/rank, *t*_46_ = 6.7, p < 0.01; see Supplementary). Participants also displayed a natural tendency to match rank orderings between contexts (e.g. that the 2^nd^ item was ranked higher than the 4^th^, regardless of set) in the absence of information about how objects were related. We quantified between-context accuracy relative to an agent that generalizes ranks perfectly between contexts (as in Nelli et al 2023; see Methods), and found participants showed above-chance performance on cross-context comparisons for both adjacent (77.1 ± 15.1%, *t*_46_ = 35.2, p < 0.001) and non-adjacent pairs (92.5 ± 11.5%, *t*_46_ = 55.5, p < 0.001) indicating spontaneous rank alignment across sets (Fig. 2A). Similarly, SDEs were evident across contexts (Fig. 2B; RT: *β* = –112.5ms/rank, *t*_46_ = –6.9, p < 0.001; Accuracy: *β* = 8.9%/rank, *t*_46_ = 9.4, p < 0.001).

To decompose behavioral strategies, we correlated reaction time matrices with three idealized *test short* models: magnitude (ordinal distance), context (set identity), and certainty (distance from center; see Methods). All three correlated with observed behavior (Magnitude Short: *t*_46_ = 7.7, p < 0.001; Context: *t*_46_ = 3.8, p < 0.001; Certainty Short: *t*_46_ = 10.0, p < 0.001). However, competitive regression revealed magnitude and certainty dominated responses (Magnitude Short: *t*_46_ = 5.7, p < 0.001; Certainty Short: *t*_46_ = 7.3, p < 0.001), while context was no longer apparent (Fig. 2E,F) (*t*_46_ = 1.0, p > 0.3). Critically, participant behavior more closely reflecting the parallel rank ordered lines predicted assembly in the next phase of the experiment (ME Assembly: *F*_1,41_ = 4.9, p = 0.03) and was enhanced under blocked initial training (ME Schedule: *F*_2,41_ = 4.0, p < 0.03; Schedule×Assembly: *F*_2,41_ = 2.8, p = 0.07). We find that blocked training boosts behavioral signatures of rank ordered structure that predict future knowledge restructuring, rather than context or uncertainty (*F*s < 0.9, p’s > 0.4).

Training schedules modulated neural processing as measured through event-related potentials (ERPs) (Fig. 3A, Fig. S3). At frontal-central sites (Cz, FCz), blocked training enhanced N1 and reduced P2 component amplitudes (ME Schedule - Cz: 4.9 < *F*_2,41_ < 20, p’s < p_FDR_ = 0.002, 127– 183ms; FCz: 5.8 < *F*_2,41_ < 11.2, p’s < p_FDR_ = 0.007, 135–178ms). These modulations are consistent with efficient sensory gain control and reduced perceptual disambiguation demands when categories are temporally separated^47–49^. Blocked training also elicited stronger N400 amplitudes at these same sites (Fig. 3A, Fig. S3) (ME Schedule – Cz: 3.5 < *F*_2,41_ < 13.6, p’s ≤ p_FDR_ = 0.002, 253–519ms; FCz: 5.6 < *F*_2,41_ < 8.6, p’s ≤ p_FDR_ = 0.007, 284–387ms). The N400 component indexes prediction error and semantic processing demands^50–54^, suggesting blocked schedules requires greater integration during unexpected, novel inferences due to strong predictive processing. At posterior site PoZ, future assemblers showed enhanced P3b amplitudes following interleaved and alternating, but not blocked, training (Fig. 3A,B) (Schedule×Assembly: 3.5 < *F*_2,41_ < 6.8, p’s ≤ p_FDR_ = 0.04, 204–489ms; ME Schedule and Assembly p’s > 0.05). As the P3b component reflects attentional allocation and memory updating processes^55,56,56,57^, this pattern suggests that flexible knowledge assembly following interleaved schedules may require heightened engagement during the initial learning phase.

**Figure 3:**
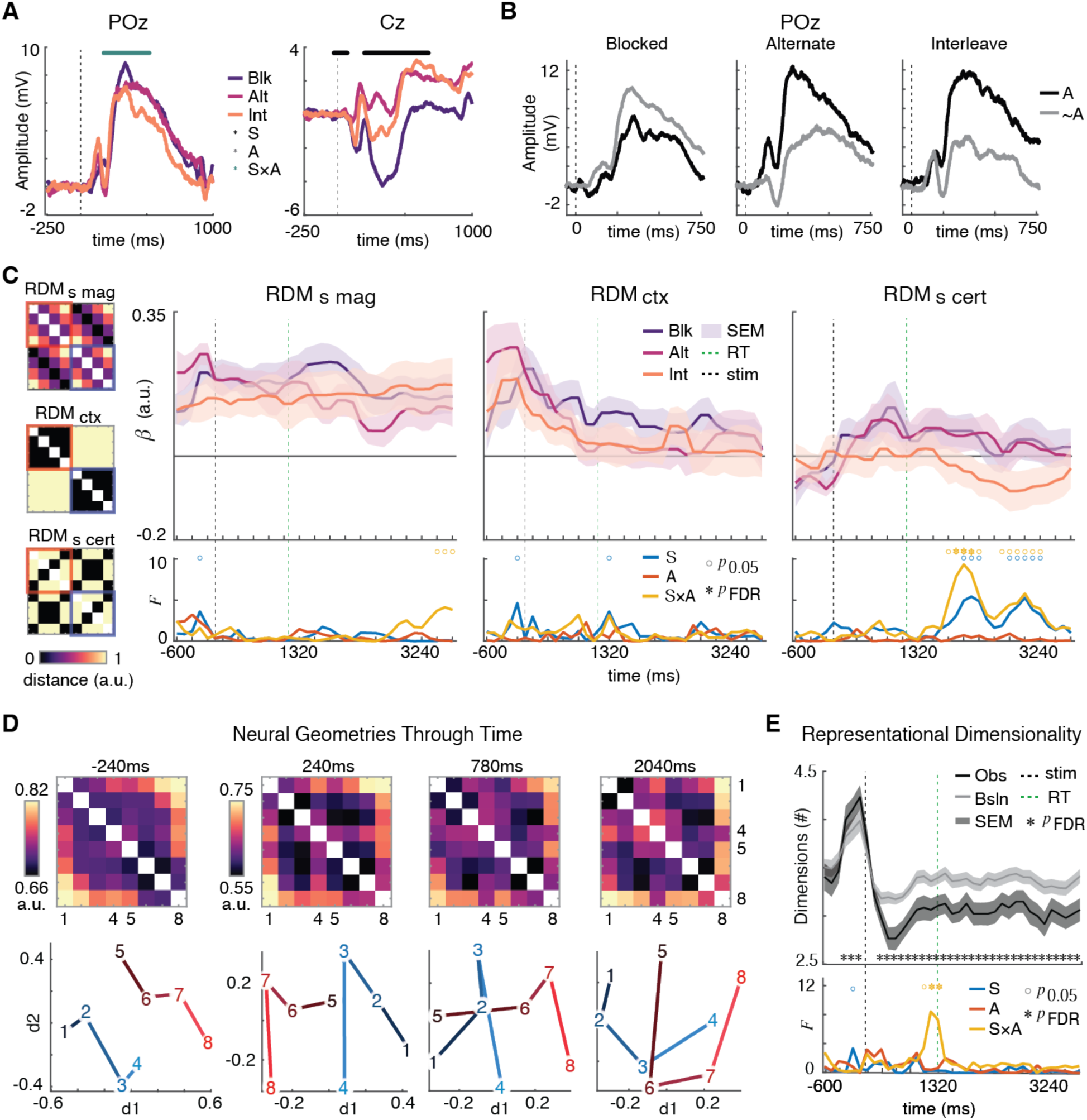
Neural processing and geometries during *test short* (**A**) Event-related potentials (ERPs) at centro-parietal (POz) and central (Cz) electrodes reveal training schedule-dependent processing. Blocked (purple) alternating (pink) and interleaved (orange) schedules are plotted. Significance is shown for 2-way ANOVA comparing Schedule (S), assembly (A), and interaction (S×A). Vertical dashed line indicates stimulus onset. (**B**) ERPs at POz in participants who would go onto assemble knowledge (A, black) versus non-assemblers (∼A, gray) across training schedules. Blocked training shows attenuated prediction error responses to violations compared to alternating and interleaved schedules, suggesting reduced context sensitivity and weaker predictive processing. (**C**) Multivariate representational dynamics indexed by three model RDMs. Top panels show beta weights over time corresponding to how well each RDM predicts neural patterns across blocked (purple), alternating (pink), and interleaved (orange) schedules. RDM_s mag_ captures item-level magnitude similarity, RDM_ctx_ captures contextual structure, and RDM_s cert_ captures the U-shaped certainty structure within each of the 2 sets. Bottom panels show F-statistics testing differences between Schedule (S) and assembly (A, S×A), with significance markers indicating when neural patterns discriminate these conditions (p < 0.05, FDR-corrected. (**D**) Neural geometries evolve through time. Top row shows RDMs averaged over all participants at -240ms, 240ms, 780ms, and 2040ms relative to stimulus onset. Bottom row shows corresponding projections via multidimensional scaling, with numbered nodes representing the 8 learned items. Early (240ms) representations are clearly organized by magnitude and separated by context, and late (2040ms) representations stabilize into certainty-weighted relational configurations. (**E**) Representational dimensionality collapses over time. Top panel tracks effective dimensions over time across conditions. with interleaved future-assemblers maintaining higher dimensionality late into the trial. Bottom panel shows F-statistics identifying robust late differences during successful interleaved learning (marked by asterisks, p < 0.05, FDR-corrected).

To characterize neural knowledge geometries, we computed representational dissimilarity matrices (RDMs) at each timepoint and correlated them with three model RDMs capturing hypothesized *test short* coding schemes (Fig. 3C) (see Methods). RDM_s mag_ indexed magnitude structure^9^, with parallel, rank-ordered manifolds for each context and RDM_ctx_ reflected context identity through categorical distinctions between sets. RDM_s cert_ quantified certainty-weighted structure^33^, capturing a U-shaped pattern in which representations were most distinct at the endpoints of each dimension but most similar (most confusable) in the center. Across participants, aligned magnitude manifolds persisted throughout the trial (Fig. S3) (4.4 < *t*_46_ < 9.1, p_FDR_ = 0.001). Context codes were apparent during stimulus onset but decayed near responses (2.5 < *t*_46_ < 6.1, p’s ≤ p_FDR_ = 0.02), while certainty-related curvature emerged only briefly before responses (720–840ms, 2.3 < *t*_46_ < 2.7, p’s ≤ p_FDR_ = 0.02; t-tests on correlations; see Supplementary).

Notably, representational geometries during intertrial intervals predicted assembly success in next phase of the experiment (*test long*) (Fig. 3C, Fig. S3D). Under blocked or alternating training schedules, future assemblers first expressed certainty manifolds (ME Schedule: *F*_1,41_ = 7.8–9.4, p < p_FDR_ = 0.001, 1920–2160ms; see Supplementary), followed by marginally stronger magnitude coding (ME Assembly: *F*_1,41_ = 3.7–4.2, uncorrected p’s < 0.03, 3480–3720ms). This temporal sequence suggests U-shaped certainty manifolds may stabilize magnitude representations during rest periods, with these processes proving integral for subsequent reorganization. In contrast, interleaved future-assemblers did not exhibit post-decision certainty curvature (Fig. 3C, Fig. S3), displaying factorized representations instead (Fig. S4). Consistent with this, interleaved assemblers showed elevated representational dimensionalities (Fig. S3), particularly near the time of responses^58^ (*F*_2,41_ = 7.4–8.5, p’s < p_FDR_ = 0.002, 1080-1200ms), compared to effectively low dimensionalities observed across all participants (Fig. 3E) (-5.4 < *t*_46_ < -3.8, p’s < p_FDR_ = 0.03; see Supplementary). These findings indicate that dynamic dimensionality expansion can enable flexible integration without relying on relational certainty. Despite robust encoding during stimulus processing, context codes did not forecast assembly success (Fig 3C; *F*_1,41_ < 2.3, p’s > 0.1), suggesting context representations are not necessary for future reorganization (see Supplementary).

After initial testing, participants completed minimal boundary training (*train long*; Fig 1A) in which they learned that object *i*_4_ (least brispy object in context A) was more brispy then object *i*_5_ (most brispy object in context B). This information was acquired over just 20 trials in which participants repeatedly judged whether item *i*_4_ or *i*_5_ was more or less brispy. Performance averaged 89.8 ± 6.5% (mean ± SD), was unaffected by schedule, and did not determine subsequent assembly (*F*’s < 2.0, p’s > 0.2; see Supplementary).

Following boundary training, participants entered the *test long* phase (Fig. 1A) which had identical structure to *test short*. Performance declined relative to *test short* (Fig. 4A,E) (Accuracy: *t*_46_ = 6.4, p < 0.001; RTs: –4.6, p < 0.001; paired t-tests), driven by non-assemblers who failed to integrate the sets (Fig 4A) (ME Assembly: *F*_1,41_ = 62.8, p < 0.001; see Supplementary). Overall accuracy (Fig. 4A) (ME Schedule: *F*_2,41_ = 2.1, p > 0.1; Schedule×Assembly: *F*_2,41_ = 0.05, p = 0.9) and assembly rates (Fig. 4C) (χ²_(2, *N*=47)_ = 0.06, p > 0.9) did not differ by schedule. However, a more nuanced pattern emerged when examining cross-context comparisons. Schedule specifically impacted between-context comparisons that required integration (Fig. 4B) (ME Schedule: *F*_2,41_ = 4.0, p = 0.03) but not within-context judgments (Within: *F*_2,41_ < 0.09, p > 0.9; see Supplementary). In line with this pattern, both blocked training and successful assembly enhanced the efficiency of cross-context judgments relative to within-context judgments (Fig. 4D,E) (ME Schedule: *F*_2,41_ = 6.8, p = 0.003; ME Assembly: *F*_1,41_ = 22.6, p < 0.001; Schedule×Assembly: *F*_2,41_ = 3.5, p = 0.04). To test whether these effects replicated under different attentional demands, we conducted a larger online study (N = 140; see Methods). Interleaved learners showed reduced *test long* accuracy (ME Schedule: *F*_2,134_ = 3.6, p = 0.03; ME Assembly: *F*_1,134_ = 43.9, p < 0.001. Schedule×Assembly: *F*_2,134_ = 4.8, p < 0.01) and lower assembly rates (Fig. 4C) (χ²_(2, *N*=140)_ = 6.3, p = 0.04). This suggests interleaved learning may require optimal attentional conditions to support higher-order relational inference. Behavioral data combined across both experiments revealed distinct subgroups of non-assemblers, highlighting the diversity of inferences drawn from few-shot boundary training (Fig. S8; see Supplementary).

**Figure 4:**
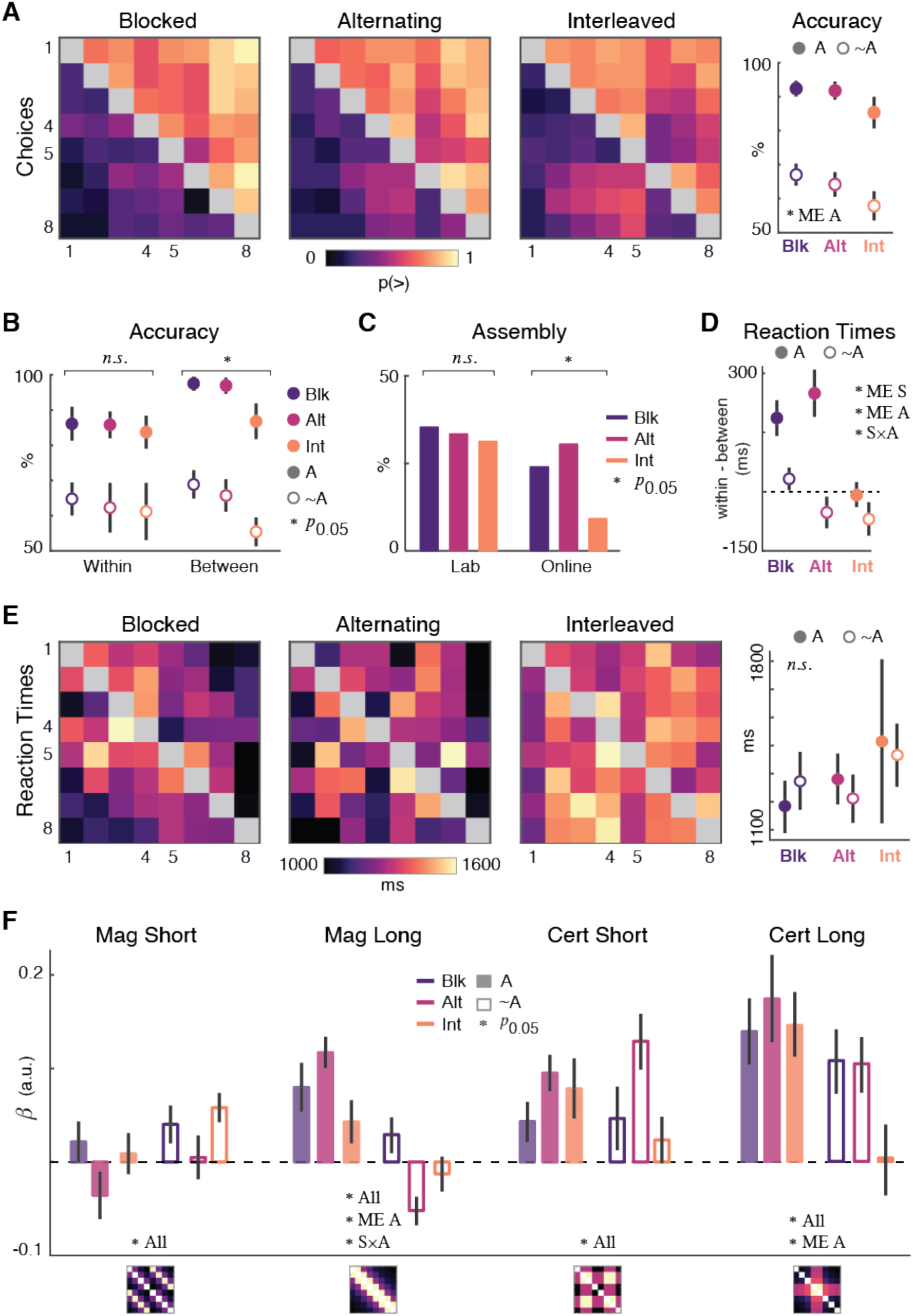
Behavioral reorganization during *test long*. (**A**) Left panels: Choice matrices for the three training schedules during the test long phase. The color of each entry indicates the probability for selecting “more” for an object given the pair defined by the row and column. Color scale shown at right. Right panel: Accuracy averaged over assemblers (A; closed circles) and nonassemblers (∼A; open circles) under the three initial training schedules. Error bars indicate SEM. (**B**) Accuracy for within-context (items from the same original context) and between-context (items from different original contexts) comparisons across the three schedules. Alternating schedule shows significantly lower accuracy for between-context judgments compared to within-context judgments. Asterisks denote significance for the main effect of schedule: * P < 0.05. Error bars indicate SEM. (**C**) Assembly rates (percentage of participants achieving successful knowledge assembly) for lab and online cohorts across schedules. Interleaved schedule shows notably lower assembly in the online cohort. Asterisks denote significance for Chi-squared test for uniformity: * P < 0.05. (**D**) Reaction Times show distinct patterns in decision efficiency between schedules as well as by assembly status. Main effects of assembly, schedule, and an interaction are observed. (**E**) Left panels: Reaction time (RT) matrices for the three schedules during test long. Color scale indicates RT in milliseconds. Right panel: Accuracy averaged over assemblers (A; closed circles) and nonassemblers (∼A; open circles) under the three initial training schedules. (**F**) Beta coefficients from idealized behavioral matrix regressions predicting choice patterns, shown separately for magnitude (mag) and certainty (cert) structures adhering to the short and long phases of the experiment. Analyses examine main effects of schedule (ME S), assembly status (ME A), and their interaction (S×A). Test long magnitude and certainty are both increased with assembly, with an interaction such that blocked and alternating schedules show enhanced elongated magnitude reaction times. Error bars indicate SEM. Asterisks denote significance: * p < 0.05.

To further decompose behavioral strategies, we tested responses against idealized matrices from both test phases (Fig. 4F, Fig. S5). Across participants, behavior correlated with magnitude and certainty models from both phases (Mag Long: *t*_46_ = 3.4, p = 0.001; Cert Long: *t*_46_ = 4.1, p < 0.001; Mag Short: *t*_46_ = 2.7, p < 0.01; Cert Short: *t*_46_ = 3.3, p = 0.002), but not context (*t*_46_ = -0.9, p > 0.3; t-tests on correlations). Assembler responses reflected an integrated ordering (Fig. S5) (ME Assembly: *F*_1,41_ = 44.7, p < 0.001) while non-assembler behavior adhered to the previous task’s structure (ME Assembly: *F*_1,41_ = 10.5, p < 0.01). Competitive regression confirmed that assembler responses reflected integrated magnitude and certainty (Fig. 4F) (ME Assembly: Mag Long – *F*_1,41_ = 27.7, p < 0.001; Cert Long – *F*_1,41_ = 6.4, p < 0.001), rather than magnitude, certainty and context from the previous task phase (p’s > 0.2; see Supplementary). Notably, blocked training induced behavior patterns reflecting assembly in non-assemblers (Fig. 4F, Fig. S5) (Mag Long – Schedule×Assembly: *F*_2,41_ = 4.7, p = 0.01; see Supplementary). Behavioral integration in blocked nonassemblers was statistically indistinguishable from interleaved assemblers (Fig. 4F) (see Supplementary), suggesting blocked learning enables implicit structural inference despite explicit assembly failure.

To investigate the neural geometries underlying knowledge assembly, we first tested whether neural representations reflected updated magnitude (RDM_l mag_) and certainty (RDM_l cert_) (Model 1; see Methods). Across participants, neural representations aligned with the integrated 8-item ordering throughout the trial (Fig. 5A,B, Fig. S10A) (4.7 < *t*_46_ < 24.4, p’s < p_FDR_ = 0.03, -600- 3720ms; t-tests against zero) with enhanced persistence under blocked training (Fig. 5A) (ME Schedule: 1920–2160ms, 2520–3480ms; 4.9 < *F*_2,41_ < 9.0, p’s ≤ p_FDR_ = 0.02). Curvature along the eight-item ordering emerged later in the trial period (Fig. 5B, Fig. S10) (2.2 < *t*_46_ ≤ 5.3, p’s ≤ p_FDR_ = 0.03, 1080–3720ms; t-tests against zero).

**Figure 5:**
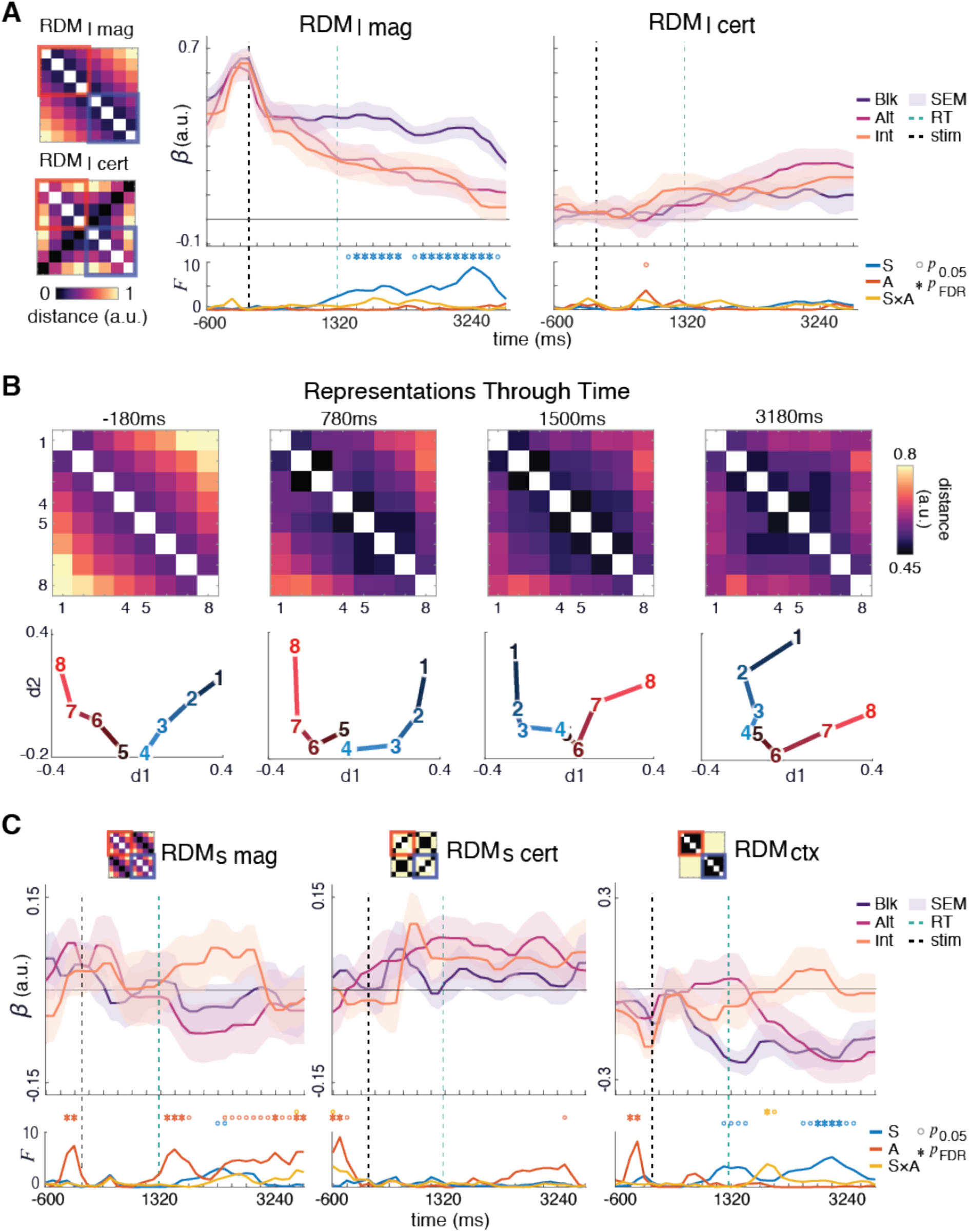
Neural reorganization during *test long*. (**A**) Multivariate representational dynamics indexed by model RDMs for an updated magnitude structure (RDM_l mag_) and certainty (RDM_l cert_). Top panels show beta weights over time corresponding to how well each RDM predicts neural patterns across blocked (purple), alternating (pink), and interleaved (orange) schedules. RDM_l mag_ captures the elongated item-level magnitude similarity and RDM_l cert_ captures one elongated U-shaped certainty structure, instead of two as in *test short.* Bottom panels show F-statistics testing differences between Schedule (S) and assembly (A, S×A), with significance markers indicating when neural patterns discriminate these conditions. Significance level of p < 0.05 marked by dots, FDR-corrected marked by asterisks (*). (**B**) Neural geometries evolve through time. Left panel: Top row shows RDMs averaged over all participants at -180ms, 780ms, and 3180ms relative to stimulus onset. Bottom row shows corresponding projections via multidimensional scaling, with numbered nodes representing the 8 learned items. Early (240ms) representations are clearly organized by magnitude, and late (2040ms) representations stabilize into certainty-weighted relational configurations. (**C**) Temporally specific re-activation of *test short* representational geometries is associated with assembly. Top panels show beta weights over time corresponding to how well each RDM predicts neural patterns across blocked (purple), alternating (pink), and interleaved (orange) schedules. Bottom panels show F-statistics testing differences between Schedule (S) and assembly (A, S×A), with significance markers indicating when neural patterns discriminate these conditions. Significance level of p < 0.05 marked by dots, FDR-corrected marked by asterisks (*).

Next, we asked whether assembly depends on reactivating prior representational geometries. To do this, we entered participant RDMs into competitive regressions including both test phases’ magnitude codes alongside certainty and context models from *test short* (Model 2; see Methods). According to the context account, binary context codes would easily allow reorganization of orderings from *test short*^9^. On the other hand, reactivation of relational certainty may help to maintain initially learned relations among items by rendering them less labile as the global geometry of the task changes^9^. This analysis revealed assembly was predicted by dynamic reactivation of *test short* geometries in specific temporal windows (see Supplementary). Pre-stimulus prior-task certainty codes supported assembly (Fig. 5C) (–600 & –480ms, 6.7 < *F*_1,41_ ≤ 9.8, p’s < p_FDR_ = 0.03), suggesting preparatory certainty signals may encourage flexible integration by indicating which relational weights to maintain versus update. In contrast, pre-stimulus magnitude geometries from *test short* interfered with reorganization (–240 & –120ms, *F*_1,41_ = 6.4–8.0, p’s < p_FDR_ = 0.03), aligning temporally with the period of strongest elongated magnitude representations (Fig. 5A, Fig. S10). This finding that parallel magnitude manifolds during stimulus processing hinder assembly is consistent with fMRI evidence, and potentially reflects rigid adherence to previously learned structure^9^. Strikingly, negative correlations with context models predicted assembly (–360 & –240ms, *F*_1,41_ = 5.7–8.8, p’s < p_FDR_), indicating successful reorganization requires integrating contexts rather than maintaining context-dependent codes. Following decisions, reactivation of initial task magnitude structure facilitated assembly (Short Mag: 1440–1680ms, *F*_1,41_’s = 5.6–7.4, p’s < p_FDR_ = 0.03; 3240–3720ms, *F*_1,41_’s = 6.6–6.9, p’s < p_FDR_= 0.03). Similarly, certainty along the two sets now marginally interfered with assembly (Short Cert: 3360ms, *F*_1,41_ = 4.4, p = 0.04 > p_FDR_ = 0.03) temporally coinciding with maximal expression of the integrated U-shaped certainty structure (Fig. 5A, Fig. S10). This temporal reversal suggests that while pre-stimulus magnitude codes constrain flexibility, post-decision reactivation may consolidate the newly reconfigured structure by reintegrating prior knowledge with updated relations.

To quantify whether representational complexity reflected the low-dimensional ground truth ordering, we computed effective dimensionality (observed dimensionality relative to within-participant baselines; see Methods). Neural dimensionality decreased from *test short* to *test long* (Fig. S10) (2.2 < *t*_46_ < 5.3, p’s < p_FDR_ = 0.03), supporting low-dimensional reorganization instead of context-dependent gating. After correction for multiple comparisons, neither schedule nor assembly predicted test long dimensionalities (Fig. S10) (F’s < 6.4, p’s > p_FDR_ = 0.002; see Supplementary). Instead, changes in dimensionality from *test short* to *test long* revealed schedule-dependent assembly mechanisms – successful assembly under blocked training was marked by lower dimensionalities during *test short* and higher dimensionalities during *test long* (Fig. S10) (Schedule×Assembly: *F*_2,42_ = 12.6, 9.2, p’s < p_FDR_ = 0.002, 1201ms, 1321ms; see Supplementary). This pattern suggests blocked learners initially form compressed, low-dimensional representations that subsequently expand during reorganization. In contrast, high-dimensional initial representations may enable assembly under interleaved conditions by maintaining discriminable representations that can be flexibly reconfigured. These dynamics demonstrate that training history constrains the complexity of neural codes available for knowledge assembly.

To identify whether recurrent dynamics are enough to induce human-like knowledge assembly and whether blocked training naturally encourages the formation of certainty weighted structures, we trained vanilla recurrent neural networks (RNNs) on the magnitude ordering task (Fig. 6A). Models underwent *train short* according to Blocked (N = 10) or Interleaved (N = 10) curricula, where training trials from each context were presented either in sequential blocks or randomly interleaved (see Methods). During *test short*, models captured several aspects of human performance, including learning magnitude relationships (Fig. S11) and above-chance generalization to novel cross-context comparisons (Fig. S11). Additionally, RNNs represented ordinal information in a systematic way across hidden layer representations, displaying parallel neural axes for magnitude across contexts (Fig. 6D, Fig. S12)^9,59,60^. However, RNNs failed to reproduce the U-shaped neural geometries characteristic of certainty-weighted magnitude representations observed in humans (Fig. 3D), even under blocked training schedules. This dissociation reveals that temporal structure alone is insufficient to generate certainty codes.

**Figure 6:**
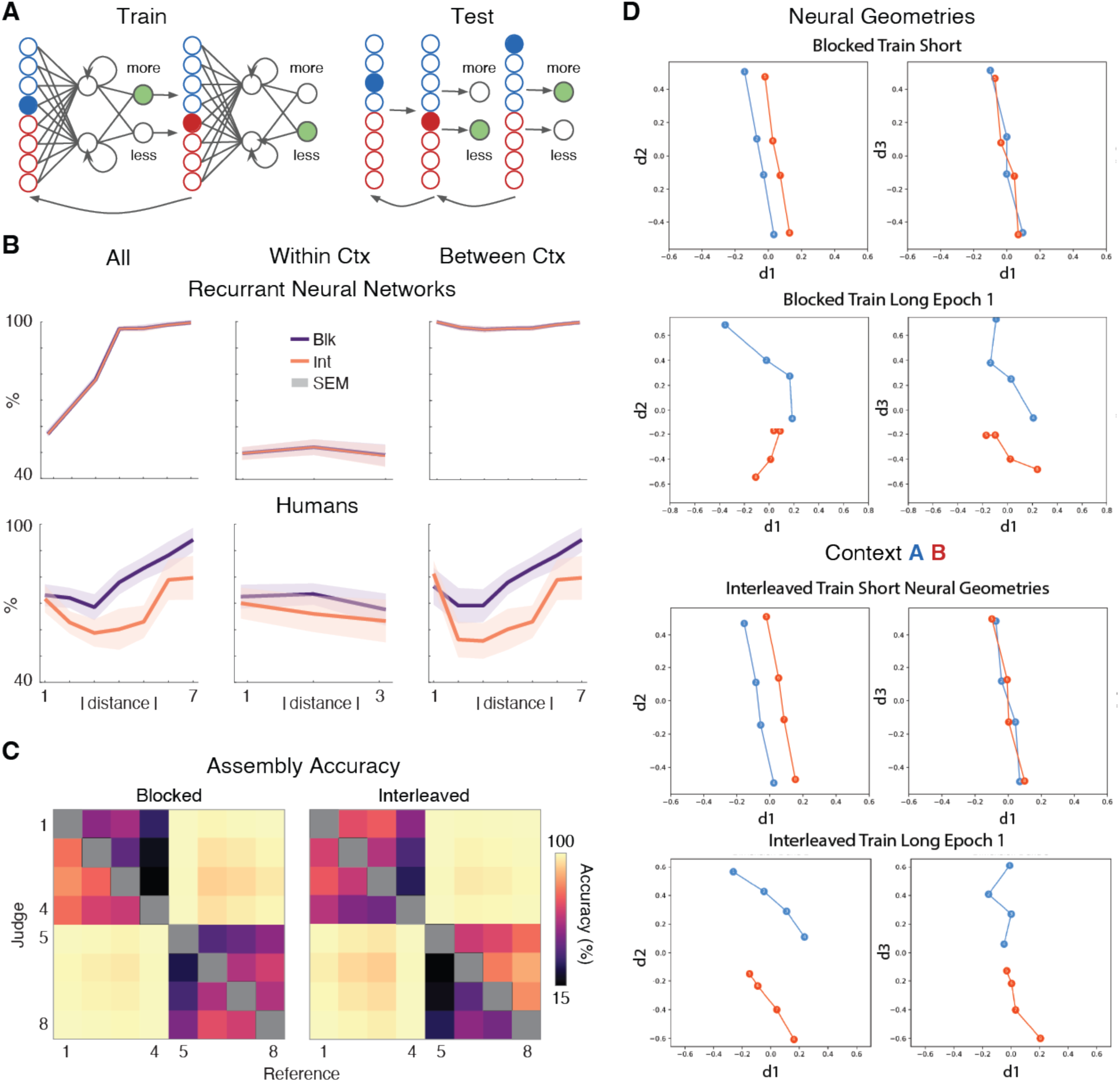
Recurrent Neural Networks Partially Assemble Knowledge. **(A)** Schematic of example training (left) and testing (right) trials. Filled blue and red circles indicate the item pair being trained on each trial. Gray arrows show one-back comparisons between successive trials. Filled green circles indicate the correct response for the example comparisons shown. **(B)** Symbolic Distance Effects during *test long* for recurrent neural networks (top row) under blocked (purple) and interleaved (orange) initial training. Human symbolic distance effects shown (bottom row) for comparison. Shaded regions indicate standard error of the mean. **(C)** Accuracy matrices averaged over blocked (left - N=10) and interleaved (right - N=10) training. Rows indicate the current judgement item; columns indicate the previous reference item. Yellow indicates high accuracy; purple indicates low accuracy. **(D)** Multidimensional scaling (MDS) visualizations after blocked (top rows) and interleaved (bottom rows) test short (1st and 3rd rows) and test long (2nd and 4th rows) training. Test short data show network representations after reaching 90% training accuracy. Test long data show neural representations after training epoch 1. Blue represents items initially trained in context A, red represents items from context B.

RNNs successfully incorporated boundary information during testing, achieving ceiling-level accuracy on between-context comparisons (Fig. 6B,C, Fig. S13). This demonstrates that recurrent networks can restructure knowledge when new relational information is introduced, a capacity absent in feedforward networks^9^. Recurrent connections may enable this flexibility by expanding network dimensionality in a manner analogous to dimensionality expansion mechanisms observed in human learners. However, RNN models exhibited marked decline in within-context performance (Fig 6B). Models showed complete performance inversion for these trial types (Fig. 6C, Fig. S13; see Supplementary), indicating that their restructuring mechanism differed fundamentally from that of humans (Fig. 4A, Fig. S8). Analysis of training dynamics revealed that model accuracy peaked after only 1-3 training epochs on the boundary relation (Fig. S12). Continued training beyond this point progressively degraded representational structure (Fig. S12), suggesting that RNNs learn new cross-context relationships at the expense of established within-context knowledge. This interference proved insensitive to initial training schedule: models showed no behavioral differences between blocked and interleaved curricula (Fig. 6B,C) even on cross-context comparisons (-0.3 < *t*_18_ < 0.4, p’s > 0.7; see Supplementary). This contrasts sharply with the schedule-dependent neural and behavioral effects observed in humans.

The progressive degradation of RNN representations might superficially suggest that assembly requires early stopping mechanisms. However, early stopping cannot explain biological learning for two critical reasons. First, unlike vanilla networks which catastrophically forget when trained on new tasks, humans demonstrably engage in continual learning without drastically overwriting prior knowledge^18,20,20,61–63^. Second, and more fundamentally, extended learning in humans produces increased discrimination between learned items—the opposite of what occurs in neural networks during extended training^64–66^. Instead, our results point to within-context certainty processes absent from vanilla RNNs, consistent with the computational framework proposed by Nelli et al. (2023). These mechanisms would enable biological systems to solve the stability-plasticity dilemma that vanilla RNNs cannot: maintaining established within-context knowledge while flexibly incorporating new cross-context relations. The certainty reactivation we observed in humans (Fig. 5C) provides direct evidence for such specificity—without comparable mechanisms, RNNs resort to wholesale rewriting.

## Conclusion

Our findings reveal that cognitive flexibility emerges through multiple neural routes shaped by learning history. Training schedules bias learners toward distinct representational strategies, each capable of supporting successful assembly. Blocked training promoted compressed certainty codes through temporal separation, yielding low-dimensional representations that encoded relational reliability via U-shaped manifolds. Interleaved training interfered with local certainty coding, instead promoting higher dimensional, discriminable codes that persisted across both test phases. P3b amplitudes similarly suggest that assembling knowledge under interleaved training requires additional cognitive resources, while blocked training relies on more efficient retrieval processes^56,57,67^. In sum, representational geometries are shaped by the temporal statistics of learning, highlighting multiple neural routes to flexible knowledge assembly biased by training regime.

The temporal specificity of representational reactivation reveals how the brain orchestrates knowledge restructuring. Pre-stimulus certainty reactivation predicted assembly, consistent with preparatory uncertainty signals guiding plasticity by indicating which representations to stabilize versus modify^9^. Reactivation of prior magnitude codes during stimulus processing enforced rigid adherence to the preceding tasks’ structure, preventing the flexibility necessary for reorganization; extending fMRI evidence of interference to reveal its temporal locus^9^. Post-response magnitude reactivation, however, may serve a consolidation function, suggesting assembly involves two stages: flexible reconfiguration during processing followed by stabilization during post-decision periods. While temporal blocking encouraged context codes during initial learning, these codes did not facilitate assembly. Instead, they may help learners establish context-bound certainty weighting without directly contributing to reorganization.

The failure of vanilla RNNs to develop curved certainty-weighted geometries despite blocked training points to missing computational mechanism. We propose several candidates for consideration in future work. Differential dendritic activation in pyramidal neurons could support independent encoding of rate-based magnitude and rank selectivity, whose interaction naturally produces curved manifolds^17,19,33^. Certainty-dependent learning rates akin to Bayesian confidence weighting could create stronger synaptic stability at endpoints where confidence is high, explaining both U-shaped geometries and protection against catastrophic interference^68,68–71^. Finally, metacognitive monitoring tracking comparison difficulty could dynamically modulate representational geometry and plasticity ^72,73,73^. Direct neural recording in animals trained on analogous tasks could characterize synaptic plasticity rules for high-versus low-certainty comparisons, bridging cellular and circuit implementation with computational-level representational transformations.

These findings intersect with multiple theoretical frameworks emphasizing the role of structured learning signals^74,75^. While complementary learning systems theory proposes rapid hippocampal encoding followed by slow cortical consolidation^6,7^, our results suggest cortical certainty processes provide an independent mechanism for rapidly restructuring existing representations^9^. The structure learning literature emphasizes distributional statistics and schema formation as drivers of transfer^45,76^. We extend this work by demonstrating that temporal organization of experience, not merely statistical properties, determines available representational strategies. This has implications for educational practice: blocked training builds strong within-domain certainty while interleaved training enhances discrimination and transfer flexibility. Finally, we link U-shaped manifolds to behavioral certainty, unifying literature concerning parietal evidence accumulation^77–79^ and representational geometries^9,30,60^ and bridging these neural mechanisms with behavioral work on positional effects during inference^80–82^. These findings suggest certainty-based reactivation reflects fundamental mechanisms within cortical circuits regulating decision making^33^.

Several limitations warrant consideration. Our paradigm employed implicit context manipulated through temporal structure, so whether explicit context cues facilitate assembly, particularly under interleaved training, remains unknown. Additionally, the generality of prior-task reactivation also requires exploration. Are previous geometries reactivated during continual learning with multiple sequential tasks, and if so, how does the brain select which representations to reinstate? These questions are particularly relevant for understanding catastrophic interference and lifelong learning in both biological and artificial systems.

Knowledge assembly represents a fundamental component of flexible cognition, enabling rapid restructuring of established knowledge in light of new information. Our findings demonstrate that this ability depends not just on having appropriate prior knowledge, but critically on *when* that knowledge is represented and reactivated in neural population codes. Successful reorganization emerges through dynamic orchestration of neural codes, with temporally specific reactivation supporting flexible reconfiguration while maintaining appropriate stability. Training history constrains available representational formats, biasing learners toward compressed certainty-based or dimensionality expansion strategies. This work demonstrates that cognitive flexibility emerges from the creative reuse of learned representations, precisely timed to balance plasticity with stability, with implications for educational practice and AI research alike.

## Methods

### Participants

Fifty-three adult participants were recruited for the experiment via the online recruitment system (SONA) at Occidental College and flyers posted around the campus, and all subjects had normal or corrected to normal vision. Five participants were excluded from final analyses due to technical difficulties with EEG equipment. For four of these participants, the session was terminated early due to pervasive 60 Hz noise, while one participant was excluded due to a failure of the EEG trigger system. One additional participant was excluded for failure to reach performance threshold during the *train short* phase. All analyses were performed on the remaining 47 participants (28 females, 14 males, and 4 non-binaries), with a mean age of 19.7 ± 1.18 years old (range 18 to 23). Participants received either course credit through Occidental college’s SONA system or $15/hour monetary compensation. All procedures were approved by the Occidental College Human Subjects Research Review Committee, and written informed consent was obtained.

### Stimulus and Task

The experiment was implemented using the Psychophysics Toolbox-3 (Kleiner et al., 2007) for MATLAB 2020a (MathWorks) and additional custom scripts on a desktop computer running Windows 10. Stimuli were novel, nonsense objects drawn from the NOUN database (Horst & Hout, 2016). Out of the 60 possible images in this database, objects that were rated as most similar to the others (e.g., an average similarity rating within 1 standard deviation of the maximum) and objects that were rated as most familiar (e.g., scoring less than 50% on an inverse familiarity score) were excluded, leaving 37 possible objects. For each participant, 8 out of the 37 objects were randomly selected. These selected objects were arbitrarily assigned a rank from 1 to 8, with 1–4 corresponding to objects assigned to set A and 4–8 denoting objects assigned to set B.

Participants engaged in a transitive inference task during the EEG recording. The task involved four experimental sessions, *train short*, *test short*, *train long* and *test long*. The first training phase, *train short*, consisted of a minimum of 2 and a maximum of 10 blocks of 48 trials each. Participants performed this task until they reached a criterion of at least s correct on both context, averaged over the preceding 48 trials (e.g. 1 block) from each context. Upon beginning the task, participants were assigned to one of three *train short* learning schedules. The fully blocked condition, intended to provide the most contextual certainty, and could consist of up to 5 consecutive blocks of 48 trials for each context. In the alternating condition, the two contexts are presented in alternating blocks of 48 trials. Lastly, in the fully interleaved condition trials from both contexts occurred within the same 48 trial training block.

During the *train short* phase, participants could encounter objects from context A (i_1_ - i_4_) or context B (i_5_ - i_8_) first, and this was randomized across participants. For all participants, each trial began with the presentation of two objects drawn from adjacent ranks within a single context (e.g. i_3_ and i_4_ or i_7_ and i_8_). These objects were shown either side of a central fixation point, and above the point the words ‘‘more brispy?’’ or ‘‘less brispy?’’. After a 200ms fixation delay, two stimuli were presented in each hemifield and participants were instructed to select the corresponding object (i.e. that which was more or less brispy) using either the ‘‘F’’ (left object) or ‘‘J’’ (right object) keys. These objects remained on screen for 5000ms, and once a response was recorded, a green (correct) or red (incorrect) box would appear around the selected object to indicate whether it was the correct selection. During this phase of the experiment, participants responded with average reaction times (RTs) of 1498.3ms ± 37.4 SD; if participants did not respond within 5000ms of stimulus presentation, participants saw a red “x” and the trial was considered incorrect. Either way, feedback persisted for a variable ITI of 500-1000ms before the next trial. Critically, participants were only trained to compare the 3 consecutive object pairs within each context (*i*_1_ – *i*_2_, *i*_2_ – *i*_3_ and *i*_3_ – *i*_4_ from context A and *i*_5_ – *i*_6_, *i*_6_ – *i*_7_ and *i*_7_ – *i*_8_ from context B). Object hemifield, the trial-order of each object pair, and whether participants were asked to select the “Less” or “More” brispy item were randomly shuffled over trials.

The format of the two test phases, *test short* and *test long*, was identical (but different to *train short*). Test phases consisted of 144 trials (2 blocks of 57 trials each) in which lone objects were presented in a random sequence, with the constraint that each combination of 8 (current trial) x 8 (previous trial) ranked objects occurred exactly once in the first half (57 trials) and once in the second half (57 trials) of the test phase. Each object was presented centrally for 750ms, after which participants had 3000ms to respond whether it was more or less brispy than the previous object. Participants were instructed about the mapping from more/less to F/J keys before each block of trials, and this mapping switched midway through the test phase. After the response deadline ended, 3750ms after stimulus onset, there was a pseudo-randomly jittered ITI of 500– 1000ms, during which the fixation dot turned blue if a response was recorded (regardless of accuracy) and a red-letter X appeared if the response was missed. Critically, participants did not receive trial-wise feedback and instead only saw accuracy over the preceding block after the last trial of each block.

The boundary training phase (*train long*) occurred between *test short* and *test long*. It was similar to *train short* except that it lasted just 20 trials, and critically, participants were only trained to compare objects *i_4_* and *i_5_*. Still, as in *train short*, hemifield-presentation of these objects was randomized over trials, as well as whether participants were asked to judge “more” or “less” brispy. As in *train short,* participants had 5000ms to respond and received fully informative feedback after each choice.

After a 200ms fixation delay, two stimuli were presented in each hemifield and participants were instructed to select the corresponding object (i.e. that which was more or less brispy) using either the ‘‘F’’ (left object) or ‘‘J’’ (right object) keys. These objects remained on screen for 5000ms, and once a response was recorded, a green (correct) or red (incorrect) box would appear around the selected object to indicate whether it was the correct selection. During this phase of the experiment, participants responded with average reaction times (RTs) of 1498.3ms ± 37.4 SD; if participants did not respond within 5000ms of stimulus presentation, participants saw a red “x” and the trial was considered incorrect. Either way, feedback persisted for a variable ITI of 500- 1000ms before the next trial. Critically, participants were only trained to compare the 3 consecutive object pairs within each context (*i*_1_–*i*_2_, *i*_2_–*i*_3_ and *i*_3_–*i*_4_ from context A and *i*_5_–*i*_6_, *i*_6_–*i*_7_ and *i*_7_–*i*_8_ from context B). Object hemifield, the trial-order of each object pair, and whether participants were asked to select the “Less” or “More” brispy item were randomly shuffled over trials.

### Debrief and Explicit Knowledge Assessment

Following the experiment, participants completed a free arrangement task to assess explicit knowledge of the relational structure. Participants were presented with eight cards (physical cards in the lab version, digital cards in the online version), each displaying one object from the experiment. They were instructed to arrange the cards by "brispness" in any spatial configuration they preferred, with no constraints on layout. In the lab version, the experimenter observed and recorded arrangement strategies (e.g., sequential pairwise comparisons versus holistic spatial organization) and photographed the final configuration. Participants could rearrange cards at any time before indicating completion. For the online version, participants used a drag-and-drop interface to position digital cards on their screen, with the final arrangement automatically recorded. For both lab and online versions, the (x, y) spatial coordinates of each object’s final position were extracted and used to calculate rotation-invariant Spearman correlations with the ground truth ordering (items ranked from 1-8). Higher correlations indicated better explicit representation of the learned relational structure.

To establish correlation thresholds for classifying participants as assemblers, we conducted Monte Carlo simulations of 8-object spatial arrangements with varying degrees of structure. Simulated arrangements included perfect lines (horizontal, vertical, diagonal), noisy lines (Gaussian noise with σ = 0.1-4.0 added to canonical positions), geometric patterns (grids, circles), clustered configurations (2-4 clusters, σ per cluster = 0.3-0.5), random arrangements, and reversed orderings.

We used a rotation-invariant Spearman correlation metric that computed correlations along six projection axes (x, y, diagonal, anti-diagonal, and cross-correlations) and selected the maximum absolute value. This approach ensured equivalent treatment of vertical, horizontal, and diagonal organizations. We evaluated four potential correlation thresholds for binary classification of participants as assemblers versus non-assemblers: Liberal criterion: *ρ* ≥ 0.5, Moderate criterion: *ρ* ≥ 0.7, Stringent criterion: *ρ* ≥ 0.85, Very stringent criterion: *ρ* ≥ 0.9. We used a rotation-invariant Spearman correlation metric that computed correlations along six projection axes - x-axis, y-axis, main diagonal (x + y), anti-diagonal (x - y), and two cross-axes (x vs y and y vs x). The final correlation coefficient for each arrangement pair was defined as the signed correlation with the maximum absolute value across all six projections. This approach ensures that arrangements organized vertically, horizontally, or diagonally receive equivalent correlation scores, removing arbitrary bias based on spatial orientation.

To characterize the relationship between spatial noise and correlation strength, we generated 100 noisy line arrangements as Position_noisy_ = Position_canonical_ + ε, where ε ∼ N(0, σ²I) represents bivariate Gaussian noise with standard deviation σ. We did this at each of 20 noise levels (σ = 0 to 4.0 in increments of 0.2) and calculated the mean correlation with the canonical line. This mapping allowed us to interpret correlation thresholds in terms of tolerable positional variance, providing an intuitive scale for understanding what different threshold values represent in terms of spatial organization quality.

Spearman correlation was chosen over Pearson correlation because it depends on rank ordering rather than precise Euclidean distances, appropriate for tasks where relational structure matters more than absolute positioning. Simulations were conducted in Python 3.12 using NumPy 1.26.4, SciPy 1.11.4, and Matplotlib 3.8.3. Visual inspection of simulated arrangements indicated that ρ ≥ 0.9 best distinguished clear spatial organization from noisy configurations. Participants with ρ ≥ 0.9 were classified as "Assemblers" (observed *ρ* range: 0.08-0.99).

### Behavioral Analysis

We quantified the number of trials it took for participants to reach criterion during *train short* by extracting performance on trials from context A and context B separately, and then for each context computing a moving average over a moving, preceding 24-trial window (i.e., half of a block of trials). Once participants performed above 90%, and their performance stayed above this 90% threshold for the remainder of trials within that context, we considered that trial to be the trial on which participants learned that context. We used a linear mixed effects model that included a random intercept for the mean accuracy level of each participant to investigate the potential interactive impact of condition and context as well as condition and first context encountered (MATLAB built-in ‘lme’).

SDE slopes were estimated using linear regression (MATLAB built-in “regress”) and then assessed for significance using a t-test against zero. The design matrix for these regression models was constructed according to the ground truth absolute distance between comparisons. Thus, in *test short* SDEs were regressed against vectors that ranged from 0 (between context, same rank; *i*_1_ and *i*_5_) to 3 (*i*_1_ vs *i*_4_ or *i*_1_ vs *i*_8_) and in *test long* ranged from 1 (*i*_1_ and *i*_2_) to 7 (*i*_1_ and *i*_8_).

We analyze behavioral data by plotting accuracies and RTs for each combination of 8 objects shown at test, and/or as a function of symbolic distance (i.e. the distance in rank between the current and previous item). For magnitude, we constructed idealized reaction time and choice matrices under the assumption that choices were noiseless triangular matrices and that RTs depended linearly on symbolic distance. Our accuracy matrices were created by setting the upper triangle of each quadrant (in *test short*) or the entire matrix (in *test long*) to 1 and the lower triangle to zero. In *test short*, this model collapses across context, meaning between-context comparisons are treated as if they came from the same context (e.g. the 3^rd^ item in one set should be ranked higher than the 4^th^ in the other). Thus, this quantifies participant’s tendency to perfectly match rank orderings between contexts (Fig. 2E). Specifically, our idealized RT matrices were constructed as

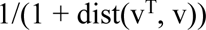

where v = [1:4 1:4] for *test short* and [1:8] for *test long*, and v^T^ is the transpose of v. Context matrices were binary, with 0 for within-context pairs, 1 for between-context. Finally, certainty matrices were quadratic transformation y(x) = 2x² - 1 applied to evenly spaced positions, creating U-shaped profiles ^33^. Note that as participants did not compare objects to themselves, diagonal elements of our design matrices were excluded from analyses. We used competitive multiple regression to isolate the unique variance explained by each model (short and long magnitude, certainty and context RT matrices), with coefficients tested against zero via t-tests. 2-way ANOVAs were used to evaluate the impact of condition and assembly success on *βs* (MATLAB built-in “anovan”)

### EEG System and Setup

EEG data were recorded using a 64-channel BioSemi ActiveTwo EEG system (Amsterdam, The Netherlands), using an online bandpass filter from 100 Hz to 0.16 Hz. The BioSemi system utilizes active electrodes and is designed to provide high signal quality while minimizing common EEG artifacts. Participants were comfortably seated in a dimly lit and sound-attenuated room during the recording sessions. Event triggers were recorded in the EEG data file to mark the time of stimulus onset and subject’s response.

Ag/AgCl active electrodes were arranged in accordance with the 10-20 system, covering frontal, central, parietal, and occipital regions bilaterally. In addition to the 64 scalp electrodes, one reference electrode was placed on each mastoid, and 6 electrodes were placed around the eyes to identify and reject trials with blink and saccade artifacts. Prior to data acquisition, electrode impedance levels were maintained below 20 kΩ for all channels, ensuring optimal signal quality. Impedance was routinely checked during breaks to address any changes in skin-electrode contact. The EEG data were recorded at a sampling rate of 1024 Hz.

### EEG preprocessing and ERPs

Data underwent PREP pipeline preprocessing using standard parameters^83^. PREP detrends the data, applies Slepian tapers using EEGLAB plugin “cleanline” to reduce line noise at 60 Hz, re-references the data to a robust average signal and identifies bad channels (2.88 channels on average for each dataset)^83^. Data were then bandpass filtered between 0.1Hz and 60 Hz using a third order Butterworth filter to attenuate slow drift and 60 Hz line noise and epoched from 750ms to 4000ms peristimulus. PREP then performs independent component analysis (ICA, “runica” method in EEGLAB) to identify additional artifacts^83^. Components reflecting ocular, heartbeat, or muscle artifacts (mean 4.2 ± 3.3 SD per participant) were removed upon visual inspection.

ERPs were computed by averaging stimulus-locked time courses at electrodes of interest (Cz, FCz, PoZ) after trial-wise baseline correction (-200 to 0ms). Cz and FCz were included based on previous results localizing knowledge assembly to fronto-parietal cortex ^9^, and posterior channel PoZ was included to investigate perceptual effects. One and two-way ANOVAs tested effects of training schedule and knowledge assembly, with resulting p-values FDR-corrected from stimulus onset to 1000ms.

### Context decoding

Binary support vector machine (SVM) classifiers were trained predict context identity from data recorded during *train short*, when participants made choices between two simultaneously presented neighboring items within each context. All analyses used MATLAB’s built-in “fitcsvm” function with a polynomial kernel (order 2) and standardization enabled. Polynomial kernels are well-suited for EEG data due to their ability to handle signal nonlinearity, noise, and non-stationarity while maintaining interpretability ^84–86^.

Neural activity was extracted from eight non-overlapping 550ms time windows spanning the trial epoch. Window centers were positioned at -275, 275, 825, 1375, 1925, 2475, 3025, and 3575ms relative to stimulus onset. For each window, voltage was averaged across the 550ms interval for each channel, yielding a single feature vector per trial per time window. We subsequently implemented two complementary decoding approaches. First, we performed within-timepoint decoding by training separate classifiers on data from each of the eight time windows independently. Second, we performed concatenated activity from all windows into a single feature vector. All classifiers used stratified 5-fold cross-validation, ensuring balanced representation of context labels across folds. Classification accuracy was computed as one minus the cross-validated loss returned by MATLAB’s built-in “kfoldLoss” function. Within-participant null distributions were computed by randomly shuffling context labels 50 times and repeating the full decoding procedure on each permutation. This approach controls for potential classification biases arising from data structure or feature distributions.

Each channels contribution to decoding accuracy was quantified using a leave-one-out approach. For each of the 64 channels, we re-trained the across-timepoint decoder using all channels except the target channel. Channel importance was calculated as the difference between baseline accuracy (all channels included) and leave-one-out accuracy (target channel excluded). Higher values indicate greater channel contribution to successful context discrimination. We compared decoding accuracies against theoretical chance (0.5) using one-sample t-tests. We compared empirical accuracies to shuffled baselines using paired t-tests. We assessed effects of learning condition (blocked, alternating, interleaved) and performance group (assemblers, non-assemblers) using repeated-measures ANOVAs with decoding accuracy as the dependent measure. We applied false discovery rate (FDR) correction to control for multiple comparisons across time windows and statistical tests (Benjamini & Hochberg, 1995). Channel importance maps were similarly tested against zero using one-sample t-tests with FDR correction across channels.

### Representational Similarity Analysis

#### Neural RDM Construction

To characterize how neural representations of magnitude structure evolved over time, we computed representational dissimilarity matrices (RDMs) for each participant at each time point using established RSA procedures adapted for EEG data^87,88^. EEG signals from all 64 channels were first z-scored across all trials and time points to normalize signal amplitudes and ensure equal weighting of channels. For each participant, we constructed a 114 trial × 8 stimulus design matrix using one-hot encoding, where each column indexed one of the eight stimuli (four ranks × two contexts). At each time point, we regressed the normalized EEG activity in each channel against this design matrix using ordinary least squares regression, yielding channel-wise *β* coefficients that captured stimulus-evoked responses for each of the eight stimuli.

Neural RDMs were constructed by computing pairwise correlation distances (1 − Pearson’s r) between the *β* patterns for each stimulus pair across the 64-dimensional channel space (Carlson et al., 2013). This yielded an 8 × 8 symmetric RDM at each time point, where smaller values indicated more similar neural representations. Each RDM was then normalized by dividing by its maximum value to facilitate comparison across time points and participants.

To balance temporal resolution with signal stability, RDMs were averaged within 300ms sliding windows (±150ms) centered at time points ranging from −750ms to +3730ms relative to stimulus onset. These windows were spaced at 120ms intervals, resulting in partial temporal overlap (50% overlap between adjacent windows) to capture smooth dynamics of representational change while maintaining adequate trial counts for stable covariance estimates. This approach follows established practices in time-resolved RSA for electrophysiological data (King & Dehaene, 2014; Cichy et al., 2014).

#### Model RDM Construction

We generated five model RDMs to test specific hypotheses about magnitude representations. Model construction followed procedures established in our prior fMRI work (Nelli et al., 2023, *Neuron*). The *test short* magnitude model (RDM_s mag_) captured ordinal distances between item ranks within each context separately (ranks 1-4 for context A; ranks 1-4 for context B), predicting local magnitude structure. This was operationalized as the Euclidean distance matrix for the vector [1, 2, 3, 4, 1, 2, 3, 4]. The *test long* magnitude model (RDM_l mag_) encoded ordinal distances between ranks across the full integrated range (ranks 1-8), representing unified magnitude knowledge. This was operationalized as the Euclidean distance matrix for the vector [1, 2, 3, 4, 5, 6, 7, 8]. The context model (RDM_ctx_) indexed categorical distinctions between contexts independent of magnitude information. This was implemented as a binary matrix with 0s for within-context stimulus pairs and 1s for between-context pairs.

Two additional models captured certainty-based representations of magnitude, following theoretical frameworks linking representational precision to distance from category boundaries ^30,31,33,79,89,89^. These models predict that extreme values (endpoints of a range) are represented with greater certainty than intermediate values, producing characteristic U-shaped dissimilarity patterns where endpoints are maximally distinct from middle values^33^. For the local certainty model (RDM_short cert_), we computed distances between stimuli based on their squared deviation from the midpoint within each context. Specifically, for ranks 1-4 within each context, stimuli were first mapped to positions evenly spaced from −1 to +1. These values were then transformed via a quadratic function y(x) = 2x² − 1, creating a U-shaped profile where extreme values (ranks 1 and 4) received similar high certainty values and intermediate values (ranks 2 and 3) received lower certainty values. Pairwise Euclidean distances between these transformed values formed RDM_short cert_. This transformation was applied independently to each context, yielding the pattern [y(−1), y(−0.33), y(0.33), y(1), y(−1), y(−0.33), y(0.33), y(1)].

The global certainty model (RDM_l cert_) applied the same quadratic transformation across the full integrated range of eight stimuli linearly spaced from −1 to +1, predicting certainty effects at the global scale. This yielded the pattern [y(−1), y(−0.71), y(−0.43), y(−0.14), y(0.14), y(0.43), y(0.71), y(1)]. All model RDMs were normalized by dividing by their maximum value to equate scale, then z-scored across the 28 unique pairwise distances (upper triangle, excluding diagonal) prior to comparison with neural data. This normalization procedure ensures that model comparison reflects pattern similarity rather than magnitude differences (Walther et al., 2016).

#### RSA Statistical Framework

Similarity between neural and model RDMs was quantified using Pearson correlation computed on the z-scored upper triangle of each matrix (28 unique pairwise distances), excluding the diagonal. For each participant and time point, we computed five correlation coefficients, one for each model RDM. To establish when each representational structure emerged, we conducted one-sample t-tests against zero across participants at each time point. Statistical significance was assessed using the Benjamini-Hochberg false discovery rate (FDR) procedure across all time points and models jointly (α = 0.05), controlling for the multiple comparisons inherent in time-resolved analysis (Benjamini & Hochberg, 1995).

To isolate unique variance explained by each representational structure while controlling for shared variance among models, we conducted multiple linear regression at each time point ^87^. For *test short* blocks, neural RDM values (28 distances per participant per time point) were regressed simultaneously on RDM_s mag_, RDM_ctx_, and RDM_s cert_, and resulting regression coefficients (*β*) were extracted for each participant and time point.:

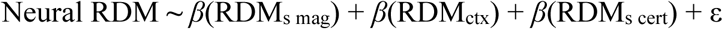

For *test long* blocks, we tested three nested models. Model 1 tested our expected prediction for how neural geometries should rearrange under assembly:

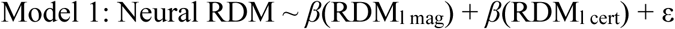

Models 2 tested whether context or certainty from the test short phase impacted assembly.

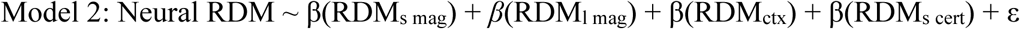

Group-level statistical inference was conducted on *β* coefficients using one-sample t-tests against zero with FDR correction across all time points and model predictors jointly (α = 0.05).

Significant positive coefficients indicate that a model RDM captures unique variance in neural representations beyond that explained by other models.

### Dimensionality Analyses

To quantify the intrinsic complexity of magnitude representations, we computed effective dimensionality of neural RDMs at each time point^90–92^. This metric captures how many independent dimensions are required to account for observed representational geometry. For each participant’s test short and test long RDMs, we applied classical multidimensional scaling (Torgerson, 1952) to the correlation distance matrix. Dimensionality (D_obs_) was defined as the number of principal components (positive eigenvalues) required to explain 80% of the total variance in pairwise distances. This criterion balances sensitivity to meaningful representational structure while excluding noise dimensions ^93^.

To assess statistical significance and control for inherent dimensionality constraints of 8-stimulus RDMs, we implemented a permutation-based null distribution (D_bsln_). For each observed RDM, we generated 10,000 permuted null RDMs by randomly shuffling the 28 unique dissimilarity values (upper triangle) while preserving matrix symmetry and zeros along the diagonal. For each permutation, we computed dimensionality using the same 80% variance criterion. The effective dimensionality for each participant at each time point was defined as the difference between observed dimensionality and the mean permuted dimensionality:

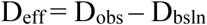

This isolates representational structure beyond chance expectations given the number of stimuli. Statistical significance was assessed using one-sample t-tests on effective dimensionality across participants, with FDR correction across time points.

### Statistics

All statistical tests employed two-sided hypothesis testing unless otherwise noted. For analyses involving time series data, we controlled the false discovery rate at α = 0.05 using the Benjamini-Hochberg procedure (Benjamini & Hochberg, 1995). In competitive regression analyses where multiple model predictors were tested simultaneously, FDR correction was applied across all time points and model regressors jointly to maintain appropriate control of the family-wise error rate. The corrected significance threshold (pFDR) is reported where applicable.

To examine the influence of curriculum condition (blocked, interleaved, baseline) and knowledge assembly performance (assemblers vs. non-assemblers) on neural representations, we conducted two-way repeated-measures ANOVAs with curriculum and performance group as between-subjects factors at each time point. For RSA measures (correlations, regression coefficients) and dimensionality metrics, ANOVA was applied to participant-level estimates. Main effects and interactions were assessed with FDR correction across time points.

Group-level inference for all RSA and dimensionality analyses followed a random-effects framework. Statistics were computed on individual participant estimates (correlation coefficients, regression weights, or dimensionality scores) using one-sample t-tests against zero, ensuring generalizability to the broader population. Effect sizes are reported for t-tests, F-values and as standardized regression coefficients (*β*) for multiple regression models. Sample sizes were determined based on power analyses from our previous fMRI investigation of magnitude knowledge assembly, adjusted for the temporal resolution afforded by EEG.

### Online Study

To replicate the behavioral results in a larger sample, the experiment was adapted from MATLAB to JavaScript using jsPsych v8.0.2 and hosted on a web server for online data collection (de Leeuw et al., 2023). We recruited 174 participants through the crowdsourcing platform Prolific. Inclusion criteria required participants to be between 18 and 40 years of age, fluent in English, located in the United States, Canada, or the United Kingdom, report no neurological or ocular conditions, and have a Prolific approval rate of at least 95%. Prior to participation, all individuals provided informed consent and confirmed their understanding of the study procedures, data handling policies, as well as their right to withdraw from the study at any time. Twenty-nine participants were screened out after ten blocks of training for failing to reach a criterion of at least 90% accuracy in each context, and five participants were excluded due to technical difficulties with their browsers; these participants received prorated compensation ($3.50 or $0.14/minute, whichever was higher). The final sample consisted of 140 participants (mean age = 28.64 ± 5.87 years, range 19–40; 49.3% female, 45.7% male, and 5.0% nonbinary). All participants who completed the entire study received a base compensation of $10 and a bonus payment of up to $5 for exemplary performance. The experimental phases and trial structure proceeded in an identical manner to the main experiment.

Following the test phase, participants completed a free arrangement task implemented as a drag-and-drop interface. All eight object images were displayed simultaneously on screen, and participants were instructed to arrange them by "brispness" in whatever spatial configuration they preferred within a bounded 2D workspace. Participants could freely reposition objects at any time before submitting their final arrangement. Spatial coordinates were recorded and analyzed using the same correlation-based criteria as the in-lab study to identify Assemblers (*r* ≥ 0.9 with ground truth structure).

### Recurrent neural network

#### Network Architecture

We implemented vanilla recurrent neural networks to test whether simple associative learning mechanisms could reproduce human behavioral and neural patterns during knowledge assembly. The network architecture consisted of three layers: an input layer with 8 units (one-hot encoding of each stimulus), a recurrent hidden layer with 64 units, and an output layer with 2 units (corresponding to "more" or "less" responses). The hidden layer used tanh activation functions, while the output layer employed softmax activation to produce response probabilities.

The hidden state at time *t* was computed as:

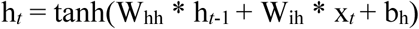

where W_hh_ represents recurrent weights, W_ih_ represents input-to-hidden weights, and b_h_ is the bias term. Output was computed as:

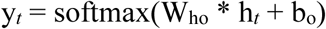

where W_ho_ represents hidden-to-output weights and b_o_ is the output bias.

#### Training Procedure

Networks underwent training that paralleled the human experimental design. Training consisted of a train short phase where networks learned two independent transitive orderings (items 1-4 and 5-8) through adjacent pair comparisons. Networks were trained under either blocked (N = 10) or interleaved (N = 10) curriculum schedules. In the blocked condition, all trials from context A were presented consecutively, followed by all trials from context B. In the interleaved condition, trials from both contexts were randomly intermixed within each training epoch.

Each network was trained using backpropagation through time with cross-entropy loss. We used the Adam optimizer with a learning rate of 0.001 and trained for a maximum of 100 epochs.

Networks advanced to the test short phase once they achieved at least 90% accuracy on both contexts. Following test short, networks received boundary training (train long) consisting of 20 trials comparing items 4 and 5, after which they were evaluated during test long.

The task structure mirrored the human experiment. On each trial, the network received a one-hot encoded input representing a single stimulus and produced a binary classification of whether that stimulus was "more" or "less" than the previous stimulus. Ground truth labels were determined by ordinal relationships within each context during train short and test short, and according to the integrated 8-item ordering during test long.

#### Behavioral and Representational Analyses

Network performance was quantified through accuracy on within-context and between-context comparisons during test long. We computed symbolic distance effects by regressing accuracy against ordinal distance between compared items. Choice matrices (8×8) were constructed showing the probability of selecting each response for all possible stimulus pairs.

We quantified hidden layer activations using representational similarity analysis. For each stimulus presentation, we extracted the 64-dimensional hidden state vector. Neural RDMs were constructed by computing pairwise correlation distances between stimulus-evoked activity patterns across the 64 hidden units, yielding an 8×8 symmetric matrix at each trial.

### Software and Dependencies

Data were collected using BioSemi ActiveTwo EEG system software, Matlab 2022a and Psychtoolbox-3 were used to collect human EEG data in this study. jsPsych v8.0.2 hosted on web server for online study. Python v3.12.8 with NumPy 1.26.4, SciPy 1.11.4, Matplotlib 3.8.3 were used to collect neural network data in this study.

All analyses were conducted using Matlab 2022a-2025a and Eeglab 2022.0-2025.0 with PREP pipeline and cleanline plugin. Installation typically requires 10 minutes on a standard desktop computer. Detailed installation instructions and system requirements are provided in the README file in the code repository.

Python v3.12.8 with NumPy 1.26.4, SciPy 1.11.4, Matplotlib 3.8.3 were used to analyze neural network data in this study. Installation typically requires 5-10 minutes on a standard desktop computer, with a simulation run-time of approximately 30 minutes per training phase (full replication of 20 models takes approximately 10 hours). Detailed installation instructions and system requirements are provided in the README file in the code repository.

## Data availability

Data that support the findings detailed in Figures 3 and 5 of this study are available at the Open Science Framework (https://osf.io/wkx5h/). Data regarding recurrent neural network simulations is available at https://github.com/CPF2002/Knowledge-Assembly-RNN. Additional data are available from the corresponding author upon reasonable request.

## Code availability

Custom code for recurrent neural network simulations (modified from https://doi.org/10.1016/j.neuron.2021.02.004) is available at https://github.com/CPF2002/Knowledge-Assembly-RNN. Custom MATLAB scripts for neural representational dissimilarity matrix (RDM) computation and analysis are available at https://osf.io/wkx5h/. Additional analysis scripts are available from the corresponding author upon reasonable request.

## Notes

https://github.com/CPF2002/Knowledge-Assembly-RNN.

### Competing Interest Statement

The authors have declared no competing interest.

### Summary of Updates

Small wording changes for clarity in the behavioral results for the initial training phase.

## Works Cited

1. Whittington, J. C. R. et al. The Tolman-Eichenbaum Machine: Unifying Space and Relational Memory through Generalization in the Hippocampal Formation. Cell 183, 1249–1263.e23 (2020).

2. Schapiro, A. C., Turk-Browne, N. B., Norman, K. A. & Botvinick, M. M. Statistical learning of temporal community structure in the hippocampus: *STATISTICAL LEARNING OF TEMPORAL COMMUNITY STRUCTURE*. Hippocampus 26, 3–8 (2016).

3. Schapiro, A. C., McDevitt, E. A., Rogers, T. T., Mednick, S. C. & Norman, K. A. Human hippocampal replay during rest prioritizes weakly learned information and predicts memory performance. Nat Commun 9, 3920 (2018).

4. LaRocque, K. F. et al. Global Similarity and Pattern Separation in the Human Medial Temporal Lobe Predict Subsequent Memory. J. Neurosci. 33, 5466–5474 (2013).

5. Kragel, J. E., Schuele, S., VanHaerents, S., Rosenow, J. M. & Voss, J. L. Rapid coordination of effective learning by the human hippocampus. Sci. Adv. 7, eabf7144 (2021).

6. Kumaran, D., Hassabis, D. & McClelland, J. L. What Learning Systems do Intelligent Agents Need? Complementary Learning Systems Theory Updated. Trends in Cognitive Sciences 20, 512–534 (2016).

7. McClelland, J. L., McNaughton, B. L. & O’Reilly, R. C. Why there are complementary learning systems in the hippocampus and neocortex: Insights from the successes and failures of connectionist models of learning and memory. Psychological Review 102, 419–457 (1995).

8. Squire, L. R., Genzel, L., Wixted, J. T. & Morris, R. G. Memory Consolidation. Cold Spring Harb Perspect Biol 7, a021766 (2015).

9. Nelli, S., Braun, L., Dumbalska, T., Saxe, A. & Summerfield, C. Neural knowledge assembly in humans and neural networks. Neuron S0896627323001186 (2023) doi:10.1016/j.neuron.2023.02.014.

10. Constantinescu, A. O., O’Reilly, J. X. & Behrens, T. E. J. Organizing conceptual knowledge in humans with a gridlike code. Science 352, 1464–1468 (2016).

11. Summerfield, C., Luyckx, F. & Sheahan, H. Structure learning and the posterior parietal cortex. Progress in Neurobiology 184, 101717 (2020).

12. Vendetti, M. S. & Bunge, S. A. Evolutionary and Developmental Changes in the Lateral Frontoparietal Network: A Little Goes a Long Way for Higher-Level Cognition. Neuron 84, 906–917 (2014).

13. Wendelken, C. & Bunge, S. A. Transitive Inference: Distinct Contributions of Rostrolateral Prefrontal Cortex and the Hippocampus. Journal of Cognitive Neuroscience 22, 837–847 (2010).

14. Bernardi, S. et al. The Geometry of Abstraction in the Hippocampus and Prefrontal Cortex. Cell 183, 954–967.e21 (2020).

15. Theves, S., Neville, D. A., Fernández, G. & Doeller, C. F. Learning and Representation of Hierarchical Concepts in Hippocampus and Prefrontal Cortex. J. Neurosci. 41, 7675–7686 (2021).

16. McCloskey, M. & Cohen, N. J. Catastrophic Interference in Connectionist Networks: The Sequential Learning Problem. in Psychology of Learning and Motivation vol. 24 109–165 (Elsevier, 1989).

17. Grutzendler, J., Kasthuri, N. & Gan, W.-B. Long-term dendritic spine stability in the adult cortex. Nature 420, 812–816 (2002).

18. Kirkpatrick, J., et al. Overcoming catastrophic forgetting in neural networks. Proc. Natl. Acad. Sci. U.S.A. 114, 3521–3526 (2017).

19. Yang, G., Pan, F. & Gan, W.-B. Stably maintained dendritic spines are associated with lifelong memories. Nature 462, 920–924 (2009).

20. Zenke, F., Poole, B. & Ganguli, S. Continual Learning Through Synaptic Intelligence. In Proceedings of the 34 th I (Sydney, Australia, 2017).

21. Abati, D., et al. Conditional Channel Gated Networks for Task-Aware Continual Learning. Preprint at 10.48550/ARXIV.2004.00070 (2020).

22. Aoi, M. C., Mante, V. & Pillow, J. W. Prefrontal cortex exhibits multidimensional dynamic encoding during decision-making. Nat Neurosci 23, 1410–1420 (2020).

23. Flesch, T., Juechems, K., Dumbalska, T., Saxe, A. & Summerfield, C. Orthogonal representations for robust context-dependent task performance in brains and neural networks. Neuron 110, 1258–1270.e11 (2022).

24. Flesch, T., Saxe, A. & Summerfield, C. Continual task learning in natural and artificial agents. Trends in Neurosciences 46, 199–210 (2023).

25. Masse, N. Y., Grant, G. D. & Freedman, D. J. Alleviating catastrophic forgetting using context-dependent gating and synaptic stabilization. Proc. Natl. Acad. Sci. U.S.A. 115, (2018).

26. Tilley, M. J., Miller, M. & Freedman, D. J. Artificial Neuronal Ensembles with Learned Context Dependent Gating. 10.48550/ARXIV.2301.07187 (2023) doi:10.48550/ARXIV.2301.07187.

27. Zeng, G., Chen, Y., Cui, B. & Yu, S. Continual learning of context-dependent processing in neural networks. Nat Mach Intell 1, 364–372 (2019).

28. Brunamonti, E. et al. Neuronal Modulation in the Prefrontal Cortex in a Transitive Inference Task: Evidence of Neuronal Correlates of Mental Schema Management. J. Neurosci. 36, 1223–1236 (2016).

29. Merritt, D. J. & Terrace, H. S. Mechanisms of inferential order judgments in humans (Homo sapiens) and rhesus monkeys (Macaca mulatta). Journal of Comparative Psychology 125, 227–238 (2011).

30. Okazawa, G., Hatch, C. E., Mancoo, A., Machens, C. K. & Kiani, R. Representational geometry of perceptual decisions in the monkey parietal cortex. Cell 184, 3748–3761.e18 (2021).

31. Summerfield, C., Luyckx, F. & Sheahan, H. Structure learning and the posterior parietal cortex. Progress in Neurobiology 184, 101717 (2020).

32. Aru, J., Drüke, M., Pikamäe, J. & Larkum, M. E. Mental navigation and the neural mechanisms of insight. Trends in Neurosciences 46, 100–109 (2023).

33. Yang, Y. & Maass, W. Neurons have an inherent capability to learn order relations: A theoretical foundation that explains numerous experimental data. Preprint at 10.1101/2025.03.17.642834 (2025).

34. Beukers, A. O. et al. Blocked training facilitates learning of multiple schemas. Commun Psychol 2, 28 (2024).

35. Carvalho, P. F. & Goldstone, R. L. Effects of interleaved and blocked study on delayed test of category learning generalization. Front. Psychol. 5, (2014).

36. Carvalho, P. F. & Goldstone, R. L. The benefits of interleaved and blocked study: Different tasks benefit from different schedules of study. Psychon Bull Rev 22, 281–288 (2015).

37. Hamid, O. H., Wendemuth, A. & Braun, J. Temporal context and conditional associative learning. BMC Neurosci 11, 45 (2010).

38. Carvalho, P. F. & Goldstone, R. L. The sequence of study changes what information is attended to, encoded, and remembered during category learning. Journal of Experimental Psychology: Learning, Memory, and Cognition 43, 1699–1719 (2017).

39. Hayes, W. M. & Wedell, D. H. Effects of blocked versus interleaved training on relative value learning. Psychon Bull Rev 30, 1895–1907 (2023).

40. Schlichting, M. L., Mumford, J. A. & Preston, A. R. Learning-related representational changes reveal dissociable integration and separation signatures in the hippocampus and prefrontal cortex. Nat Commun 6, 8151 (2015).

41. Schorn, J. M. & Knowlton, B. J. Interleaved practice benefits implicit sequence learning and transfer. Mem Cogn 49, 1436–1452 (2021).

42. Chen, O., Paas, F. & Sweller, J. Spacing and Interleaving Effects Require Distinct Theoretical Bases: a Systematic Review Testing the Cognitive Load and Discriminative-Contrast Hypotheses. Educ Psychol Rev 33, 1499–1522 (2021).

43. Zhou, Z., Singh, D., Tandoc, M. C. & Schapiro, A. C. Building integrated representations through interleaved learning. Journal of Experimental Psychology: General 152, 2666–2684 (2023).

44. Bowman, C. R. & Zeithamova, D. Training set coherence and set size effects on concept generalization and recognition. Journal of Experimental Psychology: Learning, Memory, and Cognition 46, 1442–1464 (2020).

45. Birnbaum, M. S., Kornell, N., Bjork, E. L. & Bjork, R. A. Why interleaving enhances inductive learning: The roles of discrimination and retrieval. Mem Cogn 41, 392–402 (2013).

46. Kornell, N. & Bjork, R. A. Learning Concepts and Categories: Is Spacing the “Enemy of Induction”? Psychol Sci 19, 585–592 (2008).

47. Brem, S. et al. Increasing expertise to a novel script modulates the visual N1 ERP in healthy adults. International Journal of Behavioral Development 42, 333–341 (2018).

48. Kimura, M. Prediction, Suppression of Visual Response, and Modulation of Visual Perception: Insights From Visual Evoked Potentials and Representational Momentum. Front. Hum. Neurosci. 15, 730962 (2021).

49. Nieuwland, M. S. Do ‘early’ brain responses reveal word form prediction during language comprehension? A critical review. Neuroscience & Biobehavioral Reviews 96, 367–400 (2019).

50. Curran, T., Tucker, D. M., Kutas, M. & Posner, M. I. Topography of the N400: brain electrical activity reflecting semantic expectancy. Electroencephalography and Clinical Neurophysiology/Evoked Potentials Section 88, 188–209 (1993).

51. Kuperberg, G. R. Separate streams or probabilistic inference? What the N400 can tell us about the comprehension of events. Language, Cognition and Neuroscience 31, 602–616 (2016).

52. Lindborg, A., Musiolek, L., Ostwald, D. & Rabovsky, M. Semantic surprise predicts the N400 brain potential. Neuroimage: Reports 3, 100161 (2023).

53. Brothers, T. & Wlotko, E. W. A Tale of Two Positivities and the N400: Distinct Neural Signatures Are Evoked by Confirmed and Violated Predictions at Different Levels of Representation. Journal of Cognitive Neuroscience 32, 12–35 (2020).

54. Zhao, Y., Guo, J., Li, Y., Wu, Y. & Luo, J. ERP evidence for temporal differences between cross-modal and cross-domain analogical reasoning. Behavioural Brain Research 470, 115072 (2024).

55. Wronka, E., Kaiser, J. & Coenen, A. Neural generators of the auditory evoked potential components P3a and P3b. Acta Neurobiol. Exp. 72, 51–64 (2012).

56. Polich, J. & Criado, J. R. Neuropsychology and neuropharmacology of P3a and P3b. International Journal of Psychophysiology 60, 172–185 (2006).

57. Verleger, R. Effects of relevance and response frequency on P3b amplitudes: Review of findings and comparison of hypotheses about the process reflected by P3b. Psychophysiology 57, e13542 (2020).

58. Kikumoto, A., Bhandari, A., Shibata, K. & Badre, D. A transient high-dimensional geometry affords stable conjunctive subspaces for efficient action selection. Nat Commun 15, 8513 (2024).

59. Di Antonio, G., Raglio, S. & Mattia, M. A geometrical solution underlies general neural principle for serial ordering. Nat Commun 15, 8238 (2024).

60. Sheahan, H., Luyckx, F., Nelli, S., Teupe, C. & Summerfield, C. Neural state space alignment for magnitude generalization in humans and recurrent networks. Neuron 109, 1214–1226.e8 (2021).

61. French, R. Catastrophic forgetting in connectionist networks. Trends in Cognitive Sciences 3, 128–135 (1999).

62. McCloskey, M. & Cohen, N. J. Catastrophic Interference in Connectionist Networks: The Sequential Learning Problem. in Psychology of Learning and Motivation vol. 24 109–165 (Elsevier, 1989).

63. Pisupati, S. & Niv, Y. The challenges of lifelong learning in biological and artificial systems. Trends in Cognitive Sciences 26, 1051–1053 (2022).

64. Ratcliff, R. Connectionist Models of Recognition Memory: Constraints Imposed by Learning and Forgetting Functions. Psychological Review 97, 285–308 (1990).

65. Jehee, J. F. M., Ling, S., Swisher, J. D., Van Bergen, R. S. & Tong, F. Perceptual Learning Selectively Refines Orientation Representations in Early Visual Cortex. J. Neurosci. 32, 16747–16753 (2012).

66. Lövdén, M., Wenger, E., Mårtensson, J., Lindenberger, U. & Bäckman, L. Structural brain plasticity in adult learning and development. Neuroscience & Biobehavioral Reviews 37, 2296–2310 (2013).

67. Polich, J. Updating P300: An integrative theory of P3a and P3b. Clinical Neurophysiology 118, 2128–2148 (2007).

68. Meyniel, F. Brain dynamics for confidence-weighted learning. PLoS Comput Biol 16, e1007935 (2020).

69. Payzan-LeNestour, E. & Bossaerts, P. Risk, Unexpected Uncertainty, and Estimation Uncertainty: Bayesian Learning in Unstable Settings. PLoS Comput Biol 7, e1001048 (2011).

70. Meyniel, F. & Dehaene, S. Brain networks for confidence weighting and hierarchical inference during probabilistic learning. Proc. Natl. Acad. Sci. U.S.A. 114, (2017).

71. Nassar, M. R., Wilson, R. C., Heasly, B. & Gold, J. I. An Approximately Bayesian Delta-Rule Model Explains the Dynamics of Belief Updating in a Changing Environment. J. Neurosci. 30, 12366–12378 (2010).

72. Boldt, A. & Gilbert, S. J. Partially Overlapping Neural Correlates of Metacognitive Monitoring and Metacognitive Control. J. Neurosci. 42, 3622–3635 (2022).

73. Rahnev, D. & Fleming, S. M. How experimental procedures influence estimates of metacognitive ability. Neuroscience of Consciousness 2019, niz009 (2019).

74. Gerstner, W., Lehmann, M., Liakoni, V., Corneil, D. & Brea, J. Eligibility Traces and Plasticity on Behavioral Time Scales: Experimental Support of NeoHebbian Three-Factor Learning Rules. Front. Neural Circuits 12, 53 (2018).

75. Piray, P. & Daw, N. D. A model for learning based on the joint estimation of stochasticity and volatility. Nat Commun 12, 6587 (2021).

76. Gick, M. L. & Holyoak, K. J. Schema induction and analogical transfer. Cognitive Psychology 15, 1–38 (1983).

77. Shadlen, M. N. & Kiani, R. Decision Making as a Window on Cognition. Neuron 80, 791– 806 (2013).

78. Ma, W. J. & Jazayeri, M. Neural Coding of Uncertainty and Probability. Annu. Rev. Neurosci. 37, 205–220 (2014).

79. Kiani, R., Corthell, L. & Shadlen, M. N. Choice Certainty Is Informed by Both Evidence and Decision Time. Neuron 84, 1329–1342 (2014).

80. D’Amato, M. R. & Colombo, M. The symbolic distance effect in monkeys (Cebus apella). Animal Learning & Behavior 18, 133–140 (1990).

81. Van Opstal, F., Gevers, W., De Moor, W. & Verguts, T. Dissecting the symbolic distance effect: Comparison and priming effects in numerical and nonnumerical orders. Psychonomic Bulletin & Review 15, 419–425 (2008).

82. Hinton, E. C., Dymond, S., Von Hecker, U. & Evans, C. J. Neural correlates of relational reasoning and the symbolic distance effect: involvement of parietal cortex. Neuroscience 168, 138–148 (2010).

83. Bigdely-Shamlo, N., Mullen, T., Kothe, C., Su, K.-M. & Robbins, K. A. The PREP pipeline: standardized preprocessing for large-scale EEG analysis. Front. Neuroinform. 9, (2015).

84. Li, X., Chen, X., Yan, Y., Wei, W. & Wang, Z. Classification of EEG Signals Using a Multiple Kernel Learning Support Vector Machine. Sensors 14, 12784–12802 (2014).

85. Tong, H. A Note on Support Vector Machines with Polynomial Kernels. Neural Computation 28, 71–88 (2016).

86. Zhang, M., Treder, M., Marshall, D. & Li, Y. Explaining the predictions of kernel SVM models for neuroimaging data analysis. Expert Systems with Applications 251, 123993 (2024).

87. Nili, H. et al. A Toolbox for Representational Similarity Analysis. PLoS Comput Biol 10, e1003553 (2014).

88. Kriegeskorte, N. Representational similarity analysis – connecting the branches of systems neuroscience. Front. Sys. Neurosci. 10.3389/neuro.06.004.2008 (2008) doi:10.3389/neuro.06.004.2008.

89. Zylberberg, A. & Shadlen, M. N. A population representation of the confidence in a decision in the parietal cortex. Cell Reports 44, 115526 (2025).

90. Jazayeri, M. & Ostojic, S. Interpreting neural computations by examining intrinsic and embedding dimensionality of neural activity. Current Opinion in Neurobiology 70, 113–120 (2021).

91. Stringer, C., Pachitariu, M., Steinmetz, N., Carandini, M. & Harris, K. D. High-dimensional geometry of population responses in visual cortex. Nature 571, 361–365 (2019).

92. Tang, E. et al. Effective learning is accompanied by high-dimensional and efficient representations of neural activity. Nat Neurosci 22, 1000–1009 (2019).

93. Rigotti, M. et al. The importance of mixed selectivity in complex cognitive tasks. Nature 497, 585–590 (2013).

