## supplementary for "Training constrains neural routes to knowledge assembly"

Affiliations:

### Supplementary Figures

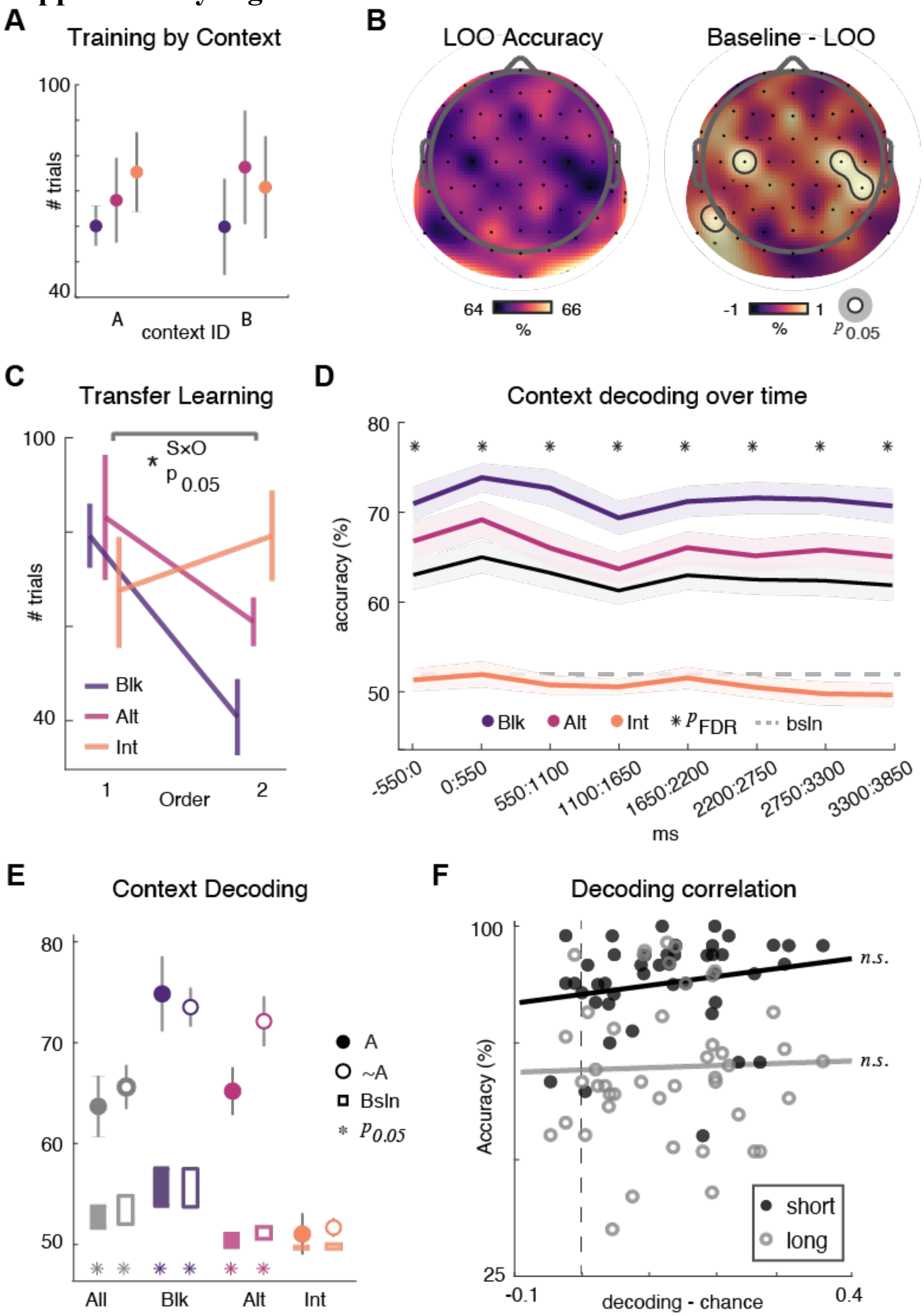


**Supplementary Figure 1: Neural context representations during curriculum learning (A)** Training performance across trials for blocked (purple), alternating (pink), and interleaved (orange) curriculum conditions, shown separately for context A and context B. Points represent mean ± standard error across trials. **(B)** Right: the topographic map of channels contributing to context decoding, as determined by leave-one-out (LOO) decoding accuracy. Channels reaching significance at alpha level 0.05 are outlined with a gray line, no channels significant after correction for multiple comparisons. **(C)** The number of training trials required to reach criterion for participants in each of the three training schedules. Data is shown separated out by the order in which participants encountered the context, irrespective of context identity. There was an interaction, but no main effects, between order and schedule on number of training trials required to learn. Vertical lines indicate standard deviation. **(D)** Context decoding accuracy over the course of *train short* trials. Figure shows temporal dynamics of decoding performance over all participants (gray) as well as for blocked (Blk, purple), alternating (Alt, pink), and interleaved (Int, orange) conditions. Dashed line shows average within-participant baseline decoding over all participants (bsln, dashed red). Asterisks indicate significant timepoints as compared with individual baseline over all participants ($P < 0.05$). Shaded regions represent standard error. **(E)** Context decoding accuracy over all timepoints in the *train short* trial. Figure shows decoding performance for all participants (gray) as well as for blocked (Blk, purple), alternating (Alt, pink), and interleaved (Int, orange). Data is separated out by participants that would go onto assemble in *test long* (A, filled) and those that would not (~A, open). Error bars represent standard error. Rectangles indicate average and standard error of within participant baselines. Asterisks indicate significant differences from within participant baselines. **(F)** Relationship between context decoding accuracy and behavioral performance during test. Figure shows a non-significant trend relating decoding to accuracy. Filled circles represent short test trials; open circles represent long test trials. Black and gray lines indicate linear fits for test short and test long accuracy, respectively. The number of training trials required to reach criterion for participants in each of the three training schedules. Data is shown separated out by the order in which participants encountered the context, irrespective of context identity. There was an interaction, but no main effects, between order and schedule on number of training trials required to learn. Vertical lines indicate standard deviation.

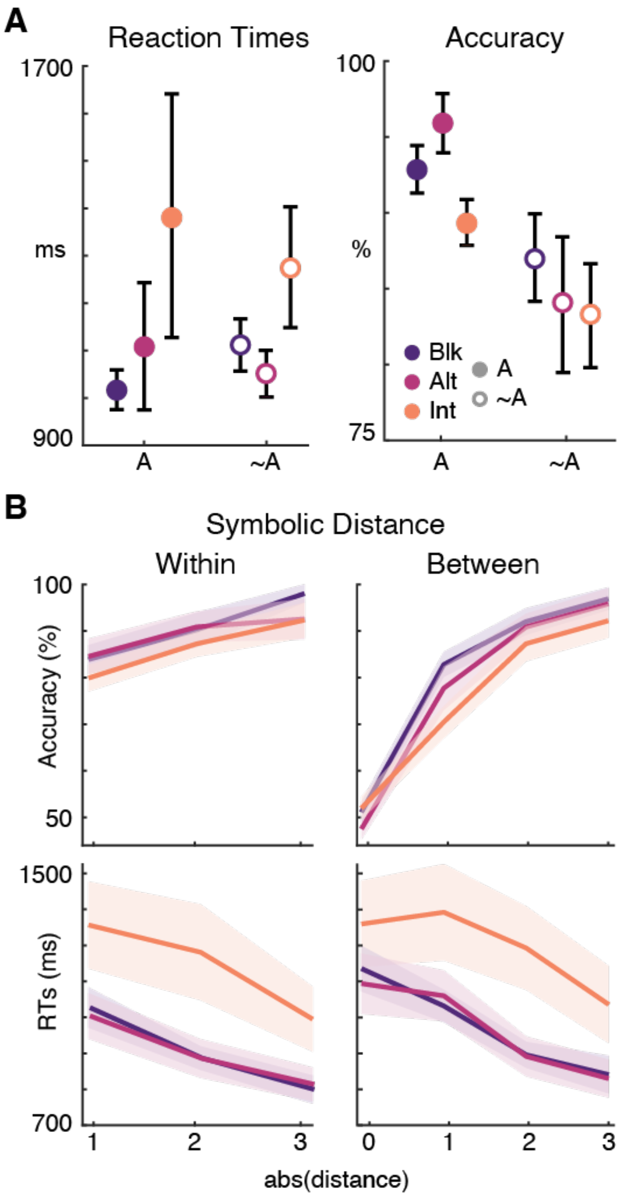


**Supplementary Figure 2: Test short performance differences reveal distinct symbolic distance effects across training schedules. (A)** Reaction times (left) and accuracy (right) during the test short phase for participants in blocked (purple), alternating (pink), and interleaved (orange) training conditions. Data are separated by participants who subsequently assembled knowledge structures in the test long phase (A) versus those who did not (~A). Error bars represent standard error of the mean. **(B)** Symbolic distance effects for comparisons within the same context (left) and between different contexts (right). Top panels show accuracy as a function of absolute symbolic distance between items. Bottom panels show corresponding reaction times. The symbolic distance effect reflects improved performance when comparing items farther apart in the learned relational structure. Shaded regions indicate standard error of the mean. All three training schedules show characteristic distance effects, with accuracy increasing and reaction times decreasing as distance grows, though the magnitude and generalization of these effects differ by condition.

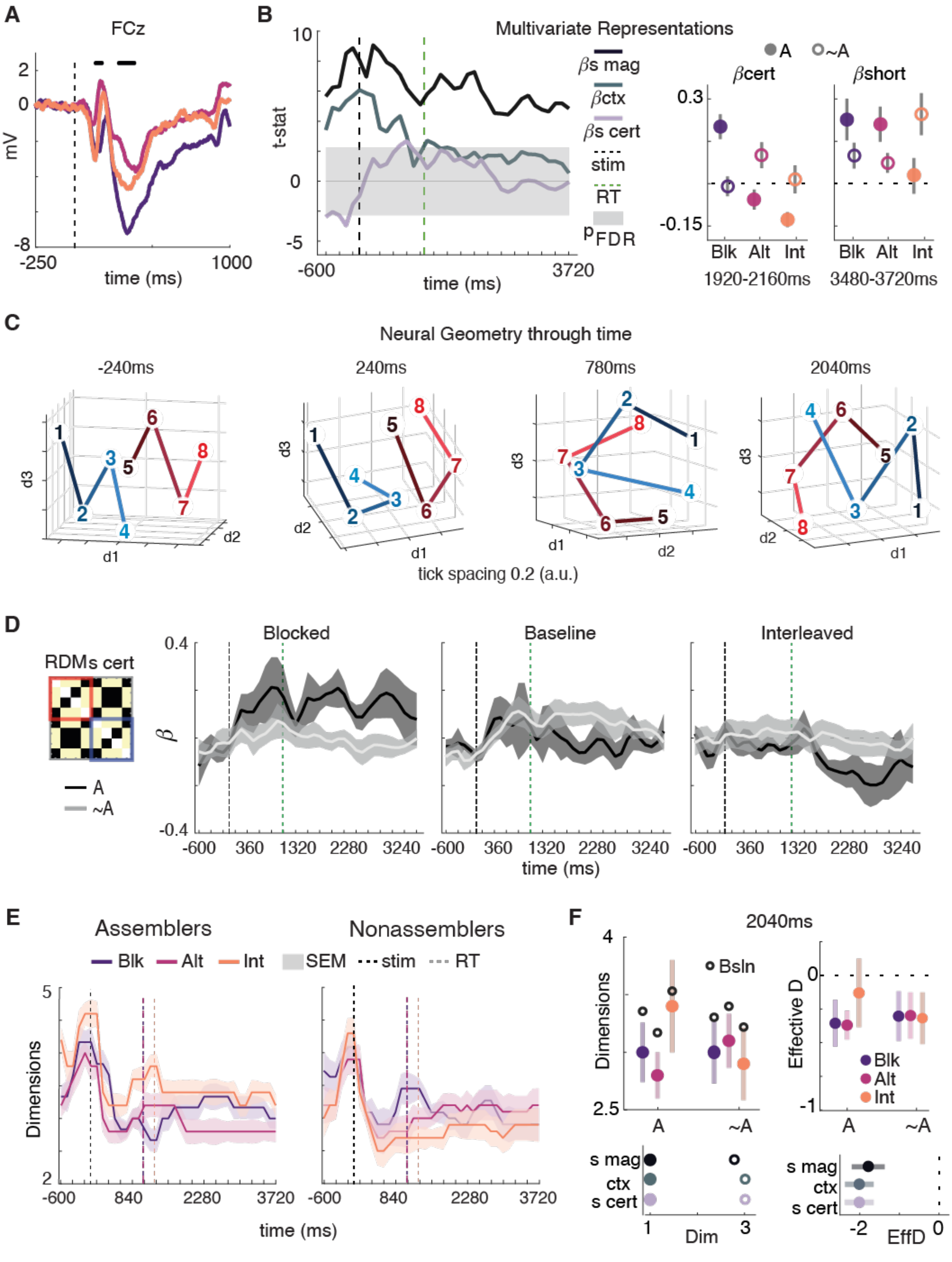


**Supplementary Figure 3: Curriculum learning differentially shapes neural knowledge representations and dimensionality dynamics. (A)** Event-related potential at electrode FCz showing the main effect of curriculum condition. Black dots indicate time points surviving multiple comparison correction at alpha = 0.05. Traces show responses for blocked (purple), alternating (pink), and interleaved (orange) conditions. Dotted vertical lines indicates stimulus onset. **(B)** Left: Multivariate representational similarity analysis testing competitive regression models including short magnitude (βs mag), context (βctx), and short certainty (βs cert) regressors against neural data. Black trace shows t-statistics over time; shaded region indicates non-significance after FDR correction. Dashed vertical lines mark stimulus onset (black) and response time (green). Right: Average beta weights for short certainty and short magnitude regressors across two time windows (1920-2160ms and 3480-3720ms) for blocked (Blk, purple), alternating (Alt, pink), and interleaved (Int, orange) conditions. Filled circles represent participants who subsequently assembled knowledge (assemblers); open circles represent those who did not (nonassemblers). **(C)** Neural geometry trajectories visualized in three-dimensional representational space at four time points (-240ms, 240ms, 780ms, 2040ms). Numbered vertices represent eight stimulus conditions; blue and red traces distinguish the two task contexts. Tick spacing: 0.2 arbitrary units. **(D)** Time-resolved certainty estimates (β values) for blocked, baseline, and interleaved conditions. Inset shows example representational dissimilarity matrix (RDM) structure for certainty predictions. Dark traces represent future assemblers (A); light traces represent future nonassemblers (~A). Shaded regions indicate SEM. Vertical dashed lines mark stimulus onset (black) and mean response time (green). **(E)** Dimensionality of neural representations over time for each curriculum condition (blocked in purple, alternating in pink, interleaved in orange). Left panel shows assemblers; right panel shows nonassemblers. Shaded region indicate standard error (SEM) and dotted lines indicate stimulus onset (stim) and response time (RT). **(F)** Left: Dimensionality at 2040ms for each condition and participant type (filled circles with shaded standard deviations) compared to within-participant noise baseline dimensionality (open circles). Right: Effective dimensionality (EffD) for each condition. Bottom panels show dimensionality (left) and effective dimensionality (right) for idealized representational matrices corresponding to short magnitude (s mag), context (ctx), and short certainty (s cert). Horizontal dashed line at 0 indicates dimensionality equivalent to the noise baseline.

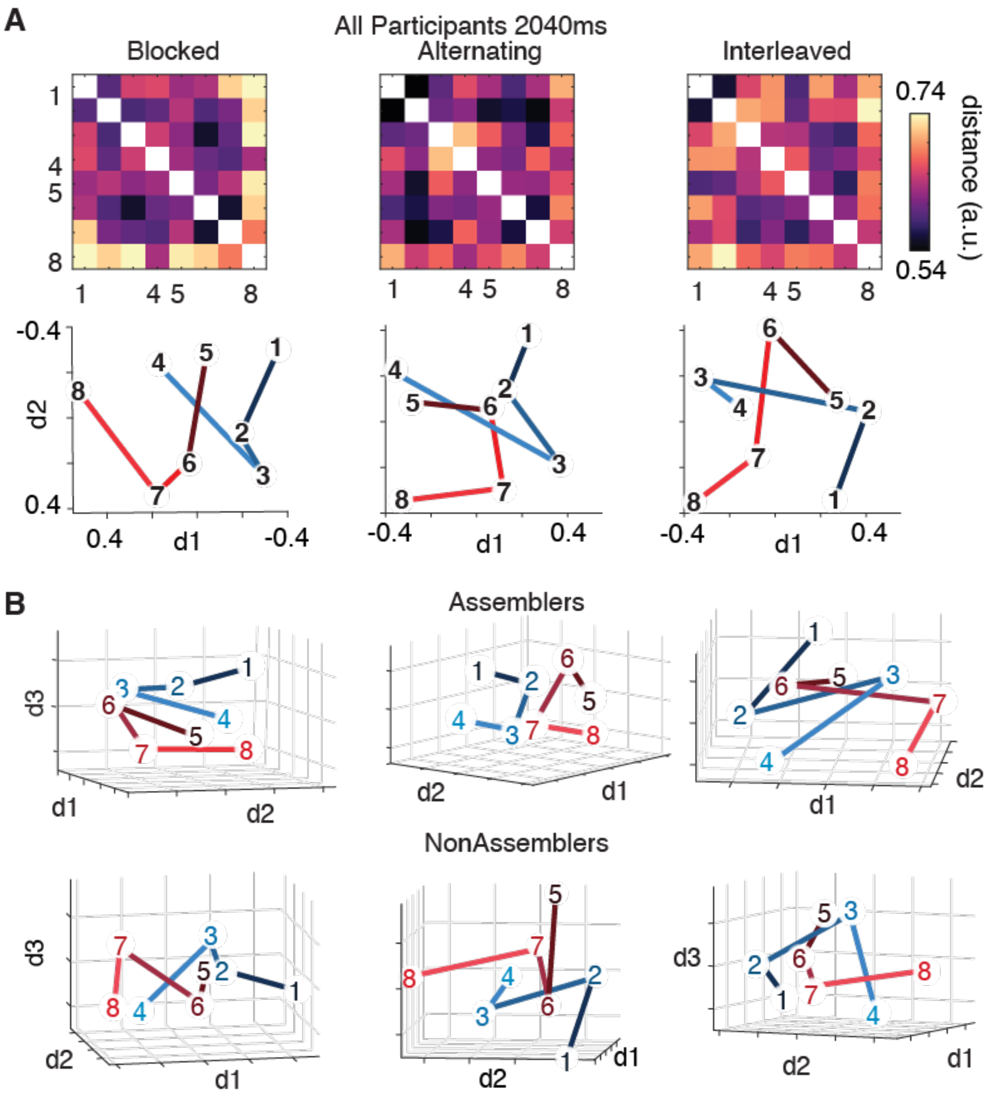


**Supplementary Figure 4: Neural geometries at 2040ms differs across curriculum conditions and between assemblers and nonassemblers. (A)** Neural representational structure averaged across all participants at 2040ms for blocked (left), alternating (middle), and interleaved (right) curriculum conditions. Top row shows representational dissimilarity matrices (RDMs) for the eight stimulus conditions (numbered 1-8). Color scale indicates dissimilarity values (arbitrary units). Bottom row shows two-dimensional multidimensional scaling (MDS) visualization of the same representational geometries. Blue and red traces distinguish the two task contexts. **(B)** Three-dimensional MDS visualizations of neural representational geometries at 2040ms separated by curriculum condition (columns) and subsequent assembly performance. Top row shows participants who subsequently assembled knowledge (assemblers); bottom row shows those who did not (nonassemblers). Numbered vertices (1-8) represent stimulus conditions; blue and red traces distinguish task contexts. Tick spacing: 0.2 arbitrary units.

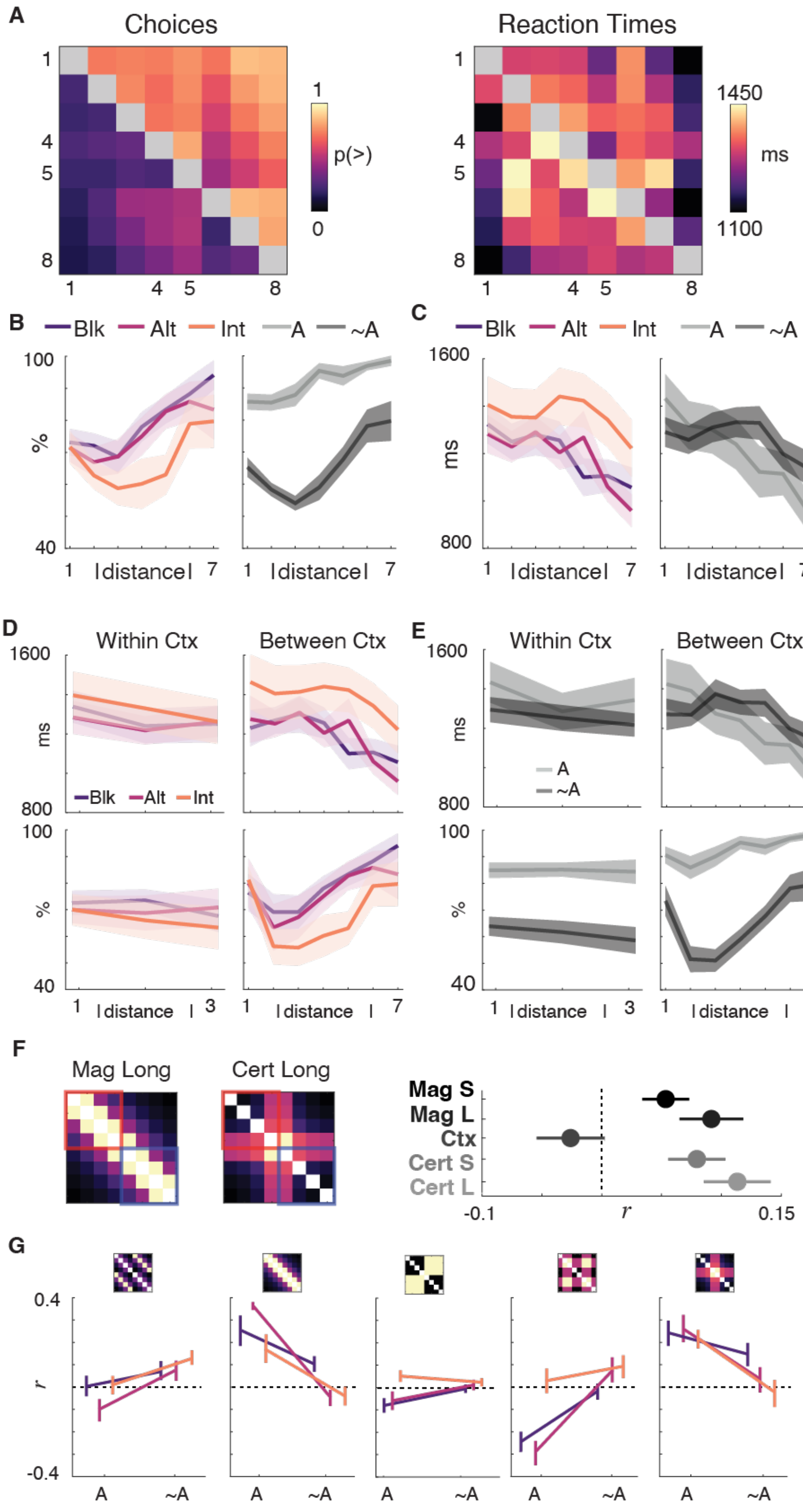

A
Choices
p(>)
Reaction Times
ms
1450
1100
B
Blk
Alt
Int
A
~A
%
Idistancel
C
ms
1600
800
D
Within Ctx
Between Ctx
E
F
Mag Long
Cert Long
Mag S
Mag L
Ctx
Cert S
Cert L
-0.1
r
0.15
G
0.4
-0.4

**Supplementary Figure 5: Behavioral performance reveals curriculum-dependent learning patterns and knowledge assembly. (A)** Heatmaps display average choice probabilities (left) and reaction times (right) across all participants during test trials. Color intensity indicates probability of selecting each option (choice matrix) and response speed in milliseconds (reaction time matrix). **(B)** Accuracy as a function of symbolic distance between choice options, separated by training condition: blocked (purple), alternating (pink), and interleaved (orange). Data are additionally split between assemblers (light gray) and non-assemblers (dark gray), showing distinct symbolic distance effects. **(C)** Reaction times as a function of symbolic distance, organized by the same training conditions and assembly groups as panel B. Both panels B and C reveal characteristic distance effects where accuracies increase and response times decrease with greater separation between comparanda. **(D)** Performance separated by comparison type for the three training conditions. Left panels show within-context comparisons; right panels show between-context comparisons. Top row displays reaction times; bottom row shows accuracy. **(E)** Same comparison structure as panel D, but contrasting assemblers (A, dark gray) and non-assemblers (~A, light gray). Shaded regions in panels B through E represent standard error of the mean (SEM). **(F)** Left: Idealized representational matrices for magnitude structure (Mag Long) and certainty structure (Cert Long). Right: Pearson correlation coefficients between neural or behavioral data and idealized matrices across different analysis scales (magnitude short/long, context, certainty short/long). Error bars indicate SEM, dotted line marks zero correlation. **(G)** Detailed correlation patterns between observed data and idealized behavioral matrices across five different structural features, separated by assembler status (A vs. ~A). Small inset matrices above each panel illustrate the specific comparison structure. Dotted lines indicate zero correlation baseline.

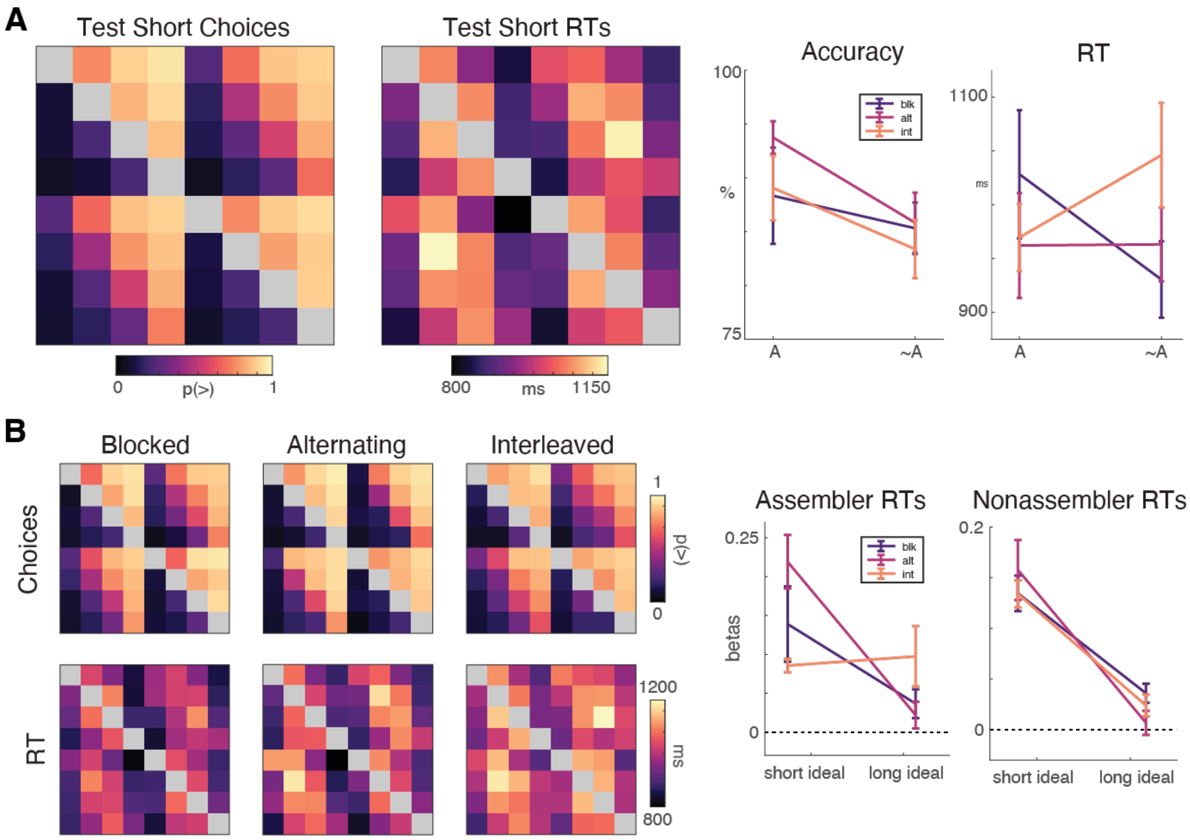


**Supplementary Figure 6: Online replication of behavioral performance during test short phase across training conditions.** Behavioral results from the online version of the experiment with expanded sample size. **(A)** Test short phase performance. Left: Average choice probability matrix (p(>)) and reaction time (RT) matrix across all participants. Right: Mean accuracy and RT as a function of assembly (A=assembler, ~A= non-assembler). Error bars represent SEM. **(B)** Test short phase performance separated by training condition. Left: Choice and RT matrices for blocked, alternating, and interleaved training schedules. Right: Beta weights from competitive regression analysis comparing observed participant matrices (blocked in purple, alternating in pink, interleaved in orange) to ideal short magnitude and long magnitude behavioral matrices. Results are shown separately for participants who did (left) and did not (right) demonstrate knowledge assembly. Dotted line at zero indicates no preference for either strategy.

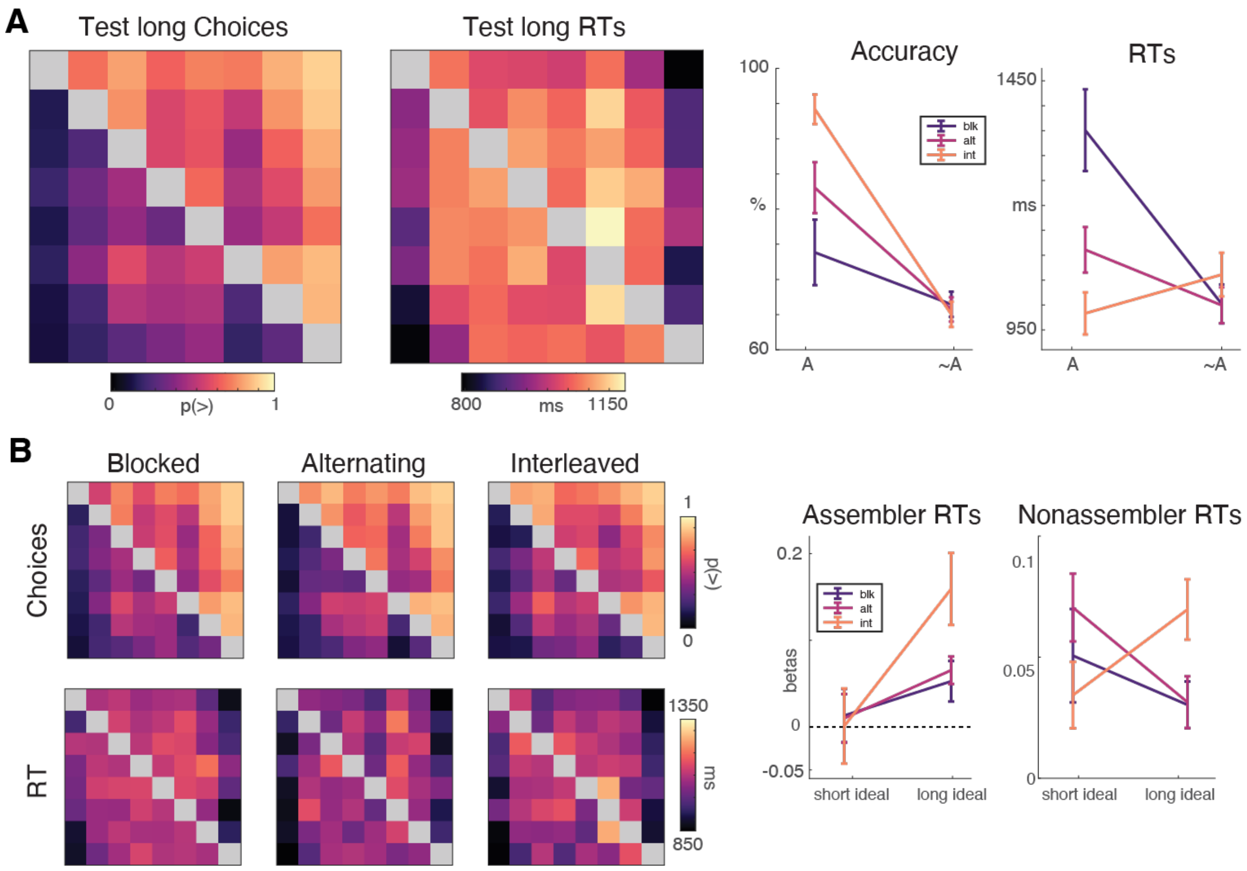


**Supplementary Figure 7: Online replication of behavioral performance during test long phase across training conditions.** Behavioral results from the online version of the experiment with expanded sample size. **(A)** Test long phase performance. Left: Average choice probability matrix (p(>)) and reaction time (RT) matrix across all participants. Right: Mean accuracy and RT as a function of assembly (A=assembler, ~A= non-assembler). Error bars represent SEM. **(B)** Test long phase performance separated by training condition. Left: Choice and RT matrices for blocked, alternating, and interleaved training schedules. Right: Beta weights from competitive regression analysis comparing observed participant matrices (blocked in purple, alternating in pink, interleaved in orange) to ideal short magnitude and long magnitude behavioral matrices. Results are shown separately for participants who did (left) and did not (right) demonstrate knowledge assembly. Dotted line at zero indicates no preference for either strategy.

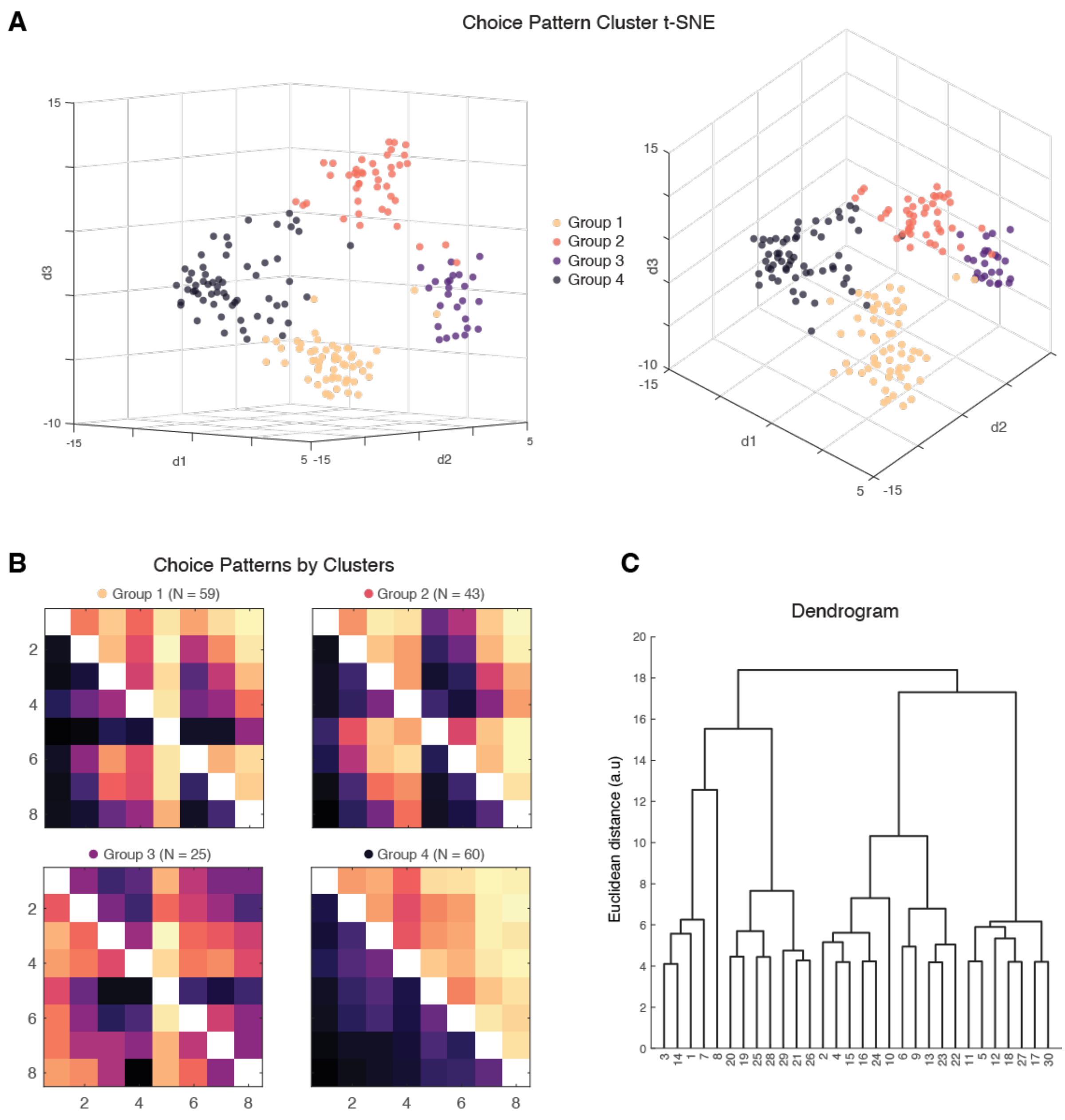


**Supplementary Figure 8: Participants exhibit distinct choice pattern clusters across experimental conditions. (A)** Two- and three-dimensional t-SNE projections of participant choice patterns reveal four distinct clusters identified via k-means clustering (Group 1, orange, N=59; Group 2, red, N=43; Group 3, purple, N=25; Group 4, gray, N=60). Data pooled from online and in-lab experiments. **(B)** Choice pattern heatmaps for each cluster show systematic differences in selection strategies across trial types. Color intensity represents choice frequency, with darker colors indicating more frequent selection of particular options. **(C)** Hierarchical clustering dendrogram supports the four-cluster solution, showing Euclidean distances between participants based on their choice patterns. The dendrogram structure validates the k-means groupings by revealing natural divisions in the data at similar linkage heights.

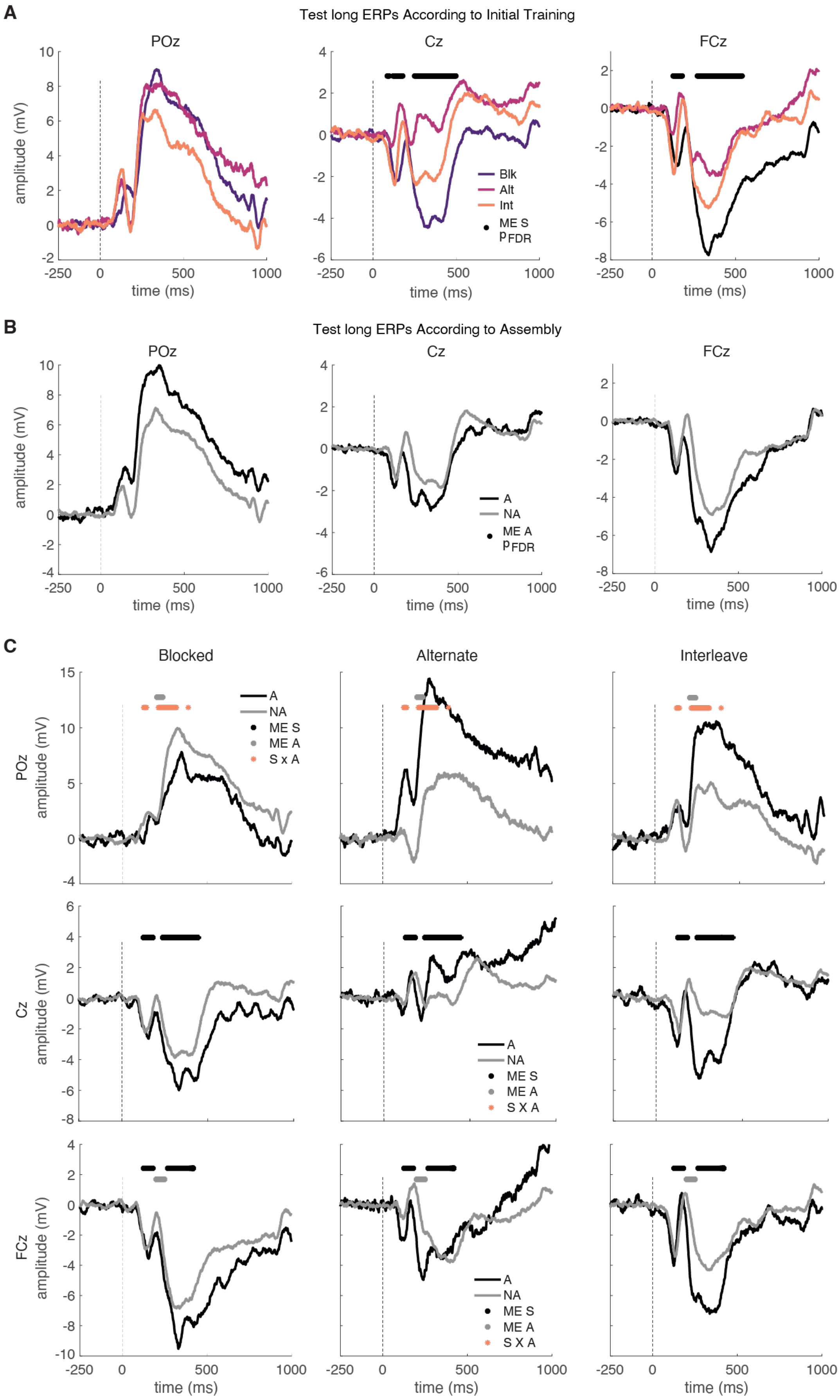
A
Test long ERPs According to Initial Training
POz
Cz
FCz
amplitude (mV)
time (ms)
Blk
Alt
Int
ME S
$p_{FDR}$
B
Test long ERPs According to Assembly
A
NA
ME A
C
Blocked
Alternate
Interleave
POz amplitude (mV)
Cz amplitude (mV)
FCz amplitude (mV)
S x A
S X A

**Supplementary Figure 9: Event-related potentials during the test long phase reveal schedule-dependent neural dynamics.** Event-related potentials (ERPs) are shown for three electrode sites (POz, Cz, FCz) during the test long phase. Time 0 indicates stimulus onset. **(A)** ERPs separated by training schedule: blocked (Blk, purple), alternating (Alt, pink), and interleaved (Int, orange). Black horizontal bars indicate time windows showing significant main effects of schedule (one-way ANOVA, FDR-corrected $p < 0.05$). **(B)** ERPs separated by assembly status: assemblers (A, dark gray/black) and non-assemblers (NA, light gray). No significant main effects of assembly were observed (one-way ANOVA, FDR correction). **(C)** Detailed breakdown showing ERPs for each schedule (columns) at each electrode (rows), with assemblers (dark) and non-assemblers (light) overlaid. Statistical markers indicate significant effects from two-way ANOVAs with FDR correction: main effect of schedule (dark dots), main effect of assembly (light dots), and schedule × assembly interaction (orange dots). ME S, main effect of schedule; ME A, main effect of assembly; S × A, schedule × assembly interaction; PFDR, FDR-corrected p-value.

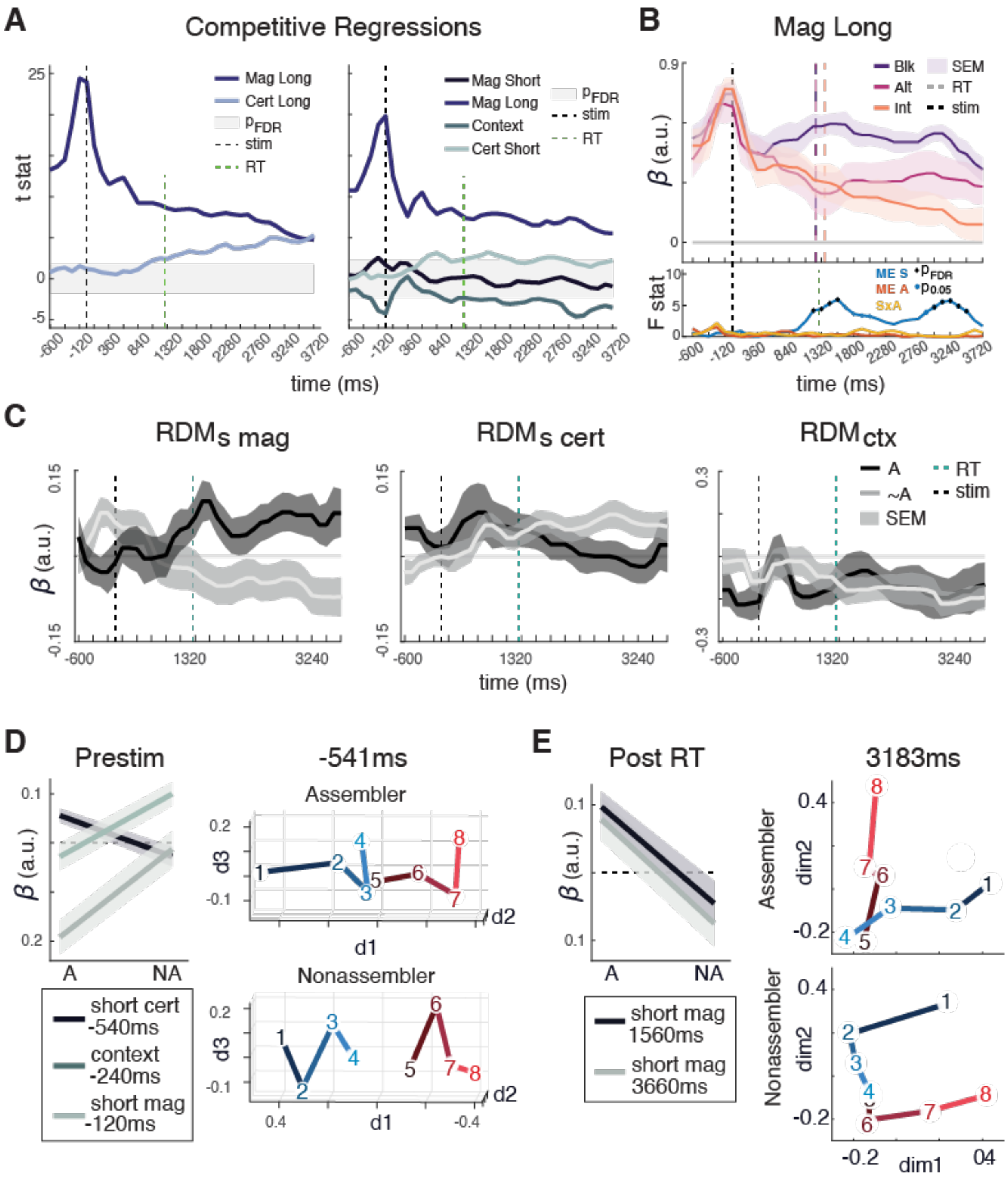


**Supplementary Figure 10: Temporal dynamics of representational geometry during knowledge assembly. (A)** Competitive regression analyses reveal temporal profiles of representational similarity. Left: t-statistics for long magnitude (purple) and certainty (light blue) RDMs regressed against participant data. Right: t-statistics for extended model including long magnitude, short magnitude, certainty, short context, and context. Shaded regions indicate non-significance after FDR correction. Vertical dashed lines denote stimulus onset (black), RT (green), P_FDR threshold (light blue), and stimulus offset (dark blue). **(B)** Long magnitude competitive regression coefficients across training schedules: blocked (purple), alternating (pink), and interleaved (orange). Shaded regions represent SEM. Lower panel shows F-statistics for main effect of schedule (ME S), main effect of assembly (ME A), and schedule-by-assembly interaction (S×A). Black dots indicate FDR-corrected significance; gray dots indicate $p < 0.05$. Vertical dashed lines as in (A). **(C)** Time-resolved representational dynamics for short magnitude ($RDM_{s\ mag}$), short certainty ($RDM_{s\ cert}$), and context ($RDM_{ctx}$). Beta coefficients shown separately for assemblers (black, solid) and non-assemblers (gray, dashed), with SEM shading. Vertical dashed lines indicate stimulus onset, RT, and stimulus offset. **(D)** Prestimulus representational geometry at three timepoints. Left: Beta coefficients for assemblers (A) versus non-assemblers (NA) across conditions. Right: Multidimensional scaling plots at -541ms showing distinct geometries for assemblers (top) and non-assemblers (bottom) across stimulus types (numbered 1-8, colored by dimension). **(E)** Post-response representational geometry. Left: Beta coefficients for short magnitude at 1560ms and 3660ms post-RT. Right: MDS plots at 3183ms showing dimensional structure for assemblers (top) and non-assemblers (bottom). Colors and numbers correspond to stimulus conditions as in (D).

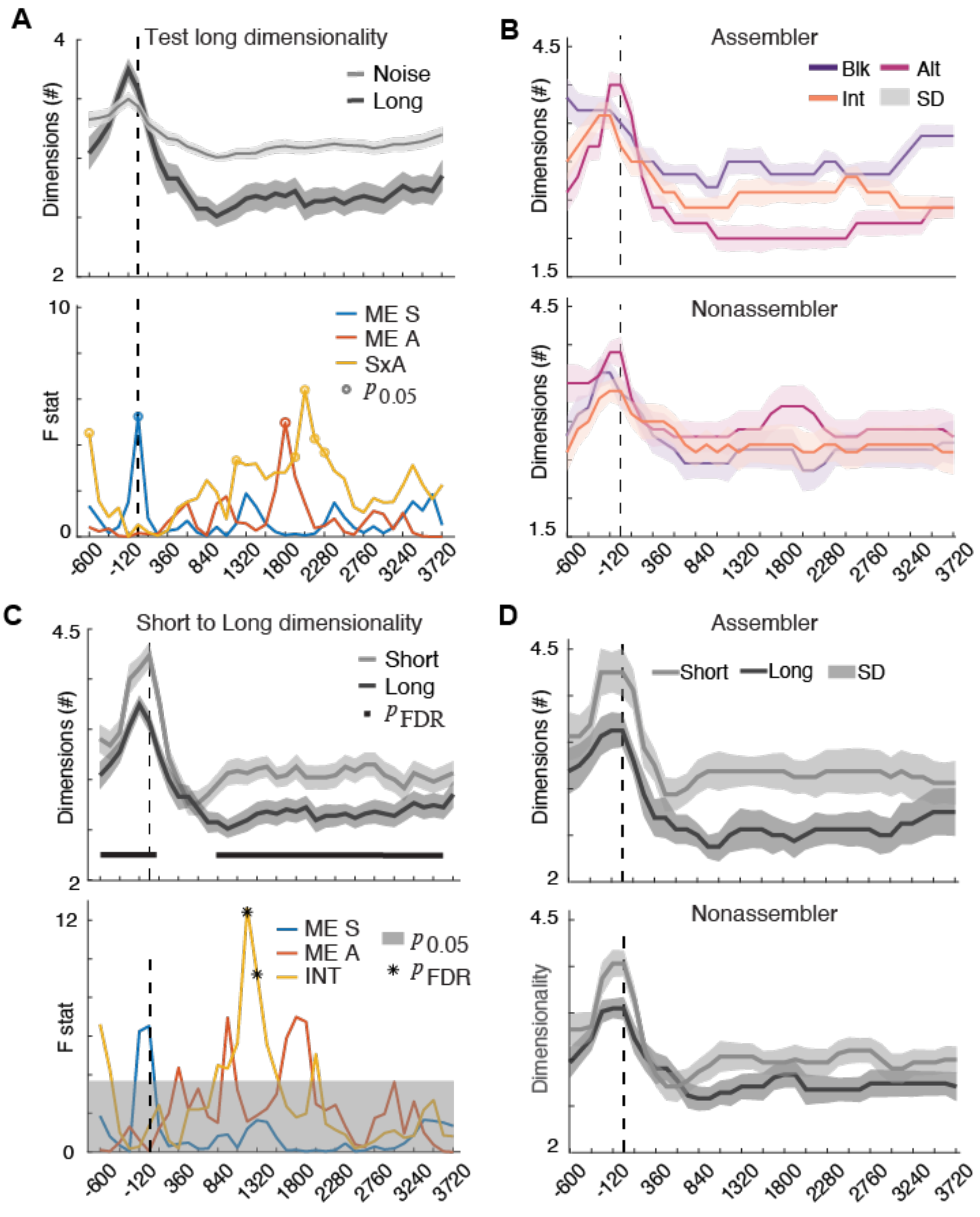


**Supplementary Figure 11: Neural representational dimensionality during test phases reveals schedule-dependent effects.(A)** Top: Neural dimensionality during the long test phase (dark line) compared to noise dimensionality (gray line), with shaded regions indicating standard deviation. Dashed vertical line marks trial 0. Bottom: F-statistics from two-way ANOVA testing main effects of initial training schedule (ME S, blue), assembly status (ME A, orange), and their interaction (SxA, red) across trials. Gray circles indicate uncorrected $p < 0.05$; no comparisons survived FDR correction. **(B)** Dimensionality trajectories during the long test phase separated by initial training schedule (blocked, purple; alternating, pink; interleaved, orange) for assemblers (top) and nonassemblers (bottom). Shaded regions show standard deviation. **(C)** Top:

Comparison of dimensionality between short test phase (gray) and long test phase (dark) with pFDR values from paired t-tests shown above. Black horizontal bars indicate time windows with significant differences. Bottom: F-statistics from two-way ANOVA examining main effects of schedule (ME S, blue) and assembly status (ME A, orange), plus their interaction (INT, red). Gray circles mark $p < 0.05$; asterisks indicate survival of FDR correction at alpha level 0.05. **d**, Dimensionality comparison between short and long test phases shown separately for assemblers (top) and nonassemblers (bottom). Line colors as in panel c.

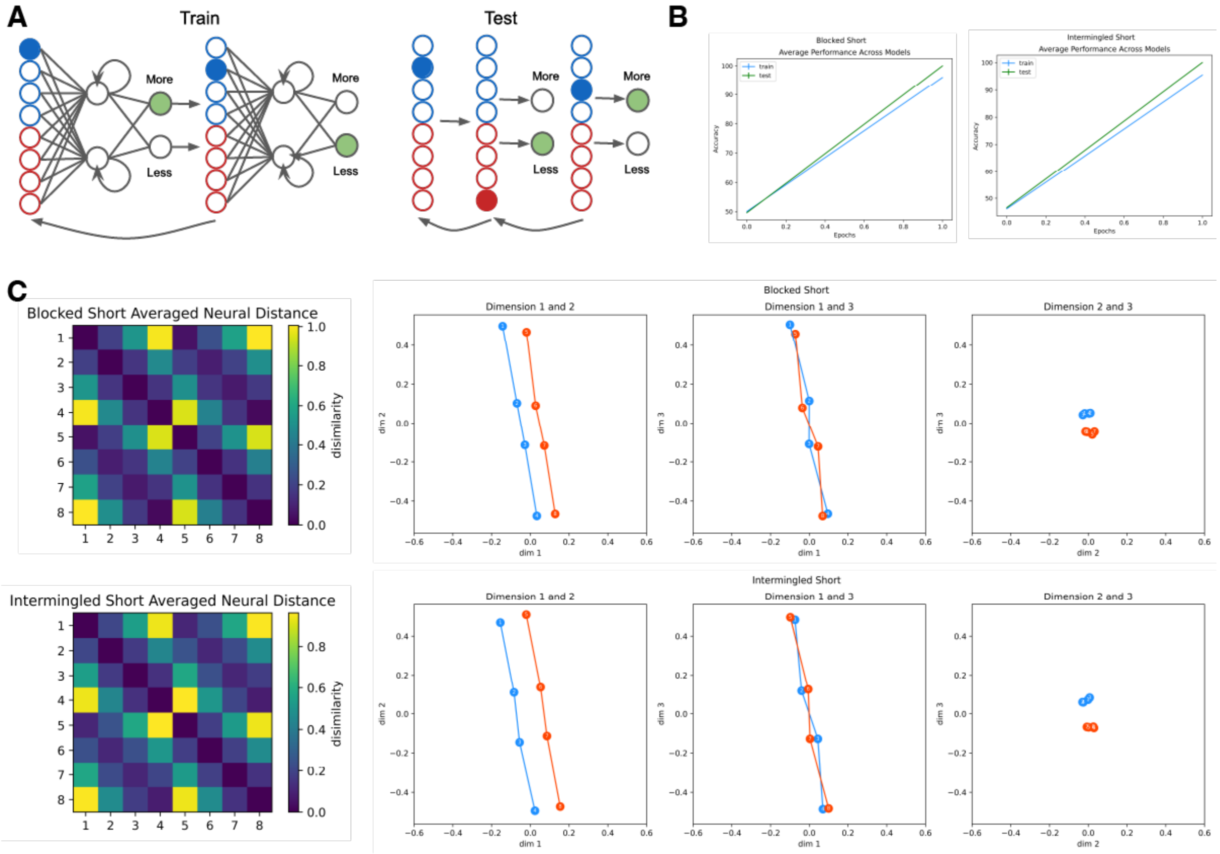


**Supplementary Figure 12: Neural representations in recurrent neural networks during blocked and interleaved training schedules. (A)** Schematic of example training (left) and testing (right) trials. Filled blue and red circles indicate the item pair being trained on each trial. Gray arrows show one-back comparisons between successive trials. Filled green circles indicate the correct response for the example comparisons shown. **(B)** Training accuracy (blue) and testing accuracy (green) over training epochs for blocked (left) and interleaved (right) curricula. Networks were trained until reaching 90% accuracy on the training set. **(C)** Representational dissimilarity matrices (RDMs; left panels) and multidimensional scaling (MDS; right panels) visualizations after blocked (top row) and interleaved (bottom row) training. Data show network representations after reaching 90% training accuracy. Blue and red lines in MDS plots represent the two contexts objects belonged to.

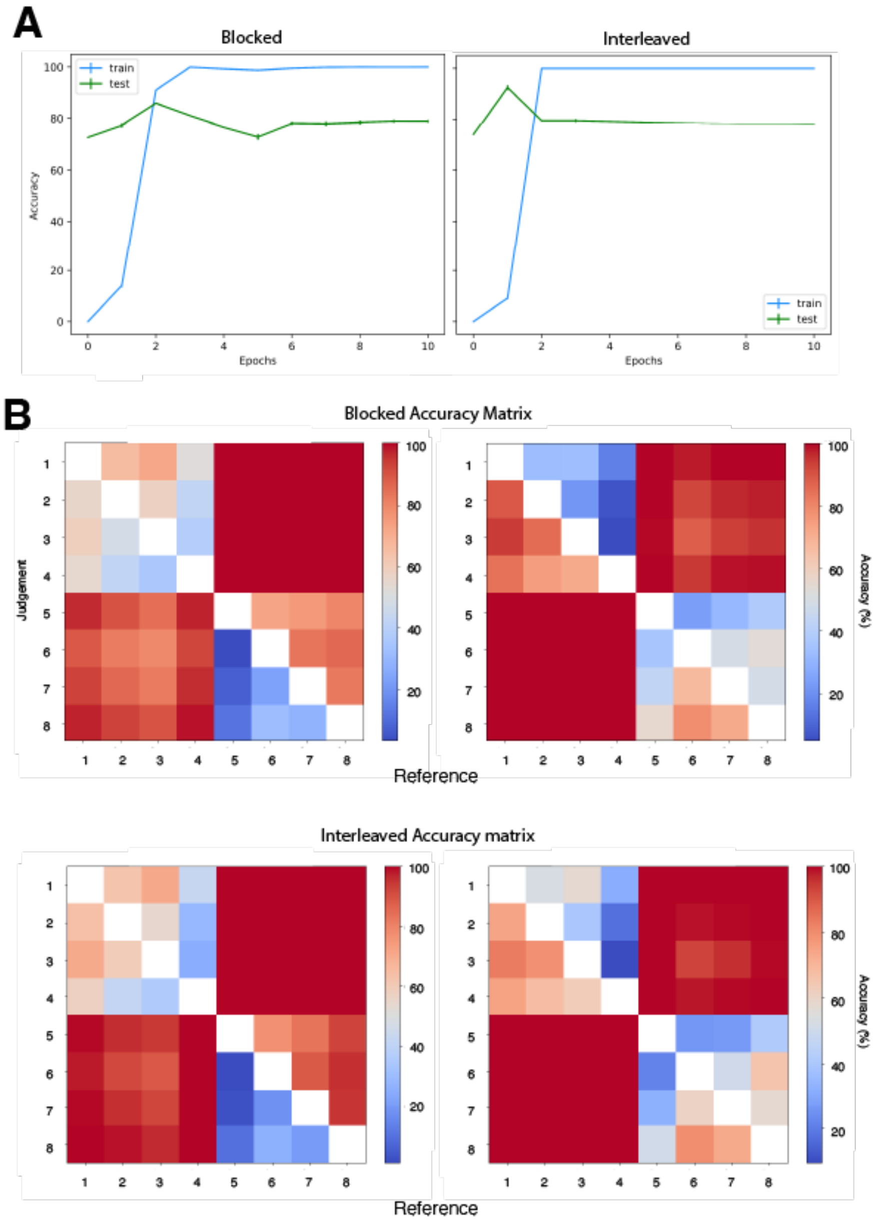


**Supplementary Figure 13: Recurrent Neural Networks Partially Assemble Knowledge. (B)** Average training and testing accuracy over training epochs for blocked (left, N=10) and interleaved (right, N=10) networks. Blue lines show training accuracy, green lines show testing accuracy. **(B)** Accuracy matrices for two example networks after blocked (top panels) and interleaved (bottom panels) training. Rows indicate judgement items, columns indicate reference items. Red indicates high accuracy, blue indicates low accuracy.

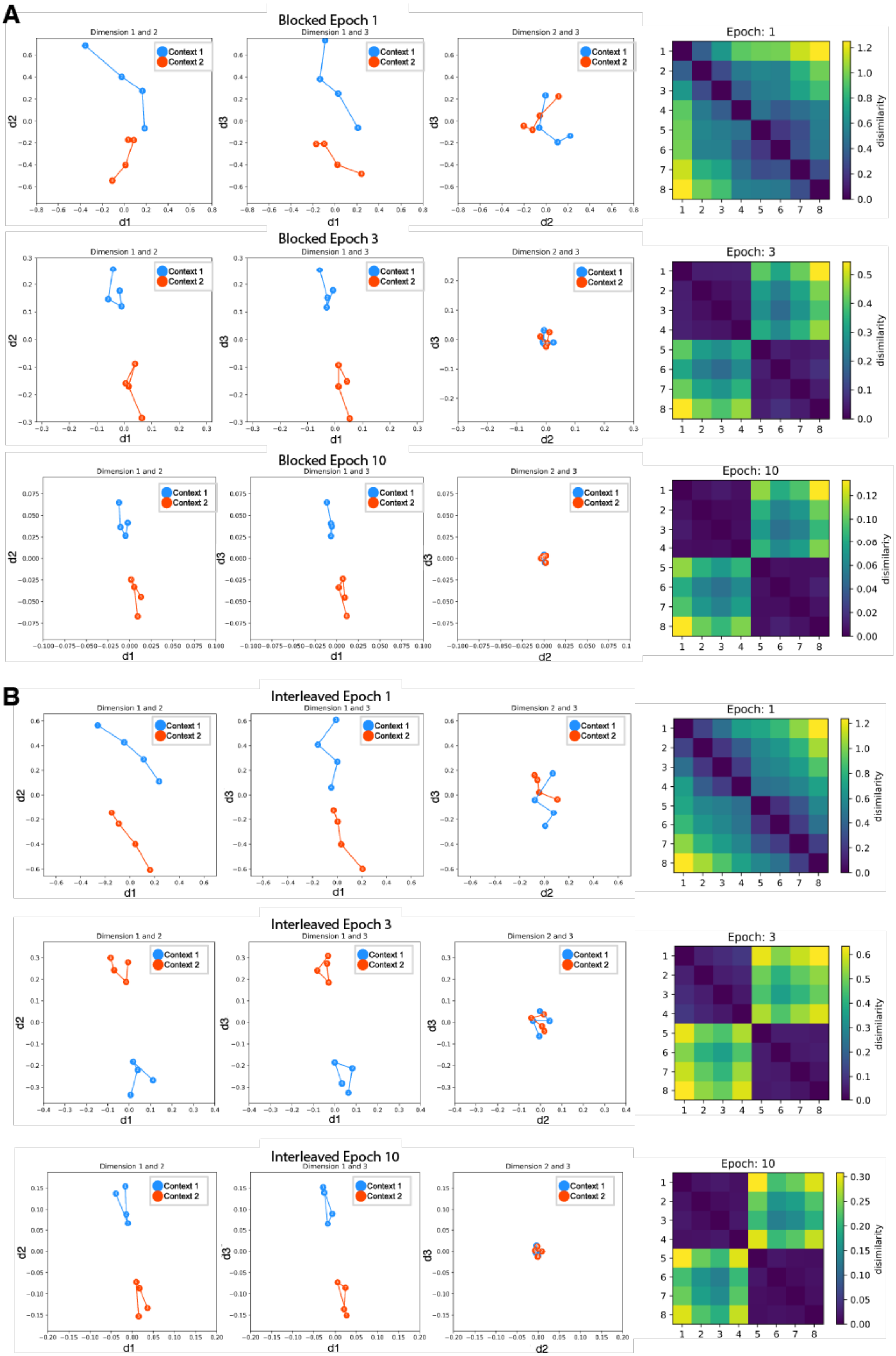
A
Blocked Epoch 1
Dimension 1 and 2
Dimension 1 and 3
Dimension 2 and 3
Context 1
Context 2
d1
d2
d3
Epoch: 1
disimilarity
Blocked Epoch 3
Epoch: 3
Blocked Epoch 10
Epoch: 10
B
Interleaved Epoch 1
Interleaved Epoch 3
Interleaved Epoch 10

**Supplementary Figure 14: Temporal evolution of neural representations under blocked and interleaved training. (A)** Multidimensional scaling (MDS) plots (left three columns) and representational dissimilarity matrices (RDMs; right column) for networks trained with blocked curricula. Neural distance is shown as dissimilarity from close (blue) to far (yellow). Neural representations are shown at epoch 1 (top row), epoch 3 (middle row), and epoch 10 (bottom row) out of 10 total training epochs. Blue circles represent items initially trained in context A, red circles represent items from context B. **(B)** MDS plots and RDMs for networks trained with interleaved curricula, shown at the same training epochs as in (A). Color coding follows the same convention as panel (A).

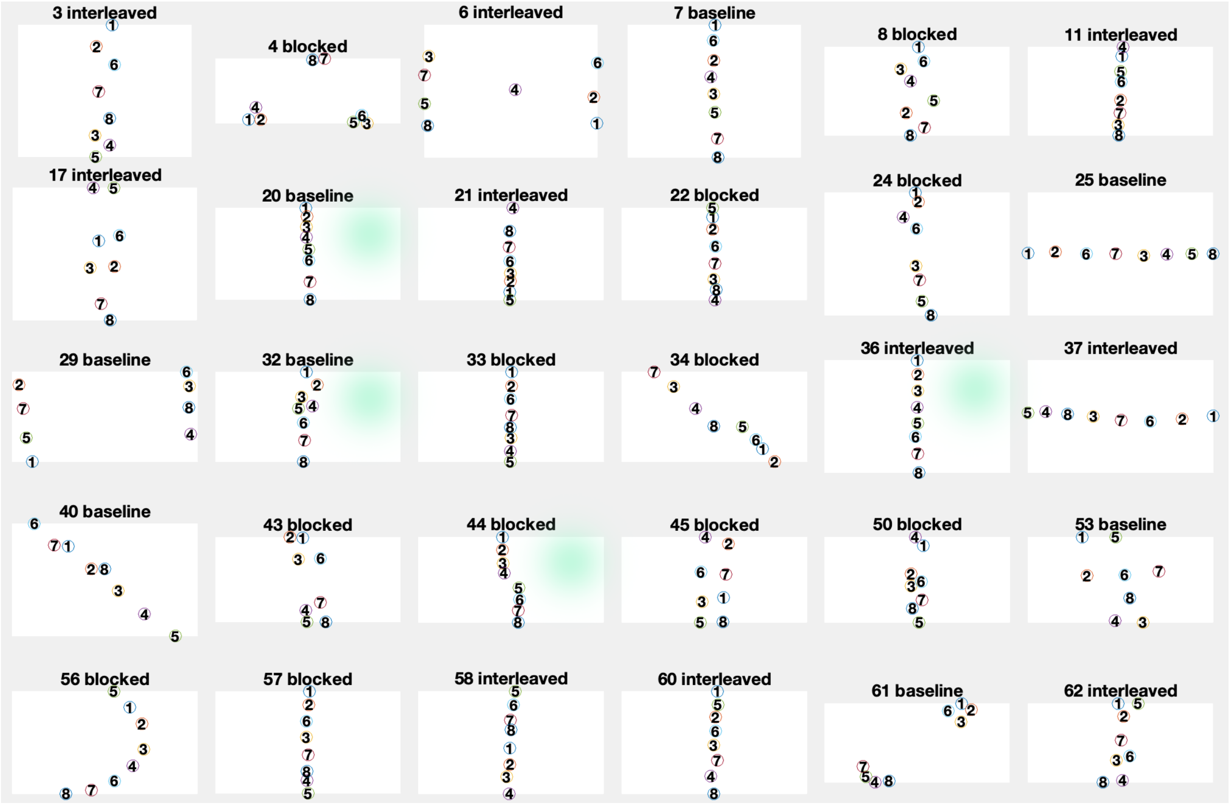


**Supplementary Figure 15: Example free arrangement task responses across participants and training conditions.** Representative spatial arrangements from the post-experimental free arrangement task, where participants organized eight object cards (numbered 1-8) by "brispness" in an unconstrained 2D workspace. Each panel shows a single participant's final arrangement, labeled with participant number and initial training condition (blocked, baseline, or interleaved). Green highlighting indicates baseline condition participants who did not receive any experimental manipulation before the arrangement task. Spatial coordinates from these arrangements were extracted and correlated with the ground truth relational structure ($i_1$-$i_8$) to classify participants as assemblers ($r \geq 0.9$) or non-assemblers ($r < 0.9$). The diversity of spatial configurations illustrates individual differences in how participants represented the learned relational structure, with some showing clear linear or grouped arrangements while others demonstrated more scattered or context-separated patterns.

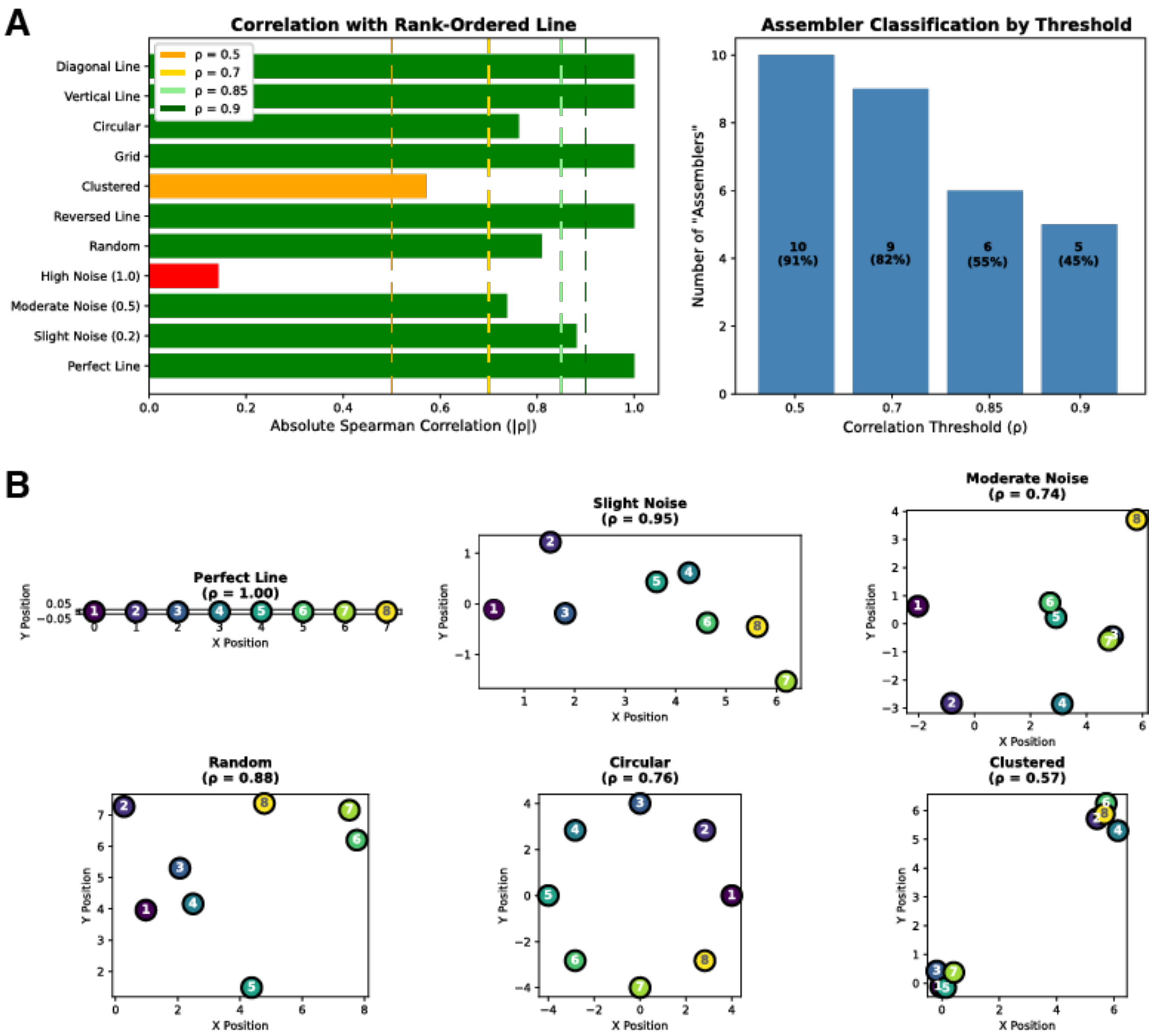


**Supplementary Figure 16: Simulations for determining correlation threshold for assembly.(A)** Left: Absolute Spearman correlations (|ρ|) between simulated 8-object arrangements and a canonical rank-ordered line. Arrangements span perfect linear structures (ρ = 1.0), noisy configurations with varying Gaussian noise (σ = 0.2-1.0), geometric patterns (grids, circles), clustered layouts, and random placements. Colored bars indicate classification under four threshold criteria: liberal (ρ ≥ 0.5, orange), moderate (ρ ≥ 0.7, yellow), stringent (ρ ≥ 0.85, light green), and very stringent (ρ ≥ 0.9, dark green). Right: Number of arrangements classified as "assembled" under each threshold. The ρ ≥ 0.9 criterion identifies approximately half of arrangements (45%), capturing only those with clear spatial organization. **(B)** Representative spatial arrangements illustrating the relationship between visual structure and correlation strength. Numbers indicate object ranks (1-8). Perfect lines achieve ρ = 1.0 regardless of orientation (horizontal, vertical, diagonal) due to rotation-invariant correlation calculation. Arrangements with ρ = 0.88 (Random) exhibit substantial positional variability despite statistically significant correlation, supporting use of the stringent ρ ≥ 0.9 threshold. Moderate noise (ρ = 0.74) and clustered arrangements (ρ = 0.57) show progressively weaker spatial organization.

#### Supplementary Results

*Train Short*

Forty-eight participants were recruited, but one interleaved participant did not reach criterion and was excluded (final N = 47; Interleaved N = 15). Criterion was reached after 4.4 ± 1.2 blocks for blocked participants, 4.8 ± 1.6 for alternating, and 4.8 ± 1.5 for interleaved (Mean ± SD). As reported in the main text, training trials did not differ according to context identity (Fig S1A; ME ID: $F_{1,82} = 0.05$, p = 0.8), indicating that the arbitrary assignment of objects to Context A versus Context B did not systematically affect learning difficulty. However, we observed an important effect of training schedule on transfer learning (Fig. 1D). Blocked and alternating training specifically facilitated transfer to the second-encountered context, reducing the number of training trials required (Schedule × Order: $F_{2,82} = 6.8$, p = 0.002; ME Order: $F_{1,82} = 3.2$, p = 0.08; 3-way ANOVA).

Leave-one-channel-out analysis suggested distributed cortical encoding with marginal contributions from lateralized midline channels (Fig. S1B; C3, P7, C4, Cp6: $2.0 < t_{46} < 3.5$, $p < 0.05$, uncorrected, p's > $p_{FDR}$= 0). This suggests context information was encoded through population-level activity patterns rather than localized sources. The temporal dynamics of context coding revealed stable representations throughout the trial period (Fig. S1C, left panel; ME Time: $F_{7,39} = 0.6$, p = 0.7), allowing us to collapse across these windows for subsequent analyses as reported in the main text.  When we examined context coding as a function of future assembly success, we again found that context representations were present in all schedules except interleaved training (Fig. S1C, right panel). This pattern held regardless of whether participants would later successfully assemble knowledge, indicating that context codes during initial learning do not directly predict assembly capacity. Along these lines, context coding strength during initial training correlated negatively with reaction times during *test short* (Fig. S1D, left panel; $r_{46} = -0.36$, p = 0.02), but not during *test long* ($r_{46} = -0.18$, p = 0.2). This suggests that context representations formed during initial learning primarily support efficient retrieval within the learned structure rather than enabling flexible reorganization across contexts. Correlations with accuracy during *test short* ($r_{46} = 0.19$, p = 0.2) and *test long* ($r_{46} = 0.03$, p = 0.8) were not significant (Fig. S1D, right panel).

*Test Short Behavioral*

Training schedule influenced the efficiency but not the accuracy of transitive inference during test short. While overall accuracy did not differ across blocked, alternating, and interleaved conditions, we observed that interleaved initial training led to slower reaction times regardless of assembly status (Fig. S2A). This is evident in both average reaction times as well as reaction times separated out by symbolic distance (Fig. S2B)

*Test Short Neural*

Neural representations during *test short* revealed temporal dynamics that extended beyond the main response window analyzed in the manuscript. Pre-stimulus certainty estimates were negative before stimulus onset (Fig. S3B; -600 to -240ms, $t_{46} \leq -2.4$, p's $\leq 0.02 = p_{FDR}$), suggesting preparatory states that differed systematically from the certainty structure that would emerge later in the trial. While certainty representations appeared marginally late in the trial period for assemblers ($F_{1,41} = 3.5–5.9$, $p_{FDR} <$ p's $< 0.04$, 2640–3240ms), this pattern was absent during the post-decision period for interleaved assemblers (Fig. S3D). During a post-decision window from 1920–2160ms representations were effectively low-dimensional across most participants when compared to within-participant baselines (Fig. 3E; paired t-tests against within participant baselines: $t_{46} = -4.8$, $p < 0.001$). However, interleaved participants who would later successfully assemble showed no such dimensional collapse ($t_5 = -0.5$, $p = 0.6$), maintaining higher-dimensional representations throughout the post-decision period (Fig. S3E).

*Train Long Behavioral*

The boundary training phase served as a critical minimal learning episode that could trigger knowledge reorganization. Performance on the single boundary item relation (comparing items 4 and 5) was uniformly high across training schedules (ME Schedule: $F_{2,41} = 0.4$, $p = 0.7$) and did not differ as a function of whether participants would later demonstrate successful assembly (ME Assembly: $F_{1,41} = 1.9$, $p = 0.2$). This null result is important because it indicates that assembly failure was not due to inadequate encoding of the linking information itself, but rather reflected differences in how this boundary information was integrated with existing knowledge structures.

*Test Long Behavioral*

The introduction of boundary information produced substantial changes in performance. Compared with *test short*, accuracies (68.4 ± 21.1%, mean ± SD) declined ($t_{46} = 6.4$, $p < 0.001$) and response times (1285.9 ± 346.8ms, mean ± SD) lengthened ($t_{46} = –4.6$, $p < 0.001$; paired t-tests against *test short*). This was largely due to a group of participants who did not assemble knowledge - performance averaged 85.6 ± 12.4% SD for assemblers and 61.5 ± 12.2% SD for nonassemblers. Reaction times showed no reliable modulation by schedule or assembly success (Table S3).

The pattern of symbolic distance effects (SDEs) during test long provided evidence for knowledge restructuring (Fig. S5B-E). Symbolic distance effects reflected the integrated 8-item ordering (Fig. S5) (RT: $t_{46}$ = -5.5, p < 0.001; Accuracy: $t_{46}$ = 6.0, p < 0.001; t-tests against zero), particularly in assemblers (ME Assembly: ACC – $F_{1,41}$ = 61.2, p < 0.001; RTs – $F_{1,41}$ = 19.0, p < 0.001; 2-way ANOVA on regression estimates). SDEs were clearly apparent for between-context comparisons in both accuracy and reaction times (Accuracy: $t_{46}$ = 2.6, p = 0.01; RT: $t_{46}$ = -4.7, p < 0.001), but critically, they were absent for within-context comparisons (Fig. S5D,E; all p's > 0.1). This asymmetry suggests that participants experienced uncertainty specifically about relations within the original contexts, not between them. When we examined which distance model better characterized behavior using a competitive regression (short-distance reflecting the original 1-4 structure in each context versus long-distance reflecting the integrated 1-8 structure; see Methods), we found that both models contributed to performance across participants (Short: $t_{46}$ = –2.6, p = 0.01; Long: $t_{46}$ = –4.8, p < 0.001). Importantly distance effects that reflected the integrated 1-8 structure were associated with both blocked initial training and successful assembly (Fig. S5 B,C,E; ME Schedule: $F_{2,41}$ = 3.8, p = 0.03; ME Assembly: $F_{1,41}$ = 11.6, p = 0.002).

Model based correlation analyses confirmed that assemblers' reaction times aligned strongly with the integrated long-distance magnitude model (ME Assembly - Mag long; $F_{1,41}$ = 44.7, p < 0.001; 2-way ANOVA on correlations), while non-assemblers continued to behave according to the short-distance model reflecting the original separate contexts (ME Assembly - Mag Short; $F_{1,41}$ = 10.5, p < 0.01; 2-way ANOVA on correlations). Blocking enhanced resemblance with the elongated magnitude structure (Mag Long - ME Schedule: $F_{2,41}$ = 3.20, p = 0.05). Blocking enhanced Long Magnitude resemblance particularly for non-assemblers (Schedule×Assembly: $F_{2,41}$ = 4.2, p = 0.02). Blocked non-assembler responses resembled Long Magnitude more than alternating or interleaved non-assemblers ($t_{30}$ = 3.3, p < 0.01; unpaired t-tests). Conversely, interleaved participants resembled the elongated magnitude structure less than blocked ($t_{31}$ = 2.4, p = 0.02; unpaired t-tests) with interleaved assemblers indistinguishable from blocked non-assemblers (p > 0.3). Behavioral alignment with updated rank-certainty tracked assembly (Cert long - ME Assembly: $F_{1,41}$ = 13.5, p < 0.001). Negative correlations with context were linked to assembly (ME Assembly: $F_{1,41}$ = 24.2, p < 0.001) and blocked training (ME Schedule - $F_{2,41}$ = 7.3, p = 0.002), indicating that successful reorganization required breaking down rather than maintaining context-dependent codes.

To adjudicate between competing models of behavioral reorganization, we performed a competitive regression distinct from the result the main text (Table S4, Table S5). This regression included context and did not include the idealized certainty matrix from *test short* (Table S7). Context was negatively correlated with Long Magnitude (Table S1) and did not explain responses when included in a competitive regression alongside Magnitude and Certainty idealized matrices ($t_{46}$ = –0.3, p = 0.8). Furthermore, context estimates obtained from a competitive regression were no longer linked to assembly, while Long Magnitude and Long Certainty effects were replicated (Table S7). Together, these results show that boundary training effectively reorganized behavior, with assemblers flexibly integrating across contexts, particularly with blocked initial training.

*Online Study Test Short Behavioral*

To establish the generalizability of our findings beyond the laboratory setting with EEG recording, we conducted an online replication study with a substantially larger sample (N = 140). During test short, average accuracy and reaction times closely matched the laboratory study (Fig. S6; 86.0 ± 15.1% and 981.6 ± 239.6ms respectively, mean ± SD). We found no significant effects of training condition or future assembly success on accuracy or reaction times during this phase (ACC: ME Schedule: $F_{2,134} = 0.5$, p = 0.6; ME assembly: $F_{1,134} = 2.4$, p = 0.1; Schedule×Assembly: $F_{2,134} = 0.2$, p = 0.8. RTs: ME Schedule: $F_{2,134} = 0.2$, p = 0.8; ME assembly: $F_{1,134} = 0.01$, p = 0.9; Schedule×Assembly: $F_{2,134} = 0.8$, p = 0.5), consistent with the in-lab results showing that training schedules primarily affected efficiency rather than basic inference ability.

When we examined the structure of reaction time matrices by regressing them against idealized models for both test phases (see Methods). Both *test short* and *test long* idealized magnitude matrices were evident in the responses over all participants (Fig. S6; $t_{139} = 14.1$, p <0.001; $t_{139} = 4.8$, p < 0.001 respectively). Training condition showed marginal effects on alignment with the test short magnitude structure (ME Schedule: $F_{2,134} = 2.7$, p = 0.07), but no clear impact of assembly status at this stage (ME assembly: $F_{1,134} = 0.04$, p=0.84; Schedule×Assembly: $F_{2,134} = 1.17$, p = 0.31). For alignment with the test long structure, we observed marginal effects of both condition and assembly (ME Condition: $F_{2.134} = 2.9$, p = 0.06; ME assembly: $F_{1,134} = 3.7$, p = 0.05; Schedule×Assembly: $F_{2,134} = 1.6$, p = 0.19).

*Online Study Test Long Behavioral*

The online test long phase largely replicated the laboratory findings while revealing some interesting differences related to testing context. Overall accuracy and reaction times again matched the laboratory study (Fig. S7; 68.8 ± 13.1% and 1052.8 ± 253.9ms respectively; mean ± SD), with participants showing reduced accuracy and increased response times compared to *test short* ($t_{139} = -12.0$, p < 0.001) and slower ($t_{139} = -4.8$, p < 0.001) from test short to test long. Training condition had significant effects on accuracy (ACC: ME Schedule: $F_{2,134} = 3.6$, p=0.03; Schedule×Assembly: $F_{2,134} = 4.80$, p = 0.01) that were driven by a monotonic pattern across schedule among assemblers. As expected, there was also a main effect of assembly success (ME assembly: $F_{1,134} = 43.93$, p < 0.001). Response times showed similar but weaker effects (ME Schedule: $F_{2,134} = 3.0$, p = 0.5; ME Assembly: $F_{1,134} = 4.9$, p = 0.03; Schedule×Assembly: $F_{2,134} = 4.4$, p = 0.01).

We then investigated the form of the RT matrices by regressing them against idealized matrices for *test short* and *test long* (Fig. S7; see Methods). Both *test short* and *test long* idealized matrices were evident in the responses over all participants (short - $t_{139} = 4.9$, p <0.001; long - $t_{139} = 8.3$, p < 0.001). The effects of training condition ($F_{2,134} = 6.4$, p = 0.002) as well as assembly status ($F_{1,134}$ = 9.0, p=0.003; Schedule×Assembly: $F_{2,134}$ =1.3, p = 0.15) on correspondence with the long magnitude matrix were robust. Critically, these effects were specific to the long magnitude structure; the test short magnitude matrix did not show reliable schedule or assembly effects during test long (ME Schedule: $F_2$ = 0.3, p = 0.8; ME Assembly: $F_{1,134}$ = 3.4, p = 0.07; Schedule×Assembly $F_{2,134}$ = 0.2, p = 0.9)

*Test Long Event Related Potentials*

Event-related potentials during test long revealed how training schedules shaped the temporal dynamics of neural processing during knowledge assembly. At central electrodes (Cz and FCz), blocked participants again showed stronger N400 components, similar to the pattern during *test short* (Fig. S9; ME Schedule - Cz: $3.6 < F_{2,41} < 20.6$, $p < p_{FDR} = 0.04$; FCz: $3.6 < F_{2,41} < 11.1$, $p < p_{FDR} = 0.03$, 238–499ms). This enhanced prediction error signal under blocked training extended into the assembly phase, suggesting that blocked learners continued to maintain strong contextual predictions even as they integrated across contexts. In contrast, interleaved and alternating participants exhibited increased P2 amplitudes during *test long* (Cz: $3.4 < F_{2,41} < 15.1$, $p < p_{FDR} = 0.04$, 126–181ms; FCz: $3.6 < F_{2,41} < 9.5$, $p < p_{FDR} = 0.03$, 126–183ms), which may reflect heightened perceptual processing demands when representations are more distributed.

At posterior electrode PoZ, ERP amplitudes clearly differentiated assembly outcomes. Assemblers showed attenuated N2 responses and faster P3 latencies, resulting in increased amplitudes (Fig. S9; $8.5 < F_{1,41} < 11.3$, $p < p_{FDR} = 0.006$, 206–239ms). This pattern suggests that successful assembly was associated with more efficient stimulus categorization and faster deployment of attentional resources. The Schedule × Assembly interaction that emerged during this period mirrored findings from test short: interleaved and alternating assemblers showed increased P3b amplitudes, while blocked assemblers did not ($4.2 < F_{2,41} < 6.4$, $p < p_{FDR} = 0.02$). This dissociation reinforces the conclusion from the main text that different representational strategies support assembly under different training regimes.

*Test Long Neural Geometries*

We first conducted “Model 1” - a competitive regression including Long Magnitude and Long Certainty (see Methods). These model RDM derived from hypothesized geometries underlying the *test long* phase were reflected in neural codes (Fig. 5A). However, neither elongated magnitude nor certainty representations distinguished assemblers from non-assemblers (Fig. 5A) ($F_{1,41} < 2.2$, p > 0.1), indicating that simply encoding the new task structure is insufficient for successful reorganization.

We then conducted “Model 2” - a competitive regression analysis that tested how neural representational dissimilarity matrices (RDMs) related to idealized magnitude models from both phases of the experiment alongside certainty and context from *test short* (Fig. S10, see Methods). Estimates reflecting $RDM_{l\ mag}$ confirmed dynamics of elongated representation during assembly reported in the main text. Estimates of the long magnitude structure ($RDM_{l}$ mag) persisted throughout the trial in participants who had received more blocked initial training (Fig. S10; ME Schedule: $4.1 < F_{1,41} < 6.0$, p's ≤ $p_{FDR} = 0.03$). This schedule effect was independent of assembly success, indicating that blocked training promoted encoding of the integrated structure regardless of whether participants explicitly recognized this organization ($F_{1,41} < 2.2$, p's > 0.1).

Neural geometries were effectively low-dimensional over most of the trial (Fig. S10A; $t_{46} < -2.5$, p's < $p_{FDR}$= 0.02; paired t-tests). Specifically, *test long* dimensionalities were consistently lower than noise dimensionalities over most of the trial (Fig. S11; -600, -480, -120ms, 240ms-3724ms), indicating that neural activity occupied a compressed representational space. Neither training schedule nor assembly status independently predicted test long dimensionality after correction for multiple comparisons (p's > $p_{FDR} = 0.002$, Schedule: $F_{2,42} < 5.3$, p > 0.009; Assembly: $F_{1,42} <$ 5.0, p's > 0.03; Schedule×Assembly: $F_{2,42} < 6.4$, p's > $p_{FDR} = 0.002$). However, critical effects emerged when examining changes in dimensionality from *test short* to *test long*.

Assembly under different training schedules was associated with distinct trajectories of dimensional change (Fig S10C). Successful assemblers who underwent blocked learning showed lower dimensionalities during *test short* and higher dimensionalities during *test long*. This crossover pattern suggests that blocked learners initially formed compressed certainty-based representations that subsequently expanded during reorganization to accommodate the integrated structure. This interaction was statistically robust at specific timepoints (1201ms and 1321ms), surviving correction for multiple comparisons (Fig. S10C, bottom panel; Schedule×Assembly: $F_{2,42} = 12.6, 9.2$, p's < $p_{FDR} = 0.002$; ME *F'*s < 7.0, p's > $p_{FDR} = 0.002$). These findings demonstrate that training history constrains not only the content of neural codes but also their complexity, and that successful assembly can be achieved through multiple routes.

*Recurrent Neural Network Test Short*

To test whether simple associative learning mechanisms could account for the observed phenomena, we trained vanilla recurrent neural networks on the magnitude ordering task under blocked and interleaved curricula. During the *test short* phase, RNNs showed robust learning performance, with training accuracy stabilizing above 90% for both blocked and interleaved conditions (Fig. S11B). However, unlike human participants, we observed no significant behavioral differences between the two training schedules during this phase (Fig. S11B). This null result suggests that the human behavioral and neural differences we observed require computational mechanisms beyond simple temporal association learning.

Neural representations in the RNN models showed some similarities to human brain activity. Specifically, the models successfully learned to represent ordinal information in a systematic way across their hidden layer representations, displaying parallel neural axes for magnitude emerged across contexts, consistent with the structured representations observed in human EEG recordings (Fig. S12C). Instead, model representations showed more linear organization of magnitude information, suggesting that additional computational principles are needed to produce certainty-weighted codes.

*Recurrent Neural Network Test Long*

During the test long phase, when the boundary item was introduced to link the two previously separate contexts, models demonstrated some capacity for learning this new relationship. Test accuracy on comparisons along the entire axis averaged approximately 70-90%, which is comparable to human performance levels (Fig. S13A). However, the pattern of performance revealed important differences from human behavior. Test long accuracies did not differ between training conditions (Overall: $t_{18} = -0.1$, $p > 0.9$; Within: $t_{18} = 0.4$, $p = 0.7$; Between: $t_{18} = -0.3$, $p = 0.8$) and symbolic distance effects did not differ between blocked and interleaved training ($t_{18} = 1.3$, $p = 0.2$). Between-context comparisons (comparing items from different contexts) reached ceiling accuracy and showed highest performance at the extreme absolute distances (distances 1 and 7). In contrast, within-context comparisons showed a marked decline toward chance levels, with many models exhibiting complete performance inversion for these trial types (Fig. S13D).

Analysis of model training dynamics revealed that accuracy peaked rapidly, typically after 1-3 epochs. Extended training beyond this point was associated with a "crumpling" of neural representations, where the structured geometry observed in earlier epochs deteriorated (Fig. S14A,B bottom rows).

*Post-Experimental Debrief*

Following completion of the experiment, participants completed a free arrangement task that provided a manipulation-check for explicit knowledge of the relational structure. The spatial arrangements participants created varied considerably (Fig. S15), ranging from clear linear orderings to grouped or scattered configurations. We extracted the spatial coordinates of the final arrangement and correlated them with the ground truth ordering. Participants whose arrangements yielded Spearman correlation coefficients of $\rho \geq 0.9$ with the ground truth structure were termed "Assemblers" (correlation range across all participants: 0.08-0.99). A threshold of $\rho \geq 0.9$ identifies participants who demonstrated strong linear ordering while allowing for minor spatial imperfections inherent to manual placement in a free arrangement task. This criterion is more stringent than the conventional threshold for a "strong" correlation ($r \geq 0.7$) and ensures that classified "Assemblers" correctly positioned most items in their proper sequence. Arrangement correlations averaged $0.75 \pm 0.31$ for blocked participants, $0.82 \pm 0.25$ for alternating participants, and $0.61 \pm 0.42$ for interleaved participants (Mean ± SD).

Qualitative responses to debriefing questions revealed that participants generally overestimated the number of objects in the experiment, reporting an average of 9.4 ± 3.0 objects (Mean ± SD). objects when there were only 8. Interestingly, assemblers were more accurate in their count estimates (mean 8.1) than non-assemblers (mean 10.0), suggesting that successful integration of the structure was associated with more accurate metacognitive representations of the task. Along these lines, both groups reported similar understanding of the test short task when asked to rate their agreement with "I understood what I was being asked to do in Test 1" on a scale from 1 (strongly disagree) to 5 (strongly agree). Assemblers and non-assemblers averaged 4.3 and 4.4 respectively on this item (4.4 ± 0.8 across all participants, Mean ± SD). However, ratings diverged markedly for test long. Assemblers reported higher understanding (4.7) compared to non-assemblers (3.4), with an overall mean of 3.8 ± 1.3 SD across all participants. This divergence suggests that non-assemblers were aware of the increased difficulty they experienced during test long, and that assembly success was accompanied by subjective feelings of comprehension. These patterns complement the behavioral and neural measures by demonstrating that knowledge assembly in our task was associated with both objective performance improvements and subjective phenomenology.

#### Supplementary Tables

| Idealized Accuracy Matrices | | | | |
|---|---|---|---|---|
| | Mag long | Context | Cert short | Cert long |
| Mag short | $r$ = 0.46, p < 0.001 | $r$ = 0.0, p = 1.0 | $r$ = 0.0, p = 1.0 | $r$ = 0.0, p = 1.0 |
| Mag long | | $r$ = 0.0, p = 1.0 | $r$ = 0.0, p = 1.0 | $r$ = 0.0, p = 1.0 |
| Context | | | $r$ = 0.0, p = 1.0 | $r$ = 0.0, p = 1.0 |
| Cert short | | | | $r$ = -0.44, p < 0.001 |
| **Idealized Reaction Time Matrices** | | | | |
| Mag short | $r$ = -0.17, p = 0.2 | $r$ = 0.32, p = 0.02 | $r$ = 0.24, p = 0.08 | $r$ = -0.11, p = 0.4 |
| Mag long | | $r$ = -0.64, p < 0.001 | $r$ = 0.13, p = 0.4 | $r$ = 0.24, p = 0.07 |
| Context | | | $r$ = 0.0, p = 1.0 | $r$ = 0.0, p = 1.0 |
| Cert short | | | | $r$ = -0.44, p < 0.001 |

**Table S1.** Idealized behavioral matrix correlations and corresponding p values for Accuracy (top) and reaction times (bottom). Tab

| | ***Test short* behavioral pattern results** | | | |
|---|---|---|---|---|
| **Matrix** | $r$ RT | $\beta$ RT | $r$ ACC | $\beta$ ACC |
| Mag short | $t_{46}$ = 7.7, p < 0.001 | $t_{46}$ = 5.7, p < 0.001 | $t_{46}$ = 26.5, p < 0.001 | $t_{46}$ = 18.8, p < 0.001 |
| Mag long | $t_{46}$ = 0.4, p = 0.7 | $t_{46}$ = 1.5, p = 0.1, | $t_{46}$ = 14.9, p < 0.001 | $t_{46}$ = 2.0, p = 0.05 |
| Context | $t_{46}$ = 3.8, p < 0.001 | $t_{46}$ = 2.0, p = 0.05 | $t_{46}$ = -0.1, p = 0.9 | $t_{46}$ = -0.2, p = 0.9 |

| | | | | |
|---|---|---|---|---|
| Cert short | $t_{46}$ = 10.0, p < 0.001 | $t_{46}$ = 7.0, p < 0.001 | $t_{46}$ = -0.5, p = 0.6 | $t_{46}$ = -0.6, p = 0.6 |
| Cert long | $t_{46}$ = -3.9, p < 0.001 | | $t_{46}$ = 1.1, p = 0.3 | |

**Table S2**. Table shows t-tests against zero. Accuracy matrices for magnitude during test long and test short are correlated (Table S2), and thus competitive regressions are not reported in main text.

| | ***Test long* behavioral results** | | |
|---|---|---|---|
| **Behavioral Metric** | ME Schedule | ME Assembly | Schedule×Assembly |
| RTs: within | $F_{2,41}$= 0.2, p = 0.8 | $F_{1,41}$= 0.9, p = 0.3 | $F_{2,41}$= 0.5, p = 0.6 |
| RTs: between | $F_{2,41}$= 1.7, p = 0.2 | $F_{1,41}$= 0.3, p = 0.6 | $F_{2,41}$= 0.2, p = 0.8 |
| RT all | $F_{2,41}$= 0.9, p = 0.4 | $F_{1,41}$= 0.01, p = 0.9 | $F_{2,41}$= 0.2, p = 0.9 |
| ACC: within | $F_{2,41}$= 0.1, p = 0.9 | $F_{1,41}$= 14.4, p < 0.001 | $F_{2,41}$= 0.01, p > 0.9 |
| ACC: between | $F_{2,41}$= 4.0, p = 0.03 | $F_{1,41}$= 65.8, p < 0.001 | $F_{2,41}$= 0.06, p > 0.9 |
| ACC: all | $F_{2,41}$= 0.2, p=0.8 | $F_{1,41}$= 62.8, p < 0.001 | $F_{2,41}$= 0.05, p > 0.9 |

**Table S3.** 2-way ANOVA results with Schedule and Assembly as factors for accuracies and reaction times during *test long*. Results are shown over all comparisons, as well as comparisons between contexts and within contexts

| | ***Test long* behavioral pattern t-tests** | | | |
|---|---|---|---|---|
| **Matrix** | *r* RT | *β* RT | *r* ACC | *β* ACC |
| Mag Short | $t_{46}$ = 2.7, p < 0.01 | $t_{46}$ = 2.5, p = 0.02 | $t_{46}$ = 5.5, p < 0.001 | $t_{46}$ = 1.5, p = 0.3 |
| Mag Long | $t_{46}$ = 3.4, p = 0.001 | $t_{46}$ = 1.7, p = 0.09 | $t_{46}$ = 9.2, p < 0.001 | $t_{46}$ = 8.2, p < 0.001 |
| Context | $t_{46}$ = -0.9, p = 0.4 | | $t_{46}$ = 0.5, p = 0.6 | |
| Cert Short | $t_{46}$ = 3.3, p < 0.01 | $t_{46}$ = 5.0, p < 0.001 | $t_{46}$ = 0.9, p = 0.4 | $t_{46}$ = 1.2, p = 0.09 |
| Cert Long | $t_{46}$ = 4.1, p < 0.001 | $t_{46}$ = 5.9, p < 0.001 | $t_{46}$ = -0.09, p = 0.9 | $t_{46}$ = 0.3, p = 0.1 |

**Table S4.** Behavioral pattern t-test results over all participants during *test long*. Results are shown for idealized matrices independently through Pearson correlations, as well as jointly through competitive regressions.

| | ***Test long* behavioral pattern ANOVAs** | | |
|---|---|---|---|
| **Behavioral Metric** | ME Schedule | ME Assembly | Schedule×Assembly |
| *r* Mag Short | $F_{2,41}$= 1.7, p = 0.2 | $F_{1,41}$= 10.5, p = 0.002 | $F_{2,41}$= 0.7, p = 0.7 |
| *r* Mag Long | $F_{2,41}$= 3.2, p = 0.05 | $F_{1,41}$= 44.7, p < 0.001 | $F_{2,41}$= 4.2, p = 0.02 |
| *r* Context | $F_{2,41}$= 7.3, p = 0.002 | $F_{1,41}$= 24.2, p < 0.001 | $F_{2,41}$= 3.8, p = 0.03 |
| *r* Cert Short | $F_{2,41}$= 0.8, p = 0.5 | $F_{1,41}$= 0.4, p = 0.5 | $F_{2,41}$= 0.1, p = 0.9 |
| *r* Cert Long | $F_{2,41}$= 1.3, p = 0.3 | $F_{1,41}$= 13.5, p < 0.001 | $F_{2,41}$= 0.9, p = 0.4 |

| | | | |
|---|---|---|---|
| *β* Mag Short | $F_{2,41}$= 2.7, p = 0.08 | $F_{1,41}$= 3.7 , p = 0.06 | $F_{2,41}$= 0.2, p = 0.8 |
| *β* Mag Long | $F_{2,41}$= 1.7, p = 0.2 | $F_{1,41}$= 27.7, p<0.001 | $F_{2,41}$= 4.7, p = 0.01 |
| *β* Cert Short | $F_{2,41}$= 2.5, p = 0.09 | $F_{1,41}$= 0.05, p=0.8 | $F_{2,41}$= 0.9, p = 0.4 |
| *β* Cert Long | $F_{2,41}$= 1.4, p = 0.3 | $F_{1,41}$= 6.4, p=0.02 | $F_{2,41}$= 1.1, p = 0.4 |

**Table S5.** Behavioral pattern 2-way ANOVA results with Schedule and Assembly as factors during *test long*. Results are shown for independent estimates through Pearson correlations (top 5 rows) and for joint estimates using competitive regression (bottom 4 rows).

| Online Test Long | ME Schedule | ME Assembly | Schedule×Assembly |
|---|---|---|---|
| ACC Within Ctx | $F_{2,134}$ = 2.4, p = 0.1 | $F_{1,134}$ = 10.7, p = 0.001 | $F_{2,134}$ = 1.9, p = 0.2 |
| ACC Between Ctx | $F_{2,134}$ = 1.1, p = 0.3 | $F_{1,134}$ = 27.7, p < 0.001 | $F_{2,134}$ = 2.7, p = 0.07 |
| RTs Within Ctx | $F_{2,134}$ = 2.0, p = 0.1 | $F_{1,134}$ = 6.3, p = 0.01 | $F_{2,134}$ = 3.2, p = 0.04 |
| RTs Between Ctx | $F_{2,134}$ = 3.8, p = 0.03 | $F_{1,134}$ = 3.9, p < 0.05 | $F_{2,134}$ = 5.1, p = 0.007 |

**Table S6.** Online experiment 2-way ANOVA results with Schedule and Assembly as factors for accuracies and reaction times during *test long*. Results are shown over for comparisons between contexts and within contexts

| ***Test long* Context behavioral pattern regression results** | | | | |
|---|---|---|---|---|
| **Behavioral Metric** | *β* Mag Short | *β* Mag Long | *β* Context | *β* Cert Long |
| ACC | $t_{46}$ = 1.5, p = 0.2 | $t_{46}$ = 8.2, p < 0.001 | $t_{46}$ = 0.5, p = 0.6 | $t_{46}$ = -0.1, p = 0.9 |
| RT | $t_{46}$ = 4.6, p < 0.001 | $t_{46}$ = 2.6, p = 0.01 | $t_{46}$ = -0.3, p > 0.8 | $t_{46}$ = 3.4, p = 0.001 |
| ME Schedule | $F_{2,41}$= 1.3, p = 0.3 | $F_{2,41}$= 0.5 , p = 0.6 | $F_{2,41}$= 5.2, p < 0.01 | $F_{2,41}$= 2.0, p = 0.1 |
| ME Assembly | $F_{1,41}$= 1.1, p = 0.3 | $F_{1,41}$= 5.2, p=0.03 | $F_{1,41}$= 3.4, p = 0.07 | $F_{1,41}$= 7.7, p < 0.01 |
| Schedule × Assembly | $F_{2,41}$= 0.7, p = 0.5 | $F_{2,41}$= 2.1, p=0.1 | $F_{2,41}$= 2.5, p = 0.1 | $F_{2,41}$= 0.3, p = 0.8 |

**Table S7.** *Test long* behavioral pattern competitive regression including context as regressor. Top two rows show t-tests against zero for beta-estimates drawn from accuracy and RT matrices. Bottom three rows show reaction time matrix 2-way ANOVA results with Schedule and Assembly as factors.
